# HIRA-mediated loading of histone variant H3.3 controls androgen-induced transcription by regulation of AR/BRD4 complex assembly at enhancers

**DOI:** 10.1101/2023.05.08.536256

**Authors:** Viacheslav M. Morozov, Alberto Riva, Sadia Sarwar, WanJu Kim, Jianping Li, Lei Zhou, Jonathan D. Licht, Yehia Daaka, Alexander M. Ishov

## Abstract

Incorporation of histone variant H3.3 comprises active territories of chromatin. Exploring the function of H3.3 in prostate cancer (PC), we found that knockout (KO) of H3.3 chaperone HIRA suppresses PC growth *in vitro* and in xenograft settings, deregulates androgen-induced gene expression and alters androgen receptor (AR) binding within enhancers of target genes. H3.3 affects transcription in multiple ways, including activation of p300 by phosphorylated H3.3 at Ser-31 (H3.3S31Ph), which results in H3K27 acetylation (H3K27Ac) at enhancers. In turn, H3K27Ac recruits bromodomain protein BRD4 for enhancer-promoter interaction and transcription activation. We observed that HIRA KO reduces H3.3 incorporation, diminishes H3.3S31Ph and H3K27Ac, modifies recruitment of BRD4. These results suggest that H3.3-enriched enhancer chromatin serves as a platform for H3K27Ac-mediated BRD4 recruitment, which interacts with and retains AR at enhancers, resulting in transcription reprogramming. AR KO reduced levels of H3.3 at enhancers, indicating feedback mechanism. In addition, HIRA KO deregulates glucocorticoid-driven transcription, suggesting a common H3.3/HIRA-dependent mechanism of nuclear receptors function. Expression of HIRA complex proteins is increased in PC compared with normal prostate tissue, especially in high-risk PC groups, and is associated with a negative prognosis. Collectively, our results demonstrate function of HIRA-dependent H3.3 pathway in regulation of nuclear receptors activity.

**Key points:** *H3.3 at enhancers promotes acetylation of H3K27Ac and retention of AR/BRD4 complex for transcription regulation

*Knockout of H3.3 chaperone HIRA suppresses PC cells growth and deregulates androgen-induced transcription

*H3.3/HIRA pathway regulates both AR and GR, suggesting a common HIRA/H3.3 mechanism of nuclear receptors function

## Introduction

Prostate cancer (PC) is the second leading cause of cancer mortality in American men (1). PC that relapses after hormonal therapies (castration-resistant PC; CRPC (2)) is the cause of almost all PC-related deaths. Pathologic growth of the prostate is controlled mainly by steroid androgens. Recently diagnosed (both local and metastatic) diseases are treated with androgen ablation therapies that suppress androgen production or block androgen binding to AR ligand-binding domain (LBD) (3), thereby preventing nuclear translocation of AR and its binding to AR Response Elements (AREs). These therapies are effective in majority of patients, yet offer only a temporary relief, and the disease eventually recurs as treatment-resistant CRPC that is characterized by either AR loss-of function (treatment-emergent neuroendocrine PC, t-NEPC (4,5)) or gain-of-function, expressing AR variants (AR-Vs) with deleted LBD (AR-DLBD) (6). Identification of additional mechanisms involved in the transition of androgen-dependent PC to CRPC presents an opportunity to improve disease diagnosis and outcome. Hence, identification of co-regulators of AR may yield targets (and drugs) for effective and sustained management of CRPC (7).

Initiation and progression of PC is determined by, among other factors, transcription reprogramming that may be controlled by epigenetic dysregulation. The main chromatin unit, the nucleosome, can be modulated by post-translational modifications of histones, and by the incorporation of histone variants that together determine transcription activity of chromatin. There are three variants of histone H3: H3.1, H3.2, and H3.3 (8). H3.1 and H3.2 are the predominant forms (9,10) and H3.3 differs from H3.1 by only five amino acids. While canonical H3.1/2 are incorporated into chromatin in S-phase, H3.3 is deposited through interphase, and is referred to as “replication-independent histone” (11,12). Histone variants are deposited by specific chaperones. H3.3 is chaperoned by HUCA (<u>H</u>IRA, <u>U</u>BN1, <u>C</u>ABIN1, <u>A</u>SF1a) (13) and Daxx/ATRX complexes (14,15), which deposit H3.3 at enhancers, transcription start sites (TSS), centromeres and telomeres (14–17). Chromatin enriched in H3.3 is associated with elevated transcription activity (18). H3.3-containing nucleosomes are less stable compared to nucleosomes with canonical histones, thereby providing DNA accessibility to the transcription machinery at regulatory elements (19). These observations prompted Hake and Allis to propose the “histone variant barcoding” hypothesis, stating that “histone H3 variants exhibit distinct posttranslational ‘‘signatures’’ that influence epigenetic states during differentiation and development”(9). Concordant with this model, H3.3 deposition coincides with H3K4me3, a mark for transcriptionally active promoters (9,14,20,21); H3.3-enriched chromatin is low with silencing marks (22,23). Epigenetic regulation by transcription-associated histone variant H3.3 was implicated in cancer by the discovery of essential function of H3.3 deposition pathways in several malignancies, suggesting oncogenic function. Multiple lines of evidence pointed to the essential function of H3.3 deposition pathways in the initiation and progression of pancreatic, brain and breast cancers (24),(25),(26),(27), implying an oncogenic function of H3.3 pathway.

H3.3 is enriched at enhancers (28), and recent findings implicated H3.3 in the activation of histone acetyltransferase p300 resulting in acetylation of H3K27 (H3K27Ac) at enhancers (29). This function required phosphorylation of H3.3S31 (H3.3S31Ph), a unique H3.3 residue, by checkpoint kinase CHK1(29),(30) and by the main activator of the inflammatory transcription factor NF-κB, kinase IKKα (31). AR binds to AREs in majority of active enhancers and super-enhancers (SE) (32) in distal intergenic or intronic regions of target genes. Together, these data suggest a function of H3.3 in the AR-driven transcription in PC. Investigating function of H3.3 pathway in PC, we observed that HIRA KO strikingly decreases androgen induced transcription. Evaluating enhancers associated with androgen-target genes, we found that HIRA KO reduces H3.3 incorporation, diminishes H3K27Ac and H3.3S31Ph. In addition, it alters dynamics of BRD4 and AR binding at enhancers, first elevating and then reducing levels of these proteins at AREs. These data suggest that H3.3-enriched enhancer chromatin serves as a platform for AR-dependent and H3K27Ac-mediated recruitment of BRD4 to enhancers, implying two-steps in the assembly of transcription complex for AR-driven transcription reprogramming. In addition, HIRA KO deregulates glucocorticoid-driven transcription, suggesting a common H3.3/HIRA-dependent mechanism of nuclear receptors function. Collectively, our data suggest a function of HIRA-dependent H3.3 pathway in PC progression.

## Methods

### Cell culture

Isogenic R1-AD1 (AR-WT) and R1-D567 (ARDLBD) cells (33) (received from Dr. Scott Dehm, University of Minnesota, Minneapolis, MN) were cultured in RPMI 1640 medium (Corning, # 10040CV) supplemented with 10% fetal bovine serum (Gibco, #10437036), and 100 U/mL penicillin and 100 µg/mL streptomycin (Gibco BRL, Carlsbad, CA) and grown at 37 °C in a humidified 5% CO2 incubator. 72 h before experiment cells were transferred to RPMI 1640 phenol-free media (Gibco # 11835030), supplemented with 10% charcoal stripped fetal bovine serum (CSS, Gibco, # 12676029), and 100 U/mL penicillin and 100 µg/mL streptomycin. Synthetic androgen R1881 (Sigma-Aldrich, # R0908) was prepared according to manufacturer recommendations and used in 1 nM concentration. JQ1 (Sigma-Aldrich, # SML1524) was prepared according to manufacturer recommendations and used in concentrations as outlined in specific experiments.

### Antibodies

The following antibodies were used in this study: HA mouse monoclonal antibody (16B12) (BioLegend, # 901502), FLAG M2 mouse monoclonal antibody (Sigma, # F1804), AR rabbit polyclonal antibody (Millipore, # 06-680) for ChIP, AR N-20 rabbit polyclonal antibody (Santa Cruz, # sc-816) for Western blotting, FKBP5 rabbit polyclonal antibody (Cell Signaling, #8245), H3K4Me1 rabbit polyclonal antibody (EpiCypher, # 13-0040), H3K27Ac mouse monoclonal antibody (EpiCypher, # 13-0045), HIRA(34), Daxx rabbit polyclonal antibody (35); BRD4 rabbit polyclonal antibody (Fortis Life Science, # A301-985A100) for Western blotting and mab BL-149-2H5 for ChIP, GR (G-5) mouse monoclonal antibody (Santa Cruz, # sc-393232), H3.3S31Ph rabbit monoclonal antibody (Abcam, # ab92628), Actin (AC-74) mouse monoclonal antibody (Sigma, # A5316).

### Production of FLAG-HA tagged H3F3A cells

Two oligos H3F3AN-1-73F CACCGTCAATGCTGGTAGGTAAGTA and H3F3AN-1-73R AAACTACTTACCTACCAGCATTGAC containing gRNA sequence were annealed and inserted into pSpCas9-2A-Puro vector (PX459, Addgene # 62988), digested with BbsI-HF enzyme (NEB, # R3539S). DNA sequence, 1 KB upstream and downstream from the start codon of the H3F3A gene, was cloned into pUC19 backbone, then sequence coding FLAG-HA tag was introduced into 5’-end of H3.3 ORF by Gibson assembly. Two nucleotide’s substitutions were made in sequence recognized by gRNA to prevent repeated targeting of the modified allele. R1-AD1 and R1-D567 cells were transfected with equimolar amount of pSpCas9-2A-Puro H3F3A-1-73 plasmid and pUC19-FLAG-HA-H3F3A plasmid using Lipofectamine 2000 (Invitrogen). After 3 days transfected cells were trypsinized, counted and placed into 96-well plates, ∼1 cell/well, for clonal growth. Three weeks later each 96-well plate was duplicated, and clones were analyzed for insertion of FLAG-HA tag by Restriction Fragment Length Polymorphism and immunofluorescence staining. Correct insertion of FLAG-HA tag was confirmed by sequencing (MGH CCIB DNA Core, Cambridge).

### Production of AR, HIRA and Daxx knockout (KO) cell lines

The Alt-R CRISPR-Cas9 System (Integrated DNA Technologies) was used to produce KO cells. The following protospacer sequences were use in crRNA design for knockout of AR, HIRA and Daxx genes: CTGGGACGCAACCTCTCTCG, CTGGCTAGCCTCATGCAGCG and GCAGACAGCAGACCACCCTG accordingly. crRNA was annealed with tracrRNA and mixed with Cas9 enzyme (Integrated DNA Technologies)). R1-AD1-FLAG/HA-H3F3A cells were electroporated using Neon Transfection system (Invitrogen). After 3-5 days cells were trypsinized, counted and placed into 96-well plates, ∼1 cell/well, for clonal growth. Knock-out of target genes was confirmed immunofluorescence, Western blotting and by DNA sequencing.

### RNA-seq

1×10^6^ cells were plated on 6 cm dish in RPMI 1640 medium, sans phenol-red (Gibco, #11835-030), 10% CSS (Invitrogen), 1% penicillin / streptomycin solution (Corning, # 30-002-CI) and grown for 72 h. Cells were treated with 1 nM R1881 for 4 h, 12 h and 24 h. RNA was isolated using RNeasy Plus Kit (QIAGENE, # 74134) according to the manual and sequenced at Novogene.

### ChIP-seq

ChIP-seq protocol (36) with some modification was used. 10×10^6^ cells were fixed with 1% formaldehyde in PBS for 10 min at room temperature, quenched with 1.25 M glycine to a final concentration of 0.125 M for 5 min. Cells were washed twice with ice-cold PBS, scraped into 6 ml Farnham lysis buffer (5 mM PIPES pH 8.0, 85 mM KCl, 0.5% Igepal CA-630) with protease inhibitors (ThermoFisher, # A32953), transferred to 15 ml Falcon tubes, and centrifuged at 800 rcf for 5 min at 4 L. The pellet was resuspended in 1 ml fresh Farnham lysis buffer and incubated on ice for 10 min. Nuclei were collected by centrifugation at 500 rcf, 4 L for 5 min, resuspended in 150 µl cold RIPA buffer (1XPBS (Sigma, # D8662), 1% Igepal CA-630, 0.5% sodium deoxycholate), 0.5% SDS, and transferred to TPX tube (Diagenode). Chromatin was sheared by Diagenode Bioraptor Pico with peak enrichment at 200 bp. After sonication SDS concentration was adjusted to 0.1% SDS with RIPA buffer, and chromatin was spin at maximum speed in a microfuge for 15 min at 4 _JC. The supernatant was transferred into new tube and snap frozen in liquid nitrogen until needed. Chromatin was pre-cleared with Pierce Protein A/G Magnetics beads (Pierce # 88802). 2-5 µg primary antibody were added per 5×10^6^ cells and incubated overnight at 4 L with rotation. Next day 25 µl Pierce Protein A/G Magnetic beads (pre-blocked with PBS/BSA), were added, and incubated for 2 h at 4 L with rotation. Beads were washed 5 times with 1 ml wash buffer (100 mM Tris-HCl, pH 7.5, 500 mM LiCl, 1% Igepal CA-630, 1% sodium deoxycholate) and 1 time with 1 ml TE buffer (10mM Tris-HCl pH 7.5, 0.1 mM EDTA), with 3 min rotation in-between for each wash. Chromatin was eluted from beads with 150 μl fresh prepared IP elution buffer (1% SDS, 0.1 M NaHCO_3_) and de-crosslinked overnight at 65 L. DNA was purified using Monarch PCR/DNA Cleanup Kit (New England BioLabs). ChIP-seq libraries were prepared using NEBNext Ultra II DNA Library Prep Kit for Illumina (New England BioLabs) and sequenced on the Illumina NovaSeq 6000 Sequencer (2×150) at the University of Florida ICBR.

### ATAC-seq

ATAC-seq libraries were prepared using ATAC-seq kit from Active Motif (Active Motif, CA, # 53150) according to the manual and sequenced on the Illumina NovaSeq 6000 Sequencer (2×150) at the University of Florida ICBR.

### Bioinformatic analysis

#### RNA-Seq

Short reads were mapped to the GRCh38 reference transcriptome using STAR (37). Quantification produced tables of FPKM values for each gene in each sample. Differential analysis was performed with DESeq2 (38), generating tables of significantly over- or under-expressed genes in each contrast.

#### ChIP-Seq and ATAC-Seq

Short reads were mapped to the GRCh38 reference genome using Bowtie2 (39). Peak calling was performed with MACS (40) using parameters appropriate for the type of signal being detected. Super-enhancers were identified using ROSE version 0.1 (41). All interval operations were performed with bedtools (https://bedtools.readthedocs.io/en/latest/), and plots were generated with custom scripts. Differential analysis of ChIP-Seq and ATAC-Seq peaks was performed with DASA (https://github.com/uf-icbr-bioinformatics/dasa).

#### Profile plots and boxplots

To generate the profile plot for a given signal in a set of regions, we extracted the normalized coverage values from the short-read alignment file for that signal for each region, and we computed the geometric average of the results to reduce the impact of outliers. Profiles for the different cell lines or timepoints were then superimposed in the same chart. A similar process was used to generate metagene profile plots, in which all gene regions were scaled to the same length, and a fixed flanking region of 1Kb was added upstream and downstream. Boxplots were generated based on the total coverage in each region. All processing was performed using custom scripts.

### Statistical analysis

Statistical analysis was performed with GraphPad Prism 9.2.0 (GraphPad Software, Inc., San Diego, CA). Homogeneity of variance was estimated using Brown-Forsythe test. The comparison of means between different groups was performed by one-way ANOVA with either Tukey or Dunnett multiple comparison test correction.

### Western blotting analysis

Protein samples were separated on 4–20% Mini-Protean TGX gel (Bio-Rad, #4561096) and transferred to nitrocellulose membrane using iBlot 2 system (Invitrogen, Thermo Fisher Scientific). Membranes were blocked with 5% nonfat milk/PBS, 0.1% Tween (PBST). Primary antibodies were diluted in 5% milk/PBST and incubated overnight at 4 °C. Membranes were washed two times with PBST and incubated for 1 h at room temperature with appropriate IRDye secondary antibody (Li-COR Biosciences). Membranes were washed three times with PBST and visualized by Odyssey CLx Imaging System (Li-COR Biosciences).

### Colony formation assay

5×10^3^ cells were seeded in 12-well plates in complete media, cultured for 7 days, fixed for 10 min with 4% formaldehyde and stained with crystal violet (0.5%). Images were acquired with Epson photo scanner and area of colonies was calculated using ImageJ software. Experiments were repeated at least three times.

### Proliferation assay

Cells (5000 cell/well, 96-well plates) were set in 100 μl of RPMI 1640 phenol-free media supplemented with 10% CSS; 10 μL Alamar® blue reagent (Thermo Scientific # 00-100) was added to each well. Data were collected using Spectra Max M3 plate reader after 4 h of signal development.

### 3D prostaspheres and treatment

The prostaspheres were grown as described (42) in Matrigel (Corning) in DMEM supplemented with 10% fetal bovine serum in 24-well plates. The basal layer was formed by mixing Matrigel and medium at a ratio 1:1; 250 μl of mixture were added per well. 3000 R1-AD1 cells (parental, AR KO, Daxx KO, HIRA KO) were suspended in media and mixed with Matrigel at a ratio of 10:1 in 200 μl of mixture and this suspension was placed on the top of pre-solidified base layer. The plate was placed in CO2 tissue culture incubator to allow the upper layer to solidify. Next, 1 ml of complete media was added to each well. Prostaspheres were treated with 200 nM JQ1 on day 11 and day 14 and documented at day 21 with inverted Leica fluorescent microscope DMI4000B.

### Xenograft experiments

Soft collagen pellets containing parental R1-AD1, AR KO, Daxx KO or HIRA KO cells were implanted subcutaneously to establish xenografts in 6 weeks old male Hsd:Athymic Nude-Foxn1nu mice. After 4 weeks, tumor growth was analyzed as we described before (43).

### Immunofluorescence

For characterization of H3.3 in cells, immunofluorescence was done as described (44). Briefly, 75 × 10^4^ cells were plated on microscope coverslip glass (Fisher Scientific) in RPMI1640/ 10% FBS media. Сells were fixed with 1% formaldehyde for 10 min, permeabilized with 0.5% Triton X-100, and incubated with HA antibodies for 1 h at room temperature. After two washes with PBS, cells were incubated with secondary antibodies conjugated with Alexa Fluor 488 or 594 dye (Invitrogen). DNA was stained with Hoechst 33342 (Sigma). Images were analyzed using either Leica DMI4000 B fluorescent microscope or Leica TCS SP5 confocal microscope.

## Results

### Characterization of CRPC cell models

H3.3 and H3.1/2 differ by only 5 amino acids, and 4 of these are in the globular part of histone (and, therefore, are not exposed in the context of nucleosome/chromatin) thus making it difficult to produce H3.3-specific ChIP-grade antibody. To overcome this challenge, we used CRISPR-Cas9 genome editing to knock-in FLAG/HA-epitope tags at 5’ of H3F3A gene, thereby producing R1-AD1 (AR-WT) cells (33) expressing endogenous FLAG/HA-H3.3. Cells were characterized by sequencing, immunofluorescence (IF), and Western blot (**Fig. S1A, B**). Using CRISPR/Cas9 genome editing on FLAG/HA-H3.3 R1-AD1 cells, we produced AR, HIRA and Daxx knockout (KO) cells. At least three subclones were characterized for each KO by sequencing, IF, and Western blot (**Fig. S1B**). Deletion of AR, HIRA or Daxx did not affect levels of remaining two proteins. HIRA-KO reduced H3.3 levels, potentially by a negative compensatory mechanism. Thus, we created and characterized unique cell models to selectively test function of endogenous H3.3 and its pathway in PC.

### HIRA affects PC cell proliferation

Using KO cells, we tested effect of manipulating H3.3 chaperones levels on cell proliferation. KO of AR or HIRA reduced the R1-AD1 cell proliferation rate and colony formation (**Fig. 1A, B**). Daxx KO had a reproducible yet lower effect on cell growth. These data establish HIRA function in proliferation of PC cells.

**Fig. 1.**
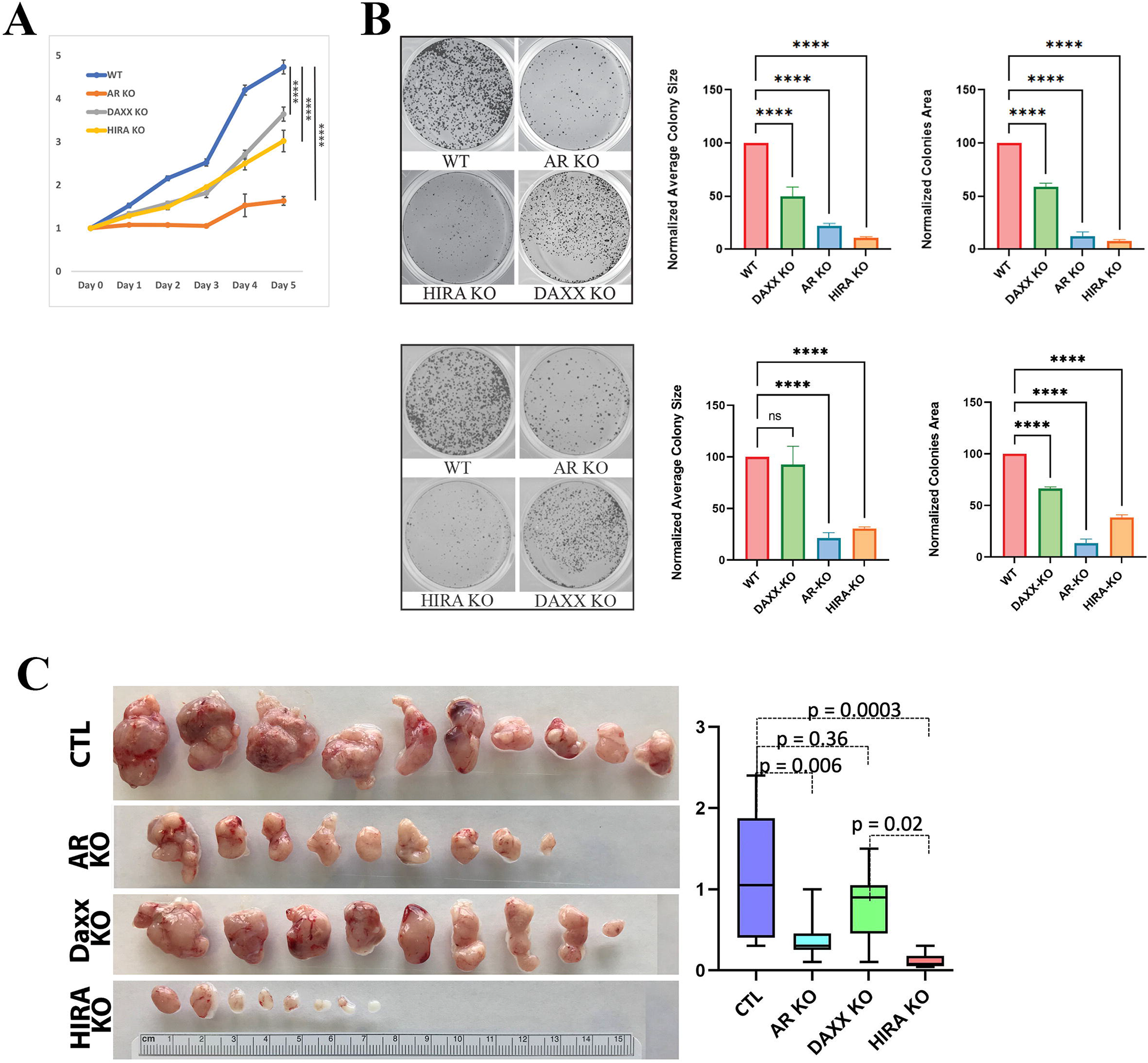
Modification of HIRA affects proliferation of PC cells. **A**: Proliferation of R1-AD1 and Daxx KO, AR KO, HIRA KO cell lines. **B**: colony formation assays with these cell lines in androgen deprived (top) and R1881 stimulated conditions (bottom). Left: representative images; middle: average colony size; right: combined colonies area calculated with ImageJ. P-values: **** < 0.0001. **C**: Modification of HIRA affects xenograft growth of PC cells. 10^6^ R1-AD1 cells (CTL), Daxx KO, AR KO and HIRA KO cells were implanted subcutaneously in athymic nude mice, 10 per group. 4 weeks later, tumors were collected and photographed (left). Tumor weight in g (right).

To further address function of HIRA in PC, effect of manipulated HIRA expression was analyzed *in vivo*. Soft collagen pellets containing parental R1-AD1, and cells with KO of AR, Daxx, or HIRA were implanted subcutaneously to establish xenografts in 6 weeks old male Hsd:Athymic Nude-Foxn1nu mice. After 4 weeks, tumor growth was analyzed as we described before(43). Both AR and HIRA KO significantly reduced the tumor size (**Fig. 1C**). Daxx KO reproducibly decreased the tumor size, albeit to a lesser degree.

### HIRA affects androgen-induced gene transcription

To address mechanisms of HIRA-dependent effect on cell proliferation, we evaluated potential influence of HIRA on androgen-dependent transcription activity using RNA-seq analysis. RNA was isolated from R1-AD1 cells (parental cells) and individual AR, HIRA and Daxx KO clones under androgen-deprived (72h) and androgen (R1881) stimulated (4h, 12h, 24h) conditions. Genes with p-value < 0.05 (adjusted using the Benjamini and Hochberg’s approach for controlling the False Discovery Rate (FDR)) found by DESeq2 were considered as differentially expressed. Expression analysis is presented in **Fig. 2**; numbers correspond to genes that are at least two-fold (p-adjusted < 0.05) up- or down-regulated following R1881 stimulation in comparison to androgen-deprived parental cells. Using these parameters, we found that 409 genes were up- and 328 down-regulated at 4h in parental cells. AR KO abolished this regulation (with only 4 up- and 4 downregulated genes remaining), confirming AR-dependence. Both Daxx and HIRA KO affected androgen-stimulated gene expression, although HIRA KO impacted androgen-dependent gene expression more strongly than Daxx KO, as evident by reduction of upregulated genes to 49 and downregulated genes to 27 (**Figs. 2**; **S2A** for Venn diagram and **Table S1** for list of genes with androgen-regulated expression affected by both AR and HIRA KO). Using the KEGG database, we identified several androgen-regulated pathways that are affected by both AR and HIRA KO (**Fig. S2B and Table S2**), including cancer- and metastasis-associated pathways, as FoxO(45), TNF(46), TGF-beta(47), PI3K-Akt(45), MAPK(48) and Wnt(49). Similar results were observed at 12h and 24h of treatment (**Fig. 2**). We conclude that HIRA is required for androgen-regulated transcription.

**Fig. 2.**
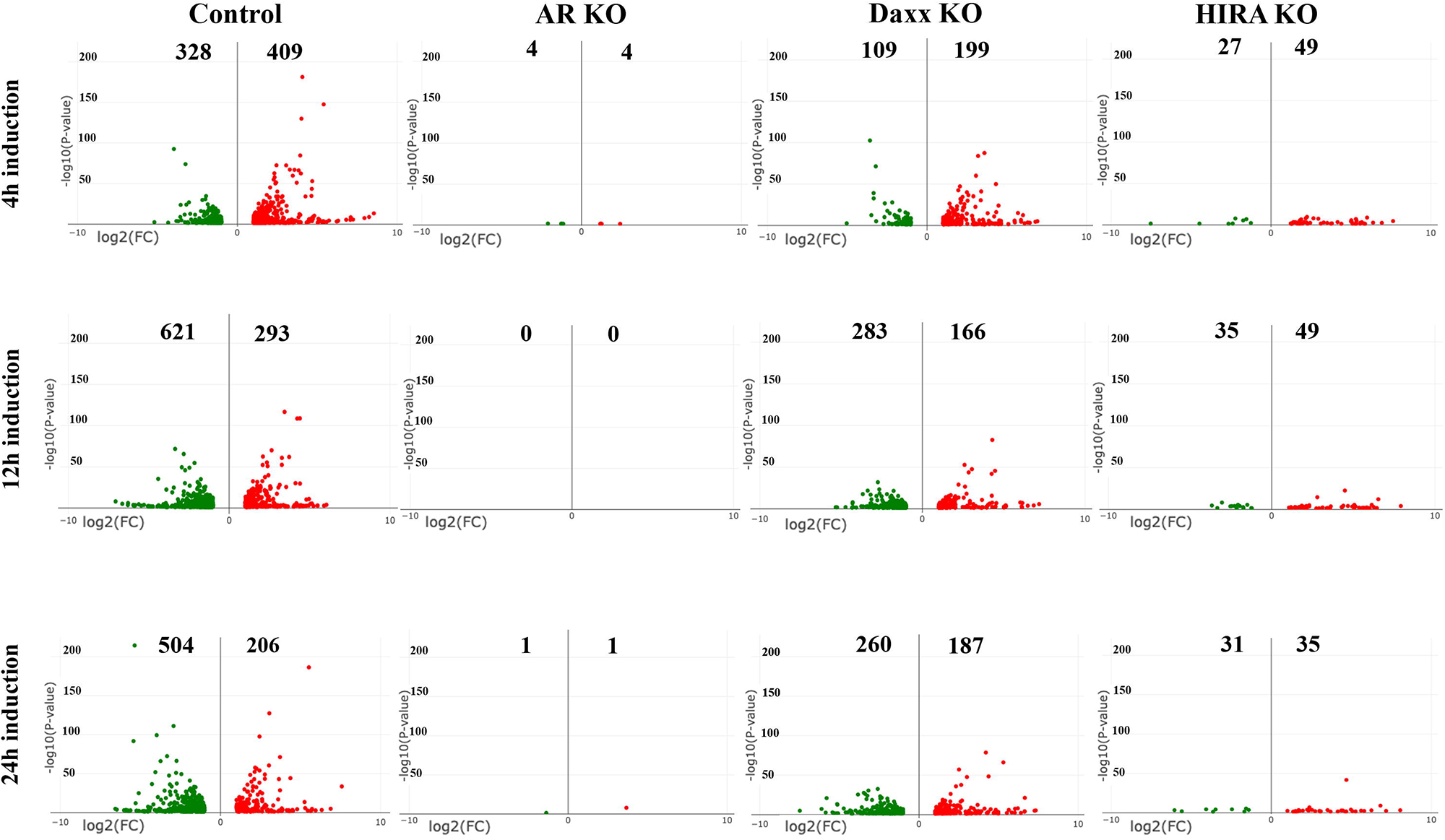
Differential gene expression after androgen stimulation. Expression analysis (RNA-seq, volcano plot) of R1-AD1 cells, parental (Control), AR KO, Daxx KO and HIRA KO. Numbers correspond to genes that are at least two-folds (p-adjusted <0.05) up- (red) or downregulated (green) by 4h (top row), 12h (middle row) or 24h (bottom row) of R1881 stimulation compared with androgen-deprived (72h) in parental cells. X: expression, log2; Y: P-value, log10. See **Fig. S2** for Venn diagrams and regulated pathways analysis.

### HIRA KO reduces levels of H3.3 and H3K27Ac at up- and downregulated genes

To address mechanism of transcription deregulation in KO cells, we performed analysis of H3.3 (endogenous HA-H3.3) and transcription-associated marker H3K27Ac at the transcription start sites (TSSs), gene bodies (GB), and transcription end sites (TES) focusing on 409 up- and 328 down-regulated genes (0h vs 4h R1881 stimulation, parental cells, **Fig. 2**). We compared ChIP-seq profiles in R1-AD1 parental (Control), AR, Daxx and HIRA KO cells under androgen-deprived (0h) and androgen-induced conditions (at 4h, 12h, 24h; levels of H3.3 were monitored by Western blot analysis, **Fig. S1C**). Compared with parental cells, AR KO reduced H3.3 levels at all three elements (TSS, GB, TES) at up-regulated genes during androgen induction, and at down-regulated genes in androgen-deprived conditions (**Figs. 3, S3** for TSS; in this and follow-up ChIP-seq analyses comparison was made between cell lines at the same time points and between time points within each cell line and **Fig. S4** for entire genes). Reduction of H3.3 in AR KO cells was concordant with a transcription-associated H3.3 deposition mechanism (50–52) that is abrogated at androgen-regulated genes in these cells. HIRA KO reduced H3.3 levels in up- and down-regulated genes at all three elements (TSS, GB, TES) at all time points, confirming function of this chaperone in H3.3 loading at coding regions (14,53). Accumulation of H3.3 in Daxx KO cells at 24h can be explained by the increased availability of H3.3 to HIRA or other H3 chaperones such as CAF-1(16) and NASP (54) for incorporation into gene coding element and is concordant with prior observations that Daxx KO reduces H3.3 at telomeres (15), centromeres and pericentromeres (55). In parental cells, accumulation of H3K27Ac was observed at TSS, gene bodies, and TES in up- and down-regulated genes at 12h and 24h (**Figs. 4, S5** for TSS and **Fig. S6**). We observed temporary elevation of H3K27Ac in Daxx KO cells (at 4h) that may be at least partly explained by a reduction of HDAC1/2 recruitment to chromatin, which is mediated by interaction with Daxx in parental cells (56,57). AR KO completely abolished, and HIRA KO strongly reduced H3K27Ac accumulation, in agreement with a recent publication (53).

**Fig. 3.**
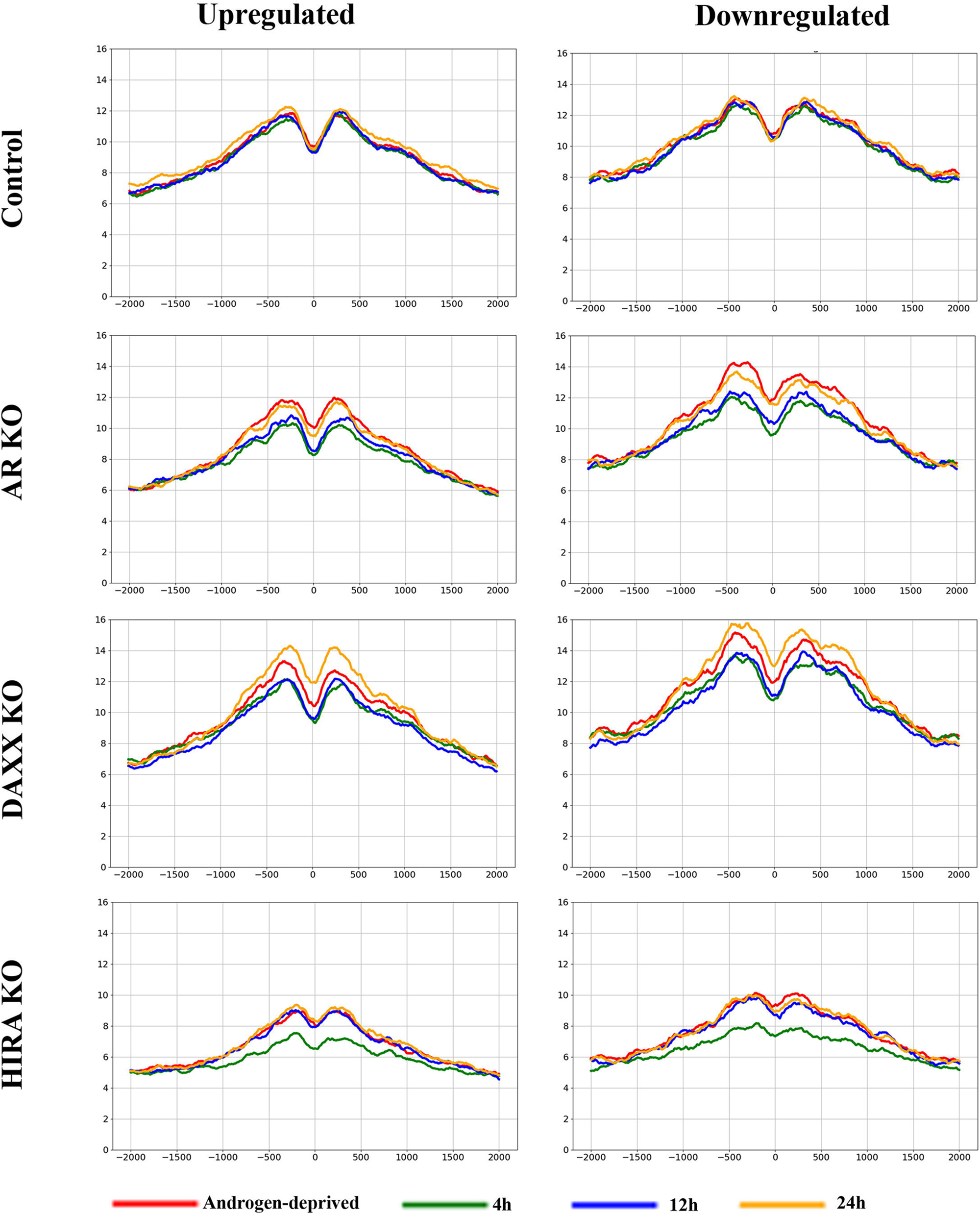
H3.3 association with TSS of androgen-regulated genes; analysis within cell lines. ChIP-seq profiles of H3.3 (endogenous HA-H3.3) at the 409 androgen-up-(**left**) and 328 androgen-downregulated (**right**) genes in R1-AD1 parental (Control), AR KO, Daxx KO and HIRA KO cells in the androgen-deprived (0h) and androgen-induced conditions (at 4h, 12h, 24h). 0: TSS. See **Fig. S3** for time points comparison.

**Fig. 4.**
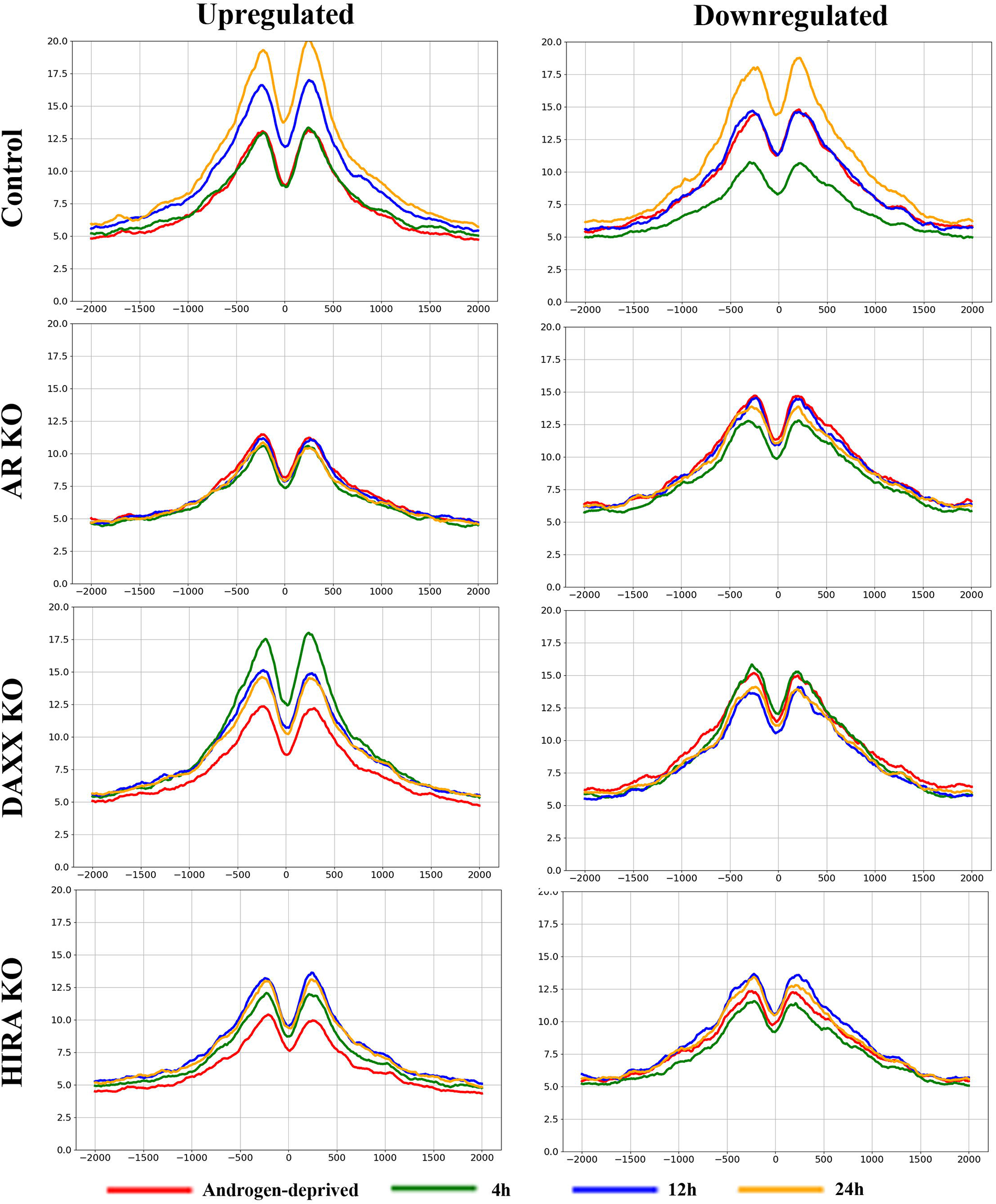
H3K27Ac association with TSS of androgen-regulated genes; analysis within cell lines. ChIP-seq profiles of H3K27Ac at the 409 androgen-up-(**left**) and 328 androgen-downregulated (**right**) genes in R1-AD1 parental (Control), AR KO, Daxx KO and HIRA KO cells in the androgen-deprived (0h) and androgen-induced conditions (at 4h, 12h, 24h). 0: TSS. See **Fig. S5** for time points comparison.

### HIRA KO affects AR association with chromatin

Massive deregulation of androgen-induced expression in HIRA KO cells (**Figs. 2, S2**) suggested altered activity of AR. Hence, we analyzed AR association with chromatin by ChIP-seq in R1-AD1 parental and KO cells (AR KO cells were used as a negative control) under androgen deprived and R1881 stimulated conditions (4h, 12h, 24h). At 4h after stimulation, we identified 5920, 5528, and 5412 AR peaks, corresponding to parental, Daxx and HIRA KO cells. In parental cells, AR association with chromatin was elevated at 4h and remained mostly unchanged at 12h and 24h (**Figs. 5** **left** and **S7A left**). We observed increased AR accumulation in Daxx KO cells at 4h of stimulation that may be explained by the reported contribution of Daxx/AR interaction to the negative regulation of AR binding to DNA (58), which is abrogated in Daxx deleted cells. In HIRA KO cells, AR association is elevated at 4h of stimulation, and is substantially reduced to almost pre-induced levels at 12h and 24h.

**Fig. 5.**
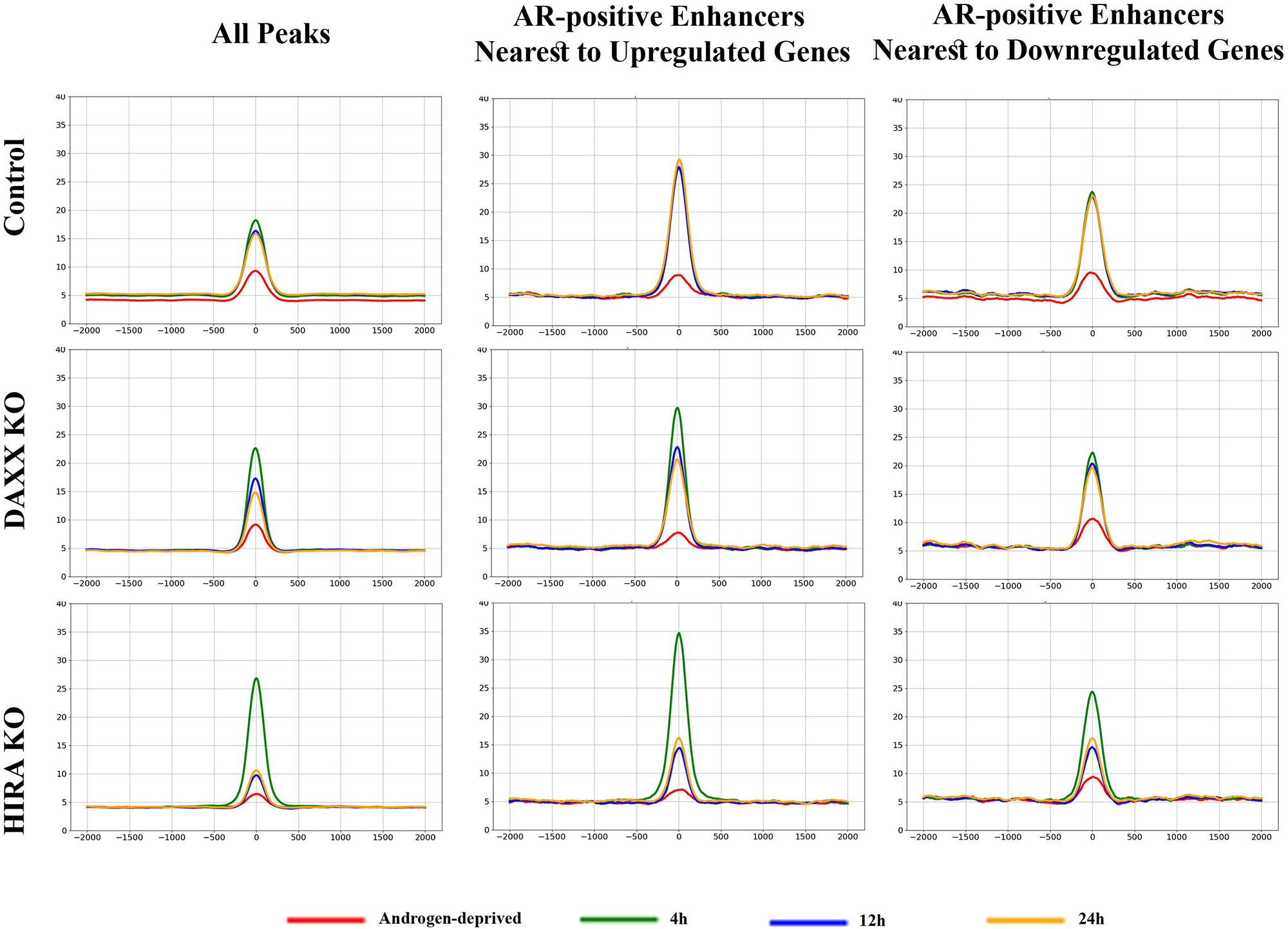
AR association with chromatin is regulated by HIRA. **Left:** Metaplot of AR peaks (position “0” at 4h in R1-AD1 parental cells) in R1-AD1 parental, HIRA KO and Daxx KO in androgen-deprived (72h, red) and R1881 stimulated for 4h (green), 12h (blue), 24h (yellow). In parental cells, AR association with chromatin is elevated at 4h of stimulation and remains at the same levels at later time points. In HIRA KO, association is reduced in deprived condition, is much higher compared with parental at 4h, and next is substantially reduced at 12h and 24h. Analysis of AR at enhancers nearest to the 409 up-(**middle**) and 328 downregulated (**right**) genes is similar to the overall AR behavior. See **Fig. S7** for time points comparison.

AR binds to active enhancers (32). Therefore, we analyzed dynamics of AR at enhancers nearest to the same group of up-(409) and down-regulated (328) genes. To map enhancers, we performed ChIP-seq for enhancer marker H3K4me1 (59) (**Fig. S8**). We identified AR-positive enhancers (by AR ChIP-seq at 4h in parental cells) nearest to each up- or down-regulated (at 4h) gene in parental R1-AD1 cells. We observed that AR KO reduced levels of H3K4me1 at AR-positive enhancers while HIRA KO had a similar effect on both AR-positive and AR-negative elements (most obvious at those associated with up-regulated genes, **Fig. S8**), implying overlapping functions of AR and HIRA/ H3.3 pathway in maintenance of lineage specific enhancers. Methyltransferases KMT2C (MLL3) and KMT2D (MLL4) monomethylate H3K4 (60), and a reported association between H3.3 and KMT2D identified by BioID (61), suggests recruitment of this methyltransferase to H3.3-enriched chromatin areas (as enhancers) for monomethylation of H3K4 and explains the reduction of H3K4me1 in HIRA KO cells.

Dynamics of AR at enhancers were similar to the AR behavior at all peaks (**Fig. 5** **middle**: for enhancers associated with up- and **Fig. 5** **right**: down-regulated genes; **S7** for time points comparison), with the highest AR amplitude at enhancers associated with up-regulated genes. HIRA KO elevated AR association with enhancers at 4h, and that was substantially reduced at 12h and 24h. Thus, HIRA KO affected dynamics of AR chromatin binding, including enhancers associated with up- and down-regulated genes. We conclude that HIRA-dependent pathway is important for the proper retention of AR at AREs after androgen induction, potentially explaining changes in transcription deregulation observed in HIRA KO cells.

### HIRA KO affects epigenetics at AR-positive enhancers associated with AR-regulated genes

H3.3 is enriched at enhancers (28), and we analyzed its association (endogenous HA-H3.3) with AR-positive enhancers nearest to genes up- and down-regulated by androgen in parental R1-AD1 cells (**Figs. 6** **and S9** for time points comparison). In androgen deprived conditions, AR binding site (“0” at the graphs) is occupied by H3.3, suggesting H3.3-containing nucleosome accumulation at AREs in the absence of AR. H3.3 levels are reduced during androgen induction at these sites, most visible in the upregulated group, indicating nucleosome displacement by AR. Levels of H3.3 at AR-positive enhancers associated with upregulated genes are elevated at 24h of induction (in parental and in Daxx KO cells), confirming transcription-associated loading of H3.3 (50–52). In AR KO cells, H3.3 is reduced at enhancers, consistent with H3.3 profiling at regulated genes (**Figs. 3, S3, S4**), suggesting role of AR-regulated enhancer transcription in maintenance of H3.3 at enhancers. We observed a major reduction of H3.3 in HIRA KO, confirming chaperone function of HIRA at enhancers.

**Fig. 6.**
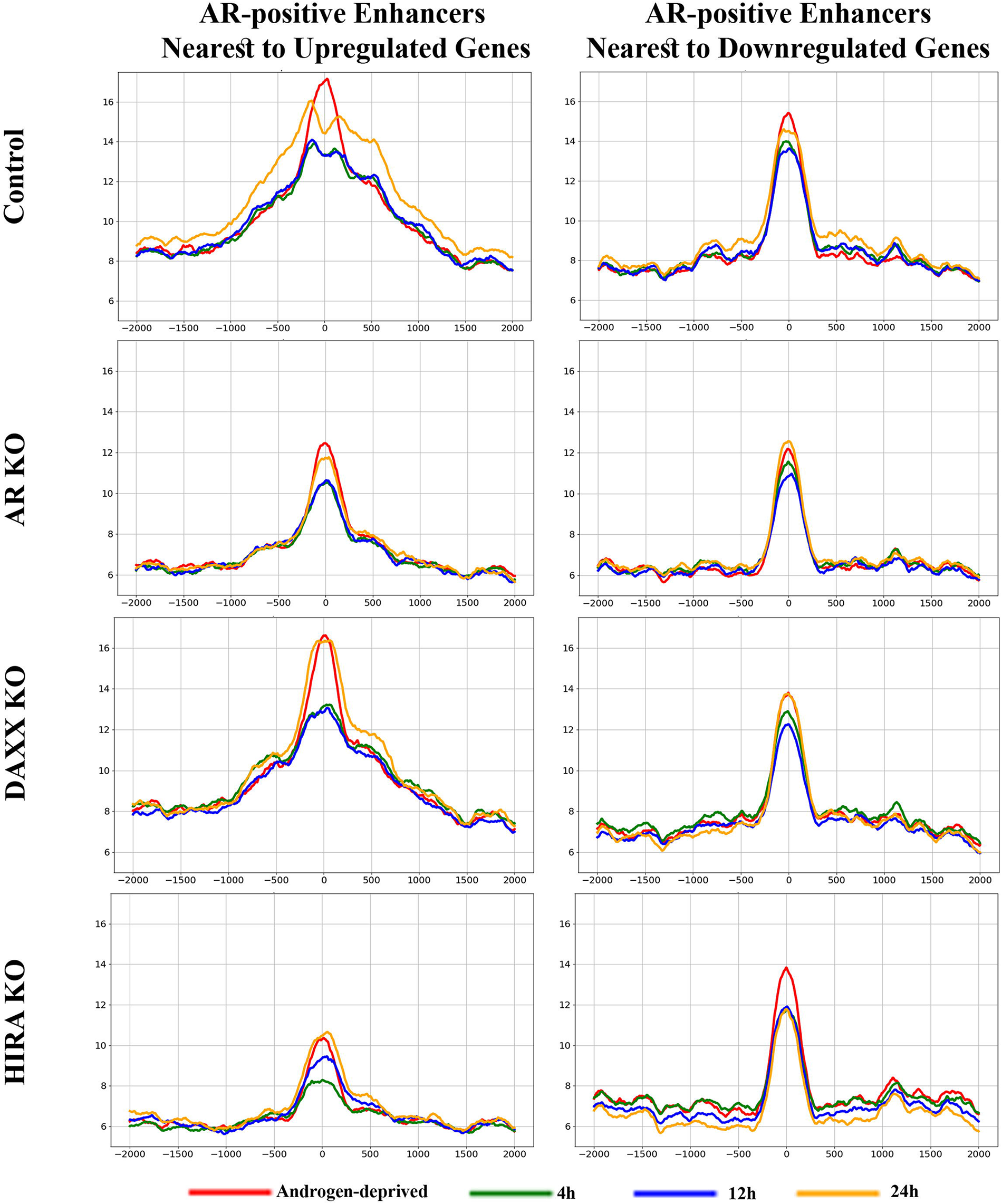
Dynamics of H3.3 at enhancers. Metaplot of H3.3 (endogenous HA-H3.3) at AR peaks (position “0”: AR at 4h in R1-AD1 parental cells) at enhancers associated with up-(left) and down-regulated (right) genes in R1-AD1 parental (Control), AR KO, Daxx KO and HIRA KO in androgen-deprived (72h, red) and R1881 stimulated for 4h (green), 12h (blue), 24h (yellow). Levels of H3.3 at AR-positive enhancers associated with upregulated genes are elevated at 24h of induction in parental and Daxx KO cells. H3.3 is reduced at enhancers in AR KO cells, suggesting AR function in maintenance of this transcription associated histone variant, and HIRA KO, confirming HIRA chaperone function. See **Fig. S9** for time points comparison.

Next, we analyzed the marker of active enhancers H3K27Ac (59) at AR-positive enhancers nearest genes regulated by androgen. In parental and Daxx KO cells, H3K27Ac was gradually elevated following androgen treatment at enhancers associated with up-regulated genes (**Figs. 7** **and S10** for time points comparison). Similar to H3.3, H3K27Ac was reduced at AR binding sites (“0” at the graphs) during androgen induction, most clear at enhancers associated with upregulated genes, suggesting nucleosome displacement by AR. Consistent with H3K27Ac profiling at regulated genes (**Figs. 4, S5, S6**), we observed a temporary (at 4h) increase of H3K27Ac in Daxx KO compared to parental cells. AR KO completely abolished, and HIRA KO strongly reduced, accumulation of H3K27Ac at enhancers associated with both up- and down-regulated genes (**Figs. 7** **and S10)**.

**Fig. 7.**
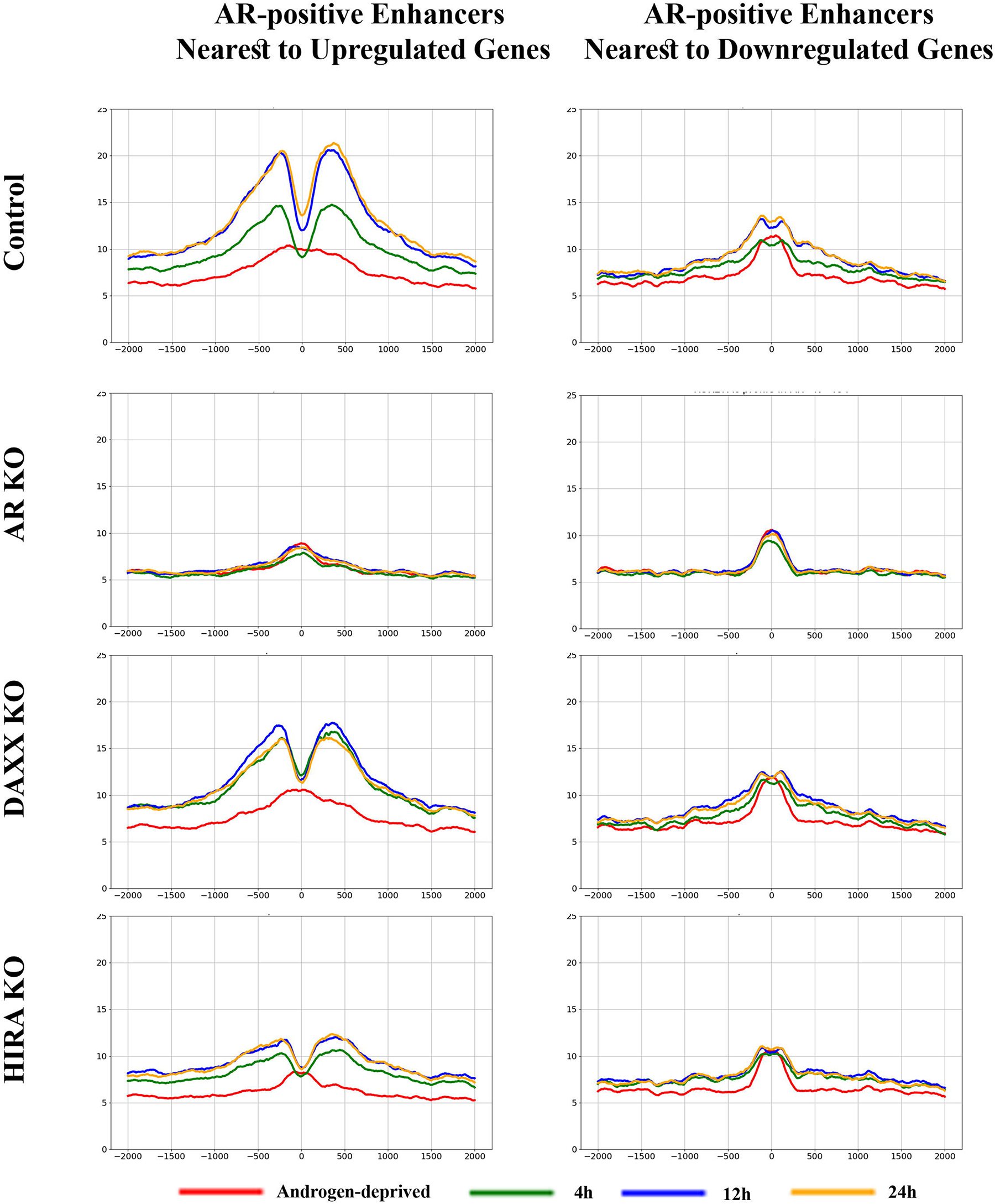
Dynamics of H3K27Ac at enhancers. Metaplot of H3K27Ac ChIP-seq analysis at AR peaks (position “0”: AR at 4h in R1-AD1 parental cells) at enhancers associated with up-(left) and down-regulated (right) genes in R1-AD1 parental (Control), AR KO, Daxx KO, and HIRA KO in androgen-deprived (72h, red) and R1881 stimulated for 4h (green), 12h (blue), 24h (yellow). In parental cells, H3K27Ac is gradually elevated during androgen treatment, to higher levels at enhancers associated with up-regulated genes. AR KO completely abolished, and HIRA KO strongly reduced accumulation of H3K27Ac at enhancers associated with both up- and downregulated genes. See **Fig. S10** for time points comparison.

Our results suggest H3.3 role in H3K27 acetylation at enhancers. Phosphorylation of H3.3S31 (H3.3S31Ph), a unique H3.3 residue, regulates activation of histone acetyltransferase p300 resulting in acetylation of H3K27 at enhancers (29). We observed androgen-induced increase of H3.3S31Ph in parental and Daxx KO cells, no changes in AR KO cells, and a reduction of H3.3S31Ph in HIRA KO cells (**Figs. 8** **and S11** for time points comparison), that, together with H3.3 data, suggests function of this histone variant in acetylation of H3K27 at enhancers.

**Fig. 8.**
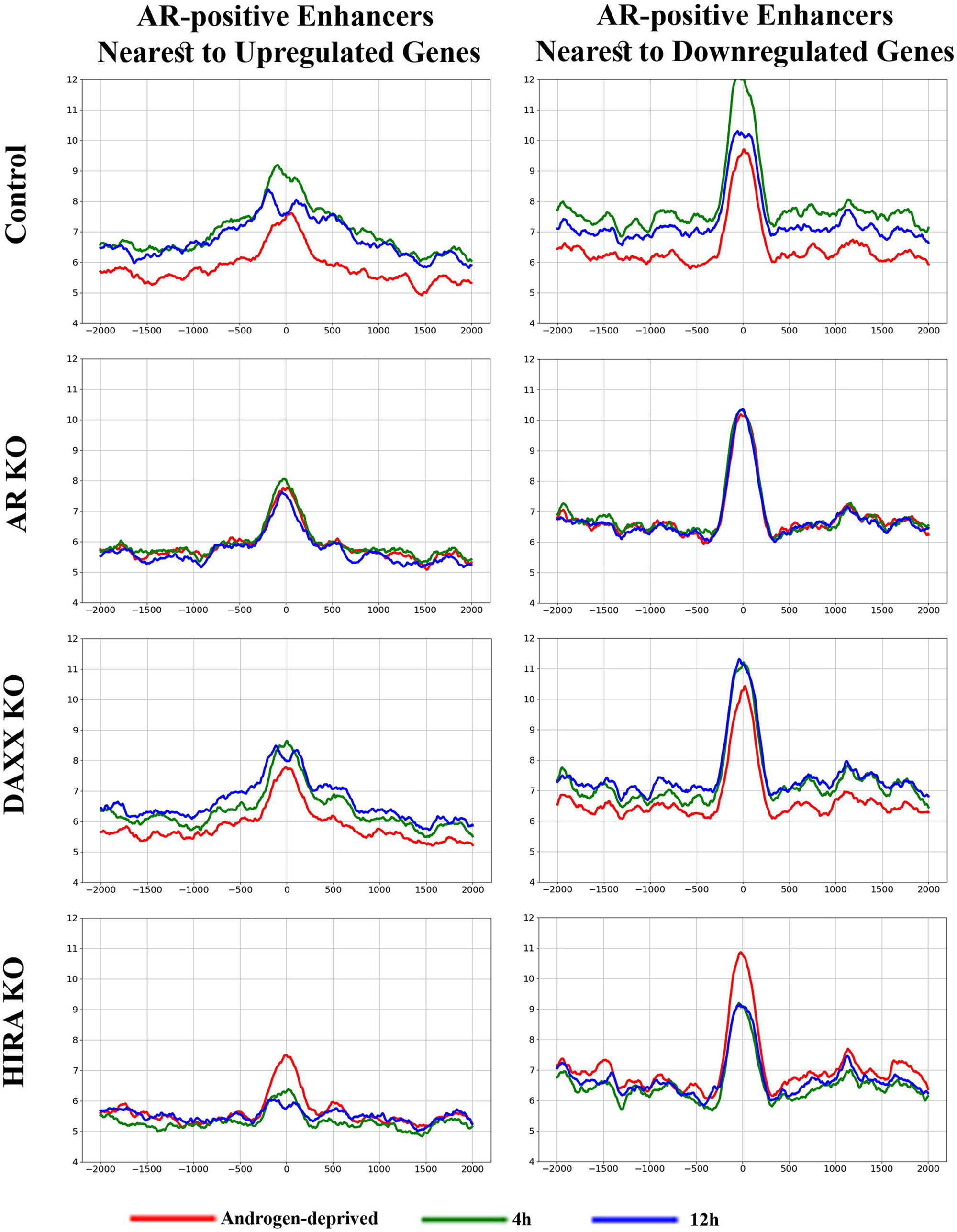
Dynamics of H3.3S31Ph at enhancers; analysis within cell lines. Metaplot of H3.3S31Ph ChIP-seq analysis at AR peaks (position “0”: AR at 4h in R1-AD1 parental cells) at enhancers associated with up-(left) and down-regulated (right) genes in R1-AD1 parental (Control), AR KO, Daxx KO and HIRA KO in androgen-deprived (72h, red) and R1881 stimulated for 4h (green), 12h (blue). Levels of H3.3S31Ph are androgen-induced in parental and Daxx KO cells, no changes in AR KO cells, and are reduced in HIRA KO cells, resembling H3.3 dynamics. See **Fig. S11** for time points comparison.

### HIRA KO affects BRD4 binding to enhancers

Epigenetic reader bromodomain protein BRD4 is indispensable for enhancer activity (62). It acts as a scaffold protein recruiting transcription factors for transcription activation, including enhancer-promoter interaction (59,63). BRD4 interacts with AR(64) and is required for AR-mediated transcription. Inhibition of BRD4 abrogates AR binding to AREs(64) and enhancers enriched with H3K27Ac recruit BRD4 to chromatin (62). In parental and Daxx KO cells, androgen treatment induced BRD4 accumulation at AR binding sites within enhancers associated with up- and down-regulated genes, with dynamics mirroring those of H3K27Ac and AR (**Fig. 9** **and S12** for time points comparison; levels of BRD4 were monitored by Western blot analysis, **Fig. S1C**). Increased BRD4 at 4h in Daxx KO can be explained by elevated H3K27Ac at this time point (**Fig. S10**). Reduced H3K27Ac by AR KO and HIRA KO (**Figs. 4** **and S10**) would predict for decreased BRD4 accumulation in these cell lines. Indeed, we observed that AR KO abolished BRD4 accumulation, confirming H3K27Ac function in BRD4 binding to chromatin (compare **Figs. 7** and **9**) and illuminating a role for AR in BRD4 recruitment at enhancers that is reciprocal to reported BRD4-dependent AR recruitment (64). In HIRA KO cells, BRD4 dynamics are similar to AR (compare **Figs. 9** and **5, S7** and **S12**): it is elevated at 4h of stimulation (potentially by co-recruitment with its interaction partner AR (64)), but is reduced to the androgen deprived levels at 12h (**Figs. 9** **and S12** for time points comparison). Together, these data suggest new H3.3/H3K27Ac function in dynamics of BRD4 and AR at enhancers and imply a two-step model of recruitment and retention of AR transcription complex at enhancers (see Discussion for details).

**Fig. 9.**
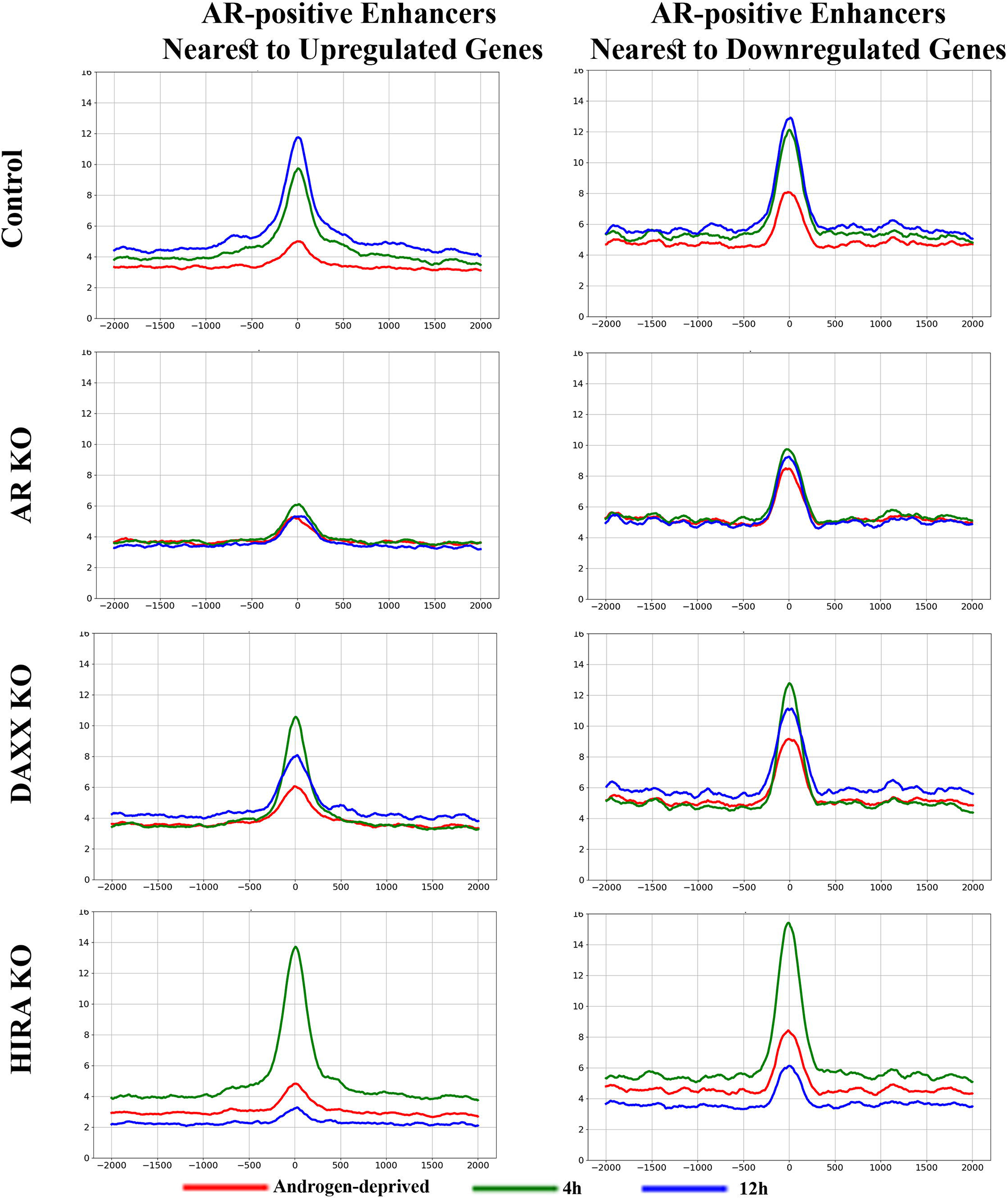
Dynamics of BRD4 at enhancers. Metaplot of BRD4 ChIP-seq analysis at AR peaks (position “0”: AR at 4h in R1-AD1 parental cells) at enhancers associated with up-(left) and down-regulated (right) genes in R1-AD1 parental (Control), AR KO, Daxx KO, and HIRA KO in androgen-deprived (72h, red) and R1881 stimulated for 4h (green), 12h (blue). BRD4 accumulated within enhancers at AR binding sites in parental cells, with dynamics mirroring those of AR (Fig. 2). AR KO abolished BRD4 accumulation. In HIRA KO cells, BRD4 dynamics is similar to AR, first accumulating at 4h of stimulation and next reducing at 12h. See **Fig. S12** for time points comparison.

### HIRA KO reduces DNA accessibility at AREs

Changes in AR binding in HIRA-KO cells may suggest alteration of DNA accessibility that was tested by ATAC-seq. Androgen stimulation for 4h increased DNA accessibility at AR binding sites within enhancers associated with up-(to the higher extend) and down-regulated genes in all but AR KO cells (**Figs. 10** and **S13** for time points comparison), perhaps due to the binding of pioneer transcription factors (65) that results in nucleosome shift/eviction as seen by the loss of H3.3 peaks at this time point (**Figs. 6, S9**). H3.3 is associated with elevated DNA accessibility at the regulatory elements (19) that affects the binding of transcription factors as was recently shown at promoters in ES cells (53). We observed that HIRA KO cells, in line with reduction of H3.3, have decreased DNA accessibility under androgen-deprived conditions at AR-binding sites within enhancers (**Fig. S13**). Therefore, elevated AR binding at 4h cannot be explained exclusively by pre-existing DNA accessibility due to reduced H3.3 nucleosomes loading in these cells.

**Fig. 10.**
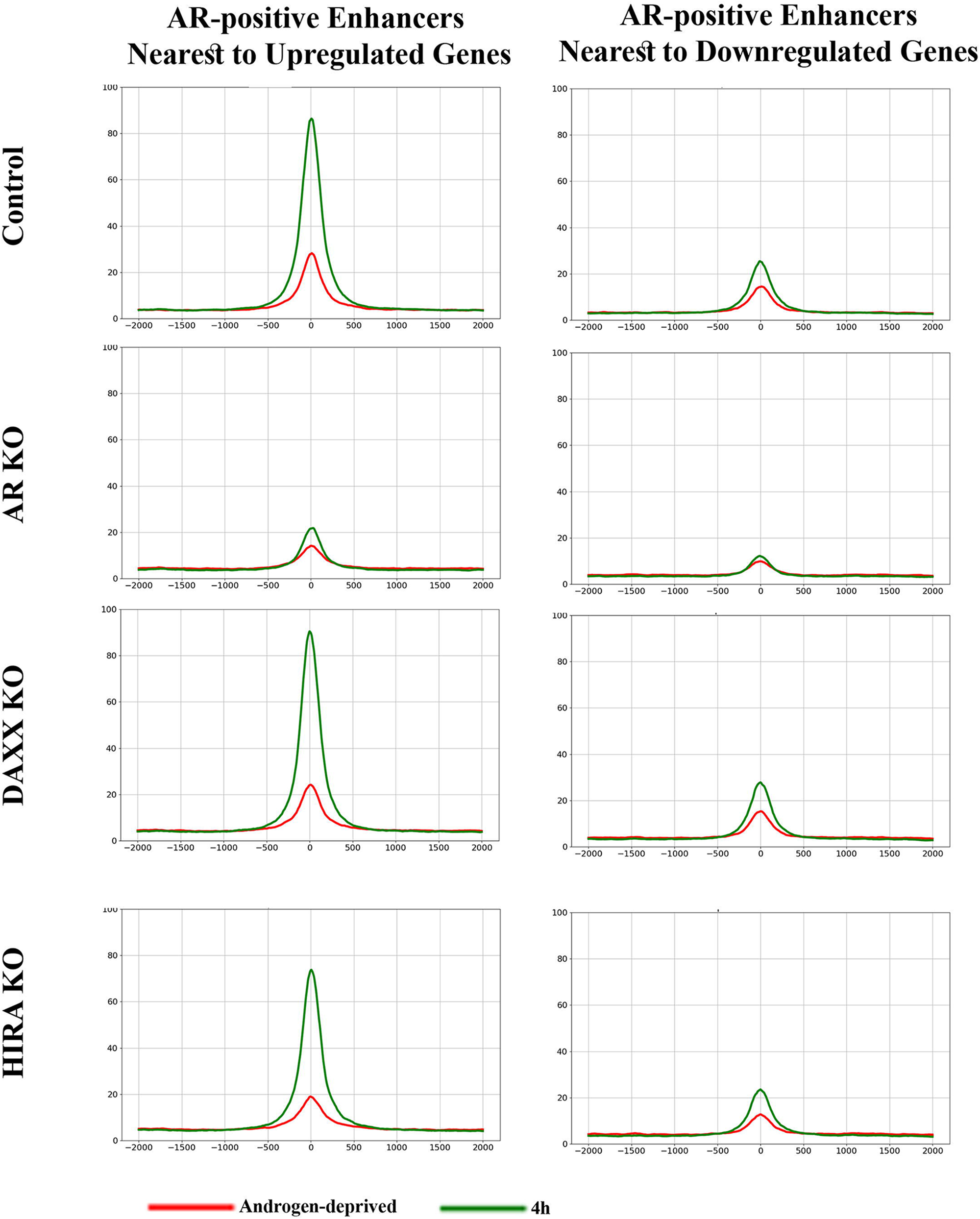
DNA accessibility analyzed of AR-positive enhancers by ATAC-seq. Metaplot of DNA accessibility at AR peaks (position “0”: AR at 4h in R1-AD1 parental cells) at enhancers associated with up-(left) and down-regulated (right) genes in R1-AD1 parental (Control), Daxx KO and HIRA KO in androgen-deprived (72h, red) and R1881 stimulated for 4h (green). Stimulation for 4h increased DNA accessibility at AR binding sites within enhancers associated with up-(to the higher extend) and downregulated genes, in all but AR KO cells; HIRA KO reduces DNA accessibility. See **Fig. S13** for time points comparison.

### HIRA KO abolishes induction of FKBP5 gene and affects epigenetics profiling at FKBP5 superenhancer

FKBP5 is a co-chaperone that, among other functions, regulates AR dimerization (66). Several AREs were identified upstream and downstream from FKBP5 TSS (67,68). FKBP5 gene transcription is induced by androgens (67), which was confirmed in our experimental system (**Fig. 11A**). AR KO abolished androgen induction of FKBP5 at the RNA and protein levels. While Daxx KO had minimal effect on FKBP5 induction, HIRA KO reduced androgen induction of FKBP5. To investigate mechanism of HIRA-dependent regulation, we performed epigenetic analysis of AR-binding region upstream of the main FKBP5 TSS (69). H3K4me1 is elevated in a 20 kB region located 50 kB upstream of the main FKBP5 TSS (**Figs. 11B** and **S14B** for the full-size images) at the position of a putative enhancer that was previously identified using reporter assay in non-small cell lung cancer cells (70). The ROSE (Rank Ordering of Superenhancers) protocol (41,71) analysis identified this region as a superenhancer (SE). HIRA KO reduced the size of H3K4me1-positive region, reproducing a tendency observed on other enhancers (**Fig. S8**). H3K27Ac ChIP-seq was used to evaluate status of this SE in androgen-deprived and androgen-induced conditions. In parental cells, H3K27Ac and H3K4me1 peaks overlap, confirming the active status of FKBP5 SE. H3K27Ac accumulated gradually during androgen induction (**Figs. 11B** **and S14B**) thus reproducing H3K27Ac dynamics at AR-positive enhancers associated with upregulated genes ((**Figs. 7** **and S10**). In AR KO cells, H3K27Ac is drastically reduced in androgen-deprived and stimulated conditions, indicating a poised SE. Daxx KO had minor effects on H3K27Ac at FKBP5 SE. Similar to the profile of gene bodies and enhancers, we observed a temporary (at 4h) elevation of H3K27Ac in Daxx KO compared to parental cells. HIRA KO strongly reduced H3K27Ac in androgen deprived and stimulated conditions. We next profiled H3.3 and H3.3S31Ph at FKBP5 SE. In parental cells,

**Fig. 11.**
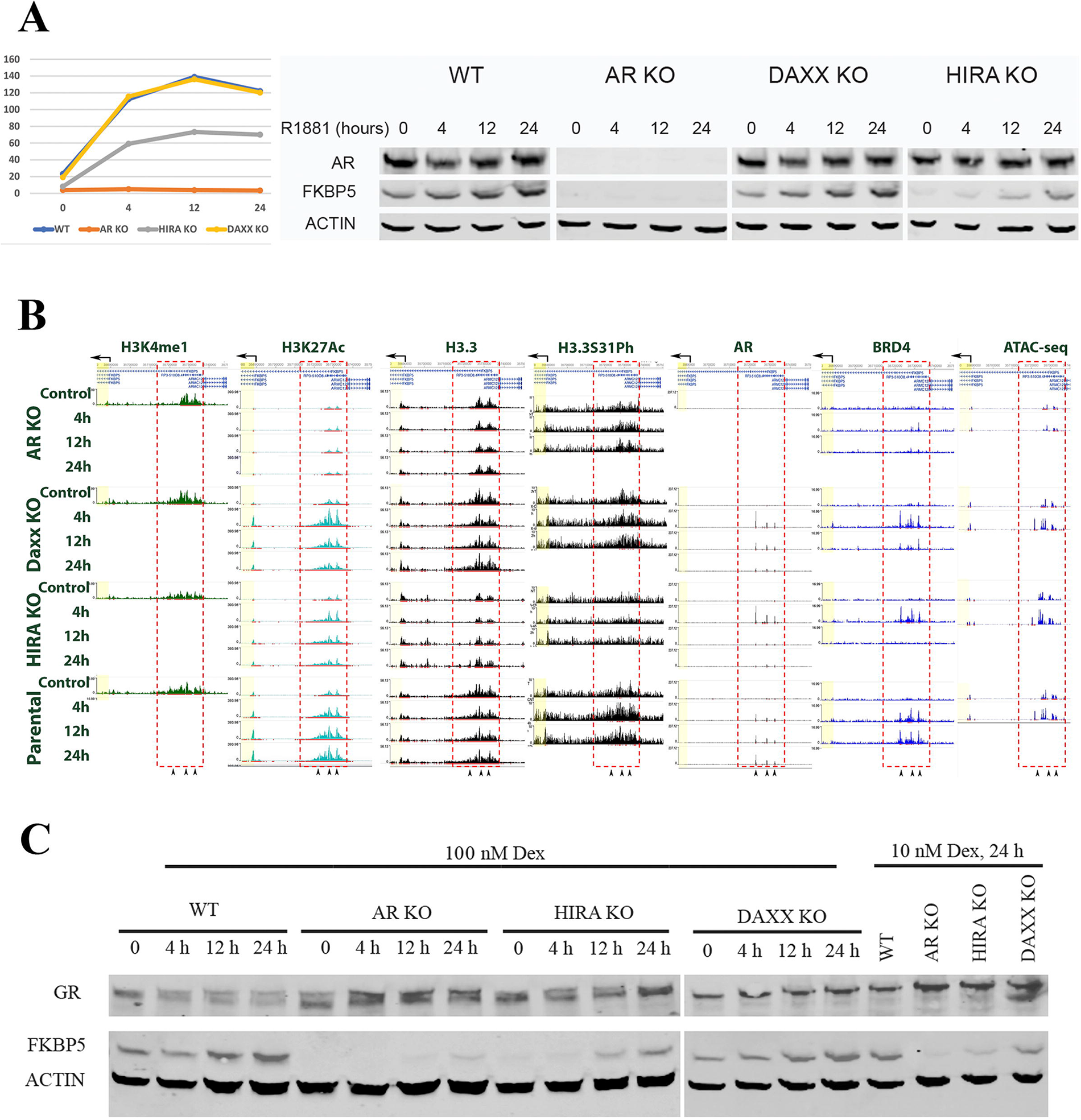
FKBP5 gene and SE are regulated by AR, GR and HIRA. **A.** Expression of FKBP5. R1-AD1 cells (parental, AR KO, Daxx KO and HIRA KO) were androgen-deprived for 72h (control), induced with 1 nM of R1881 for 4h, 12h, 24h and subjected by FKBP5 RNA analysis at three bioreplicates (left) and Western blot analysis with corresponding antibodies (right). AR KO and HIRA KO substantially reduced induction of FKBP5 at RNA and protein levels. **B.** Epigenetic profiling of FKBP5 SE. R1-AD1 cells (parental, AR KO, Daxx KO, HIRA KO) were androgen-deprived for 72h (Control) and induced with 1 nM of R1881 for 4h, 12h, 24h. H3K4me1 (enhancer), H3K27Ac (active enhancer), H3.3, H3.3S31Ph, AR, and BRD4 profiling were analyzed by ChIP-seq. DNA accessibility analyzed by ATAC-seq. **C.** HIRA KO abolishes induction of FKBP5 gene by glucocorticoid receptor (GR). R1-AD1 cells (parental, AR KO, Daxx KO, HIRA KO) were hormone-deprived for 72h (0), induced with indicated concentrations of dexamethasone (Dex) for indicated time, and analyzed by Western blot with corresponding antibodies. AR KO elevates GR levels; AR and HIRA KO substantially reduced induction of FKBP5 protein levels by Dex. Actin: loading control.

H3.3 was enriched at FKBP5 SE (**Figs. 11B** **and S14B**). Whereas Daxx KO had minor effects on H3.3 deposition, HIRA KO strongly reduced H3.3 at SE. H3.3S31Ph was accumulated at FKBP5 SE during induction in parental cells and, similarly to H3.3, was substantially reduced in HIRA KO cells (**Figs. 11B** **and S14B**).

### HIRA KO affects AR and BRD4 binding to FKBP5 SE

Three AR peaks that were previously mapped in VCaP and LNCaP cells (69) within FKBP5 SE identified above are conservative in R1-AD1 cells (**Fig. 11B**). In parental cells, AR is chromatin-associated at all induction time points at all three peaks, while in HIRA KO it is induced at 4h and is diminished at 12h and 24h (**Figs. 11B** **and S14B**), in concordance with dynamic of AR at enhancers (**Figs. 5** **and S7**). We also observed that BRD4 accumulates at all three AREs within FKBP5 SE in parental and in Daxx KO cells during induction (**Figs. 11B** **and S14B**). AR KO abolished BRD4 accumulation, confirming AR and H3K27Ac function in BRD4 recruitment. In HIRA KO cells, BRD4 accumulates at 4h of stimulation and disappears at 12h, resembling dynamics of AR. In summary, HIRA/ H3.3 dependent epigenetic changes observed at FKBP5 SE confirmed results of metaplot profile analyses. Same tendencies were documented at enhancers of additional genes deregulated by AR and HIRA KO (e.g., androgen up-regulated IL1R1, LONRF1, and down-regulated ID2; **Fig. S14**).

### HIRA KO affects DNA accessibility at FKBP5 SE

Next we analyzed DNA accessibility at three AR binding sites within FKBP5 SE. PWMtools analysis (https://ccg.epfl.ch/pwmtools/) identified half-site ARE (72) at the peak 1 (ARE-1, **Fig. S15**) and canonical AREs (72) at peaks 2 and 3 (ARE-2 and -3, **Fig. S15**). Androgen stimulation for 4h induced an ATAC peak at the ARE-1 (half-site) in parental, Daxx KO and HIRA KO cells (**Figs. 11B, S14 and S15**), indicating nucleosome shift/eviction at this half-site ARE. Pioneer transcription factor FOXA1 opens compact chromatin, increasing DNA accessibility (73). It interacts with AR (74), thereby regulating AR binding to DNA, mostly at half-site ARE, that requires FOXA1 for AR deposition to AREs resulting in nucleosomes displacement (75). FOXA1 binding consensus was observed to overlap with ARE-1 (p: 1.55e-06), suggesting FOXA1 function for chromatin opening and AR binding to this ARE. Therefore, increased DNA accessibility at this site can be explained by the androgen-induced binding of FOXA1 or another pioneer transcription factor that precluded AR binding. In androgen-deprived conditions, ARE-2 is more accessible in HIRA KO cells compared with parental cells (**Fig. S15**), suggesting an H3.3-dependent partial block of AR binding at some ARE, that is abrogated by HIRA KO.

### HIRA KO does not affect CTCF binding

DNA binding protein CTCF regulates chromatin structure, including formation of topology associated domains (TADs) (76,77) that are important to maintain interchromatin interactions, including those of promoters/enhancers. Potential participation of CTCF in HIRA/H3.3-dependent chromatin structure was addressed by CTCF ChIP-seq. We identified two CTCF peaks at the 3’ end of FKBP5 SE and additional two peaks within FKBP5 gene body. None of these peaks were substantially affected in all KO cells, including HIRA KO (**Fig. S14B**), therefore excluding CTCF changes as a mechanism of deregulated FKBP5 expression.

### HIRA KO abolishes induction of FKBP5 gene by glucocorticoid receptor (GR)

AR and GR have diverse physiological functions, yet their activation mechanisms are similar. Both of these nuclear receptors bind to almost identical DNA elements and are associated with a similar set of co-activators and co-repressors (78). In addition, both AR and GR induce expression of FKBP5 (69). Given this similarity, we asked whether H3.3/ HIRA dependent mechanism of AR-driven transcription regulation can be extrapolated to GR. To this end, we tested effect of GR induction by dexamethasone (dex) on expression of FKBP5. Using the same cell lines and induction time point as in **Fig. 11A**, we observed that GR induced FKBP5 similarly to AR in parental and Daxx KO cells (compare **Figs. 11** **A and C**). AR-KO elevated GR expression (by RNA-seq) and protein accumulation (**Fig. 11C**), confirming a function of AR in suppression of GR expression (79,80). Surprisingly, AR KO, while elevated GR accumulation, almost completely abolished FKBP5 induction by dex, suggesting functional connection between AR and GR. HIRA KO strongly reduced FKBP5 induction (**Fig. 11C**), indicating that H3.3/ HIRA dependent mechanism is common for AR and GR, and, potentially, for other members of nuclear receptors family.

### In AR-V expressing CRPC cells, HIRA KO is lethal, and HIRA KD affects H3.3 levels at AR-positive enhancers associated with AR-regulated genes

Gain-of-function mutations of AR (81) are implicated in the development of CRPC and in therapy resistance (82). A recently identified molecular mechanism of aberrant AR activation in CRPC is expression of AR variants (AR-Vs) with deleted LBD (AR-DLBD) (6). Clinical data and animal models confirm function of AR-DLBD in CRPC (6). Expression of AR-Vs, including AR-DLBD, was identified in metastatic CRPC(83); it facilitates treatment resistance to anti-androgens (6), and is associated with negative survival prognoses. Identification of co-regulators of AR-Vs may yield novel targets for CRPC treatment. To this end, we compared the effect of manipulation of the H3.3 pathway in isogenic R1-AD1 (AR-WT) and R1-D567 (AR-DLBD) cells (33). Characterizing these cells, we observed that AR-WT is retained in the cytoplasm and relocates to the nucleus upon androgen treatment (**Fig. S16A**). Distinctly, AR-DLBD is a nuclear protein under both androgen-deprived and androgen-treated conditions, confirming ligand-independent nuclear localization of AR-V. We modified R1-D567 cells for expression of FLAG/HA-tagged H3.3 (**Fig. S16B**) and produced AR KO and Daxx KO cells (**Fig. S16C**). In three independent experiments we failed to produce HIRA KO in R1-D567 cells. HIRA gRNA deleted both alleles with similar efficiency in R1-AD1 and R1-D567 cell lines (by sequencing and IF). Nonetheless, R1-D567 (AR-DLBD) cells did not survive HIRA KO during clone expansion. As R1-AD1 and R1-D567 cells are isogenic and differ only in AR (AR-WT vs AR-DLBD), our data suggest that CRPC cell lines are HIRA-dependent and those that express AR-DLBD (and, potentially, other AR-Vs) developed HIRA addiction, suggesting synthetic lethality. HIRA KD (combination of shRNA and siRNA) reduced cell survival (tested by colony formation assay) much stronger in R1-D567 compared with R1-AD1 cells (**Fig. S16D**). As in R1-AD1 cells (**Fig. S1**), downregulation of HIRA reduced H3.3 (**Fig. S16C**).

We next analyzed H3.3 association with AR-positive (using AR binding data in R1-D567 (33)) enhancers (using H3K4me1 ChIP-seq analysis in R1-AD1 cells) nearest to genes up- and down-regulated by AR KO compared to parental R1-D567 cells (identified by RNA-seq). In AR KO cells, H3.3 is reduced at enhancers associated with both up- and down-regulated genes (**Fig. S16E**) in a manner similar to H3.3 profiling in R1-AD1 (**Figs. 6, S9**), confirming AR-mediated transcription in maintenance of this histone variant. As in R1-AD1 cells, we observed a major reduction of H3.3 in HIRA KD R1-D567 cells (**Fig. S16E**), validating H3.3/ HIRA pathway in PC cells expressing AR-V.

### HIRA complex deregulation in PC

HIRA accumulation induces breast cancer metastasis by H3.3-dependent aggressive transcriptional reprogramming (84). Exploring the function of H3.3 in PC (TCGA and the Human Protein Atlas), we found that expression of two HIRA complex components, proteins HIRA and UBN1, is increased in tumor compared to normal prostate tissue (**Fig. S17A**). HIRA expression was also elevated in high/very high-risk PC groups as determined by Gleason scores (**Fig. S17A**). Accumulation of HIRA (**Fig. S17B, C)** and UBN1 (85) is associated with negative PC survival prognosis. HIRA expression was elevated in glandular epithelial cells, which are the main source of prostate adenocarcinoma (single cell analysis, the Human Protein Atlas https://www.proteinatlas.org; **Fig. S17D**). Levels of BRD4 (86) and p300 (87,88) are significantly elevated in high-risk PC patients. p300 and BRD4 inhibitors downregulate AR target gene expression and abrogate PC growth (64,89,90).

HIRA depletion obliterates PC growth *in vitro* and in xenograft settings, deregulates androgen-induced expression and reduces AR binding at enhancers of target genes. In addition, CRPC cells expressing gain-of-function AR variants (AR-Vs, implicated in antiandrogen resistance and development of metastatic CRPC (mCRPC)) are HIRA-dependent. Combined with our results (**Fig. 1**), these data suggest potential oncogenic function of HIRA-dependent H3.3 pathway in PC initiation and progression and point at H3.3/ HIRA pathway as a potential target for therapeutic intervention. Several inhibitors of CHK1, p300 and BRD4 are currently in clinical trials (phase 1 to 3), including p300 inhibitor CCS1477 (Inobrodib) (91), BRD4 inhibitor AZD5153 (92), and CHK1 inhibitors SRA737 (93). Hence, inhibition of BRD4, p300, CHK1 and IKKα is predicted to impair progression of PC by inactivating BRD4-dependent enhancers and the consequent decline of AR-driven transcription by reduced AR binding to AREs. Indeed, we observed that HIRA KO increases sensitivity of PC cells to BRD4 inhibition by JQ1 in cell culture and in 3D prostasphere models (**Fig. S18**), suggesting that reduced chromatin affinity of BRD4 makes HIRA KO cells more vulnerable to BRD4 inhibition.

## Discussion

Chromatin enriched in histone H3 variant H3.3 and histone H2A variant H2A.Z (25) is associated with elevated transcription activity (18). According to Jin and Felsenfeld (19), H3.3/H2A.Z-containing nucleosomes are less stable compared to nucleosomes with canonical histones, thereby providing DNA accessibility to the transcription machinery at the regulatory elements. During the last decade, multiple experimental evidences were accumulating that further explain H3.3 function in transcription regulation, including H3.3 association with transcriptionally active markers (H3K9Ac and H3K4me3 (9,14,20,21)), prevention of silencing markers (22,23), and recruitment of p300 activity for H3K27Ac at enhancers (29). These findings emphasize transcription-associated properties of H3.3, and recent results from the Banaszynski group demonstrated H3.3-dependent transcription factors recruitment to regulated elements (mostly TSS) of transcribed genes (53). In the current study, we demonstrated function of H3.3 chaperone HIRA in dynamic regulation of the main PC driver, nuclear receptor AR. Depletion of HIRA prevents PC cells growth *in vitro, in vivo* and deregulates androgen-induced expression. H3.3 is enriched at boundaries of highly expressed genes (14) and at enhancers (28). Investigating mechanisms of this HIRA-dependent deregulation, we found that HIRA KO reduces H3.3 incorporation on androgen regulated genes and on AR-positive enhancers, confirming function of HIRA in deposition of H3.3 at multiple genome elements in PC models with AR-WT and AR-DLBD. Androgen treatment induces accumulation of H3K27Ac at gene bodies, TSS and enhancers. We also observed that H3K27Ac levels first increase at enhancers (4h) and then at TSS (12h), suggesting potential transition of histone acetyl transferase activity from one genome element to another. HIRA KO diminishes levels of H3K27Ac in androgen-deprived conditions and reduces accumulation of this modification during androgen induction on regulated genes and nearby enhancers. Recent findings uncovered a function of H3.3 in activation of p300 that results in acetylation of H3K27 at enhancers (29). This function requires phosphorylation of H3.3S31 (H3.3S31Ph), a unique H3.3 residue, by the checkpoint kinase CHK1 (29),(30) and the NF-κB kinase IKKα (31). These findings are consistent with our observations that reduction of H3.3 and H3.3S31Ph correlates with diminished levels of H3K27Ac in HIRA-KO cells.

Further analysis of HIRA-dependent deregulation of androgen-induced transcription revealed that HIRA KO changes accumulation dynamics of AR and BRD4 with enhancers associated with target genes. Specifically, AR and BRD4 levels were increased at 4h after androgen stimulation followed by a reduction to pre-stimulation levels at later time points. Reduction of H3.3 at enhancers in HIRA KO cells did not increase accessibility of most AREs in androgen deprived conditions, therefore excluding the simple explanation of elevated AR binding at 4h of induction in these cells. Instead, our data suggest a two-step assembly mechanism of functional AR complex at enhancer. At the first step (“recruitment”), AR interacts with AREs (in some cases, with help of pioneer factors like FOXA1, as, potentially, at ARE-1 in FKBP5 SE) and BRD4 is co-recruited with AR via protein/protein interaction (64). At the next step, (“retention”), BRD4 is transitioned from AR to H3K27Ac nucleosomes of enhancer. This results in stabilization of complex (including formation of multi-protein phase separation domain (94)) that may prevent dissociation of both BRD4 and AR from AREs, resulting in formation of active AR transactivation complex at enhancers and gene regulation (**Fig. 12**). Confirming this model, it was shown that BRD4 is sufficient for AR-mediated transcription and BRD4 inhibition by JQ1 reduced both BRD4 and AR binding and androgen-dependent transcription (64). Our results documented interdependence between AR and BRD4 recruitment and dynamics of complex assembly, importantly, in the context of H3.3/ HIRA pathway: HIRA KO elevated the “recruitment” and abrogated the “retention”, thus separating the first and the second (H3.3-dependent) steps of transactivation complex assembly suggested by our model (**Fig. 12**).

**Fig. 12.**
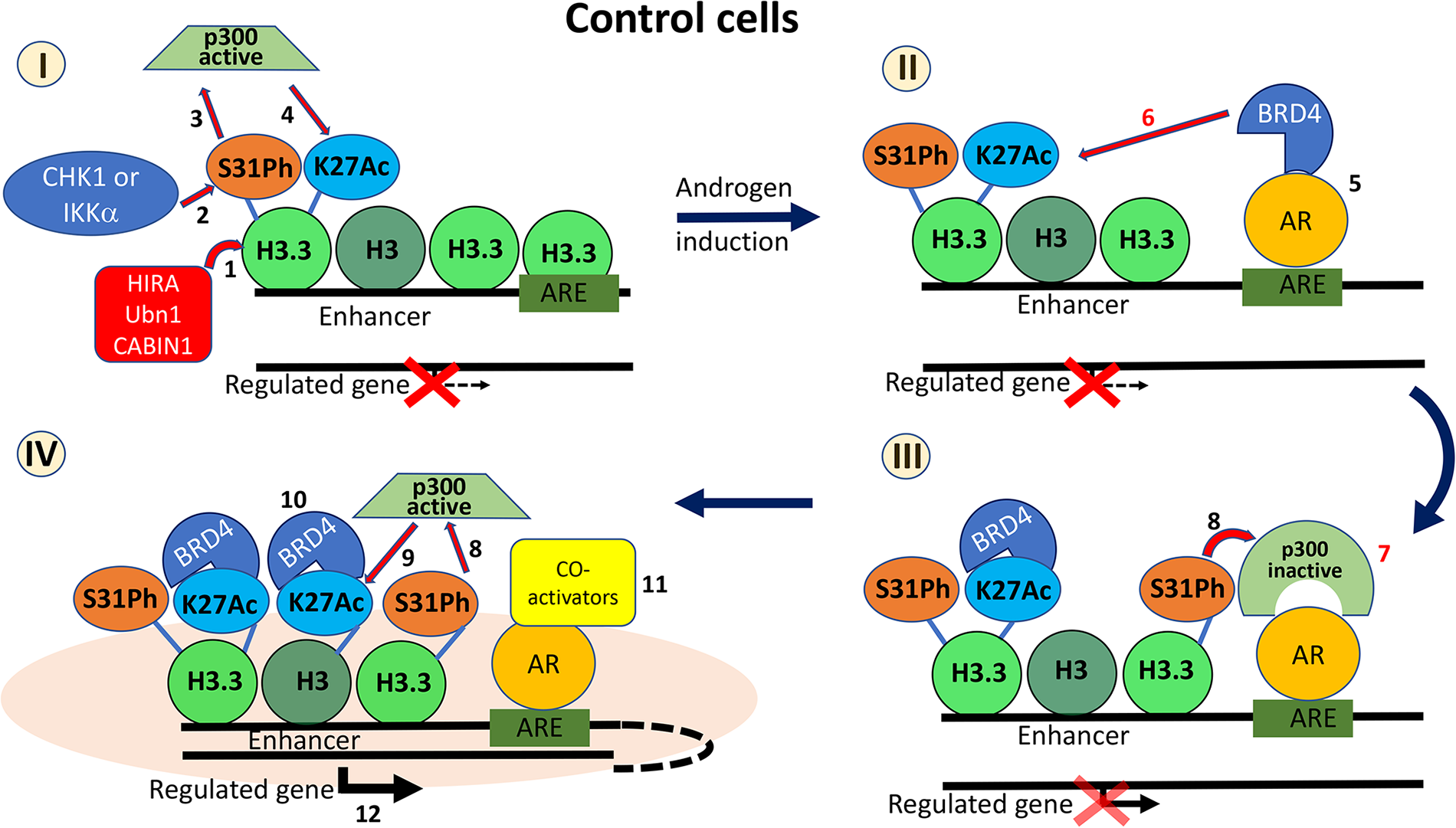
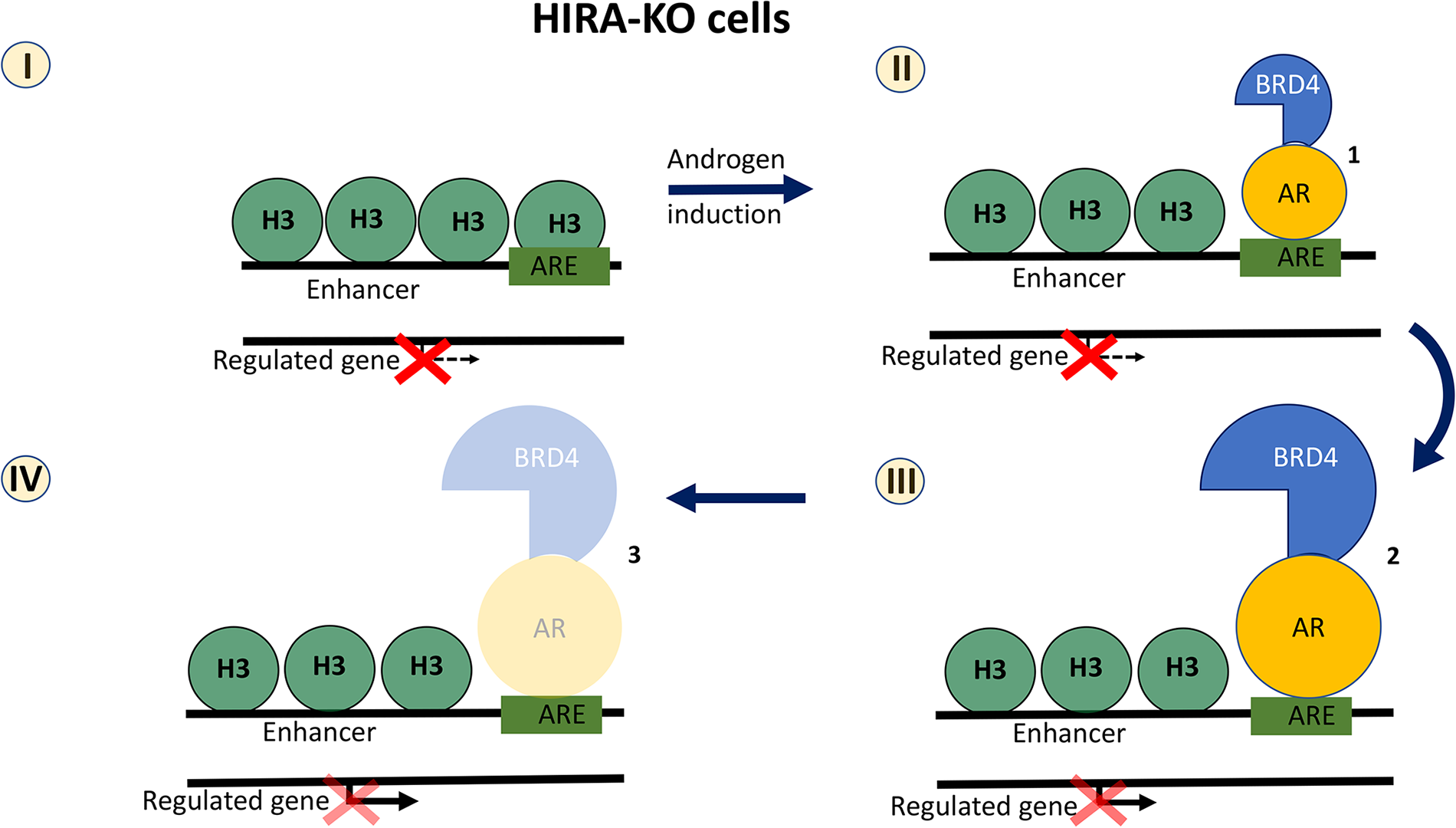
Model of HIRA/ H3.3 function in AR transcription regulation. Control cells. **I:** HIRA chaperone complex recruits H3.3 at enhancers (1). CHK1 and IKKa phosphorylates H3.3 at S31 (S31ph; 2), that activates p300 (3) for acetylation of H3K27 (4). **II:** androgen treatment recruits AR to ARE that interacts with and co-recruits bromodomain protein BRD4 and displaces nucleosome at ARE (5); BRD4 binds to H3K27Ac at enhancer (6). **III:** release of BRD4 from AR opens AR-NTD for recruitment of p300 (7) that is activated by increased levels S31ph (8, similar to 3). **IV:** active p300 further acetylates H3K27 (9, similar to 4), resulted in additional binding of BRD4 to H3K27Ac at enhancers (10, similar to 6), recruitment of Co-activators (11) and formation of completed transcription complex (pink cloud), enhancer-promoter contact and transcription activation (12). Steps 6 and 7 are speculative and thus highlighted in red. Model does not reflect several details and complementary scenarios, including but not limiting: effect of AR-dimer, co-recruitment of p300 and AR co-regulators, transfer of PolII from enhancer to promoter, BRD4-mediated transcription elongation. **HIRA-KO cells. I:** no H3.3 recruitment, reduced H3K27Ac. **II:** androgen induction recruits AR that interacts with and co-recruits BRD4 at AREs (1). **III:** BRD4 cannot bind to chromatin due to reduced H3K27Ac, thus accumulating together with AR at AREs, and blocking recruitment of AR co-regulators (2). **IV:** in the absence of co-regulators and reduced BRD4 binding to H3K27Ac at enhancers, AR and BRD4 have decreased retention at AREs (3); AR transcription complex does not form, and transcription regulation is weaker compared to parental cells.

The combination of RNA-seq and ChIP-seq results indicated that at 4h of androgen induction, when both AR and BRD4 are enriched at enhancers in HIRA KO cells, androgen-induced transcription is minimal in comparison with parental cells. Why the highly elevated AR and BRD4 are not able to regulate transcription? Our data suggest that recruitment of both AR and BRD4 in H3.3-reduced environment (e.g., in HIRA KO cells) is insufficient for the proper transcription regulation, potentially due to several (hypothetically interconnecting) possibilities: 1) Due to the reduced H3K27Ac, BRD4 cannot bind to/retained at enhancers. BRD4 is required for AR activity (64) and BRD4 inhibition abrogates AR binding to AREs (64) that may reduce AR retention and transcription regulation; 2) Deregulation of BRD4 binding may induce changes in chromatin structure. Enhancers/ SE are located distal to regulated genes (up to several megabases (95)) and they activate transcription by loop formation (96). BRD4 is indispensable for enhancer/SE activity, including enhancer/gene contact (71,97). Formation of chromatin loop between TSS and SE was previously proposed in activation of FKBP5 gene (70). Together with our emerging results, it suggests function of HIRA/ H3.3/ BRD4/ AR in chromatin structure regulation, specifically in establishing enhancer/SE contact with regulated genes that may be abrogated in HIRA KO cells; 3) HIRA KO can deregulate recruitment of AR co-activators or co-repressors. AR binds to BRD4 via NTD (aa 94 – 186 of AR), while BRD4 binds to AR via N-terminus that contains bromodomains BD1 and BD2 (64). In HIRA KO cells, levels of H3K27Ac are reduced at enhancers in androgen-deprived and androgen-induced conditions. It may diminish BRD4 binding to chromatin via H3K27Ac, thus hampering its dissociation from AR and blocking AR-NTD for recruitment of co-repressors or co-activators, including p300 that, in parental cells, may create a positive loop, further acetylating H3K27 and promoting BRD4 binding to enhancers; 4) HIRA/ H3.3 pathway regulates binding of PolII at TSSs (53) that correlates with reduced DNA accessibility in HIRA/ H3.3 KO ESC (53). We observed that HIRA KO cells reduced DNA accessibility (by ATAC-seq) at TSSs of AR-regulated genes, in androgen deprived and stimulated conditions (e.g., ATAC-seq at FKBP5 TSS, **Fig. S14B**), suggesting reduced PolII binding. Our results suggest additional H3.3-dependent mechanism(s) that promotes PolII transition from enhancer to TSS via promoter/enhancer contact within phase separated domains that are induced by BRD4 accumulation ((97); reviewed in (98)), or transition of PolII from initiation to elongation step, that is promoted by BRD4 ((99); reviewed in (100)). Altogether, our model opens several questions that require further investigation.

A recent publication demonstrated that H3.3 is sufficient for H3K27Ac enrichment and binding of BRD4 and transcription factors at promoters in ESCs (53). We observed that HIRA KO deregulates AR- and GR-driven transcription, suggesting common H3.3/ HIRA-dependent mechanism of nuclear receptors function at enhancers. Surprisingly, AR KO reduced GR activation response despite accumulation of GR protein (**Fig. 11C**), suggesting functional interplay between two pathways. One possible explanation is that inactivation of enhancers by AR KO (e.g., reduction of H3K27Ac, **Figs. 7, 11**) diminishes activity of another nuclear receptor, GR, that will bind to the similar DNA elements within these enhancers during activation by dex. Our data indicate that deregulation of enhancer activity by AR disfunction may require several cell cycles to deregulate enhancers, potentially by the negative feedback between reduced enhancer transcription that results in diminished transcription-associated loading of H3.3 (50–52), followed by H3.3-dependent H3K27Ac decrease. This model suggests functional connection between AR and GR and is important in the context of GR role in the enzalutamide resistant CRPC (79).

Advanced PC is a terminal disease and despite decades of intense laboratory- and clinical-based investigations, to date there is no cure or long-lasting effective treatment options. Initiation and progression of PC, including transition to CRPC, is determined by, among other factors, transcription reprogramming that is controlled by epigenetic dysregulation. Multiple lines of evidence point to the essential function of H3.3 deposition pathways in initiation and progression of bone tumors (101), pancreatic (102), brain (103) and breast (84) cancers. Exploring the function of H3.3 in PC, we found that expression of two members of H3.3 chaperone complex HIRA (HIRA and UBN1), is increased in tumor compared with normal prostate tissue, is elevated in high/very high-risk PC groups (concordant with increased Gleason score), and their accumulation is associated with negative prognosis. We observed that HIRA depletion obliterates PC cell growth *in vitro* and *in vivo* and deregulates expression of androgen-induced genes, including those essential for metastatic progression. Depletion of HIRA reduces AR binding to AREs. In addition, we found that CRPC cells expressing gain-of-function AR-Vs are HIRA-dependent. Our emerging data strongly suggest an oncogenic function of HIRA-dependent H3.3 pathway in PC. Importantly, it was recently shown that HIRA accumulation induces breast cancer metastasis by an H3.3-dependent aggressive transcriptional reprogramming (84), thus emphasizing function of H3.3/ HIRA pathway in cancer versus normal conditions and providing additional scientific premise to our observations.

In summary, our data suggest that H3.3-enriched enhancer chromatin serves as a platform for H3K27Ac-mediated BRD4 recruitment, which interacts with and retains AR at enhancer AREs, resulting in AR-driven transcription reprogramming, thus identifying H3.3 function in nuclear receptors activity and placing HIRA complex at the epicenter of H3.3 pathways in PC initiation and/or progression.

## Acknowledgements

We thank Dr. Scott Dehm (University of Minnesota Masonic Cancer Center) for generous gift of R1-AD1 and R1-D567 cell lines and Dr. Peter Adams (Sanford Burnham Prebys Medical Discovery Institute) for HIRA antibody. We thank our colleagues in the Ishov and Daaka laboratories. We thank Drs. Jorg Bungert and Michael Kladde (Department of Biochemistry, University of Florida College of Medicine) for extensive discussions. Work in the Ishov laboratory is supported by the NIH (R01DE026707 and R21CA198820) and UF Cancer Center Pilot & Exploratory Studies Award. S.S. is partially supported by Fulbright Scholar Grant (IIE, USEFP, and ECA). L.Z. supported by GM106174. Work in the Licht laboratory is supported by 1R01CA266078, a Leukemia and Lymphoma Society Specialized Center of Research, Florida Department of Health (22L03). Analysis and visualization of NGS data was performed by the University of Florida ICBR Bioinformatics Core (RRID: SCR_019120).

## Data availability

The datasets generated and analyzed during the current study are available in the NCBI Gene Expression Omnibus repository in superseries GSE230744, which includes the following three series: GSE230529 (ATAC-seq), GSE230741 (RNA-seq), GSE230740 (ChIP-seq).

## Authors Contribution

V.M.M. and A.M.I. designed the study; V.M.M. performed experiments with help from S.S., W.J.K., J.L; A.R. and L.Z. performed bioinformatic data analysis, V.M.M. and A.M.I. wrote the manuscript with the critical input from J.D.L. and Y.D.; A.M.I. supervised the study.

## Literature

1. Siegel, R.L., Miller, K.D., Fuchs, H.E. and Jemal, A. (2021) Cancer Statistics, 2021. CA Cancer J Clin, 71, 7–33.

2. Egan, A., Dong, Y., Zhang, H., Qi, Y., Balk, S.P. and Sartor, O. (2014) Castration-resistant prostate cancer: adaptive responses in the androgen axis. Cancer Treat Rev, 40, 426–433.

3. Rathkopf, D. and Scher, H.I. (2013) Androgen receptor antagonists in castration-resistant prostate cancer. Cancer J, 19, 43–49.

4. Akamatsu, S., Inoue, T., Ogawa, O. and Gleave, M.E. (2018) Clinical and molecular features of treatment-related neuroendocrine prostate cancer. Int J Urol, 25, 345–351.

5. Formaggio, N., Rubin, M.A. and Theurillat, J.P. (2021) Loss and revival of androgen receptor signaling in advanced prostate cancer. Oncogene, 40, 1205–1216.

6. Dehm, S.M. and Tindall, D.J. (2011) Alternatively spliced androgen receptor variants. Endocr Relat Cancer, 18, R183–196.

7. Attard, G., Richards, J. and de Bono, J.S. (2011) New strategies in metastatic prostate cancer: targeting the androgen receptor signaling pathway. Clin Cancer Res, 17, 1649–1657.

8. Henikoff, S. (2008) Nucleosome destabilization in the epigenetic regulation of gene expression. Nat Rev Genet, 9, 15–26.

9. Hake, S.B. and Allis, C.D. (2006) Histone H3 variants and their potential role in indexing mammalian genomes: the "H3 barcode hypothesis". Proc Natl Acad Sci U S A, 103, 6428–6435.

10. Verreault, A., Kaufman, P.D., Kobayashi, R. and Stillman, B. (1996) Nucleosome assembly by a complex of CAF-1 and acetylated histones H3/H4. Cell, 87, 95–104.

11. Szenker, E., Ray-Gallet, D. and Almouzni, G. (2011) The double face of the histone variant H3.3. Cell Res, 21, 421–434.

12. Ahmad, K. and Henikoff, S. (2002) Histone H3 variants specify modes of chromatin assembly. Proc Natl Acad Sci U S A, 99 Suppl 4, 16477–16484.

13. Tagami, H., Ray-Gallet, D., Almouzni, G. and Nakatani, Y. (2004) Histone H3.1 and H3.3 complexes mediate nucleosome assembly pathways dependent or independent of DNA synthesis. Cell, 116, 51–61.

14. Goldberg, A.D., Banaszynski, L.A., Noh, K.M., Lewis, P.W., Elsaesser, S.J., Stadler, S., Dewell, S., Law, M., Guo, X., Li, X. et al. (2010) Distinct factors control histone variant H3.3 localization at specific genomic regions. Cell, 140, 678–691.

15. Lewis, P.W., Elsaesser, S.J., Noh, K.M., Stadler, S.C. and Allis, C.D. (2010) Daxx is an H3.3-specific histone chaperone and cooperates with ATRX in replication-independent chromatin assembly at telomeres. Proc Natl Acad Sci U S A, 107, 14075–14080.

16. Drane, P., Ouararhni, K., Depaux, A., Shuaib, M. and Hamiche, A. (2010) The death-associated protein DAXX is a novel histone chaperone involved in the replication-independent deposition of H3.3. Genes Dev, 24, 1253–1265.

17. Wong, L.H., McGhie, J.D., Sim, M., Anderson, M.A., Ahn, S., Hannan, R.D., George, A.J., Morgan, K.A., Mann, J.R. and Choo, K.H. (2010) ATRX interacts with H3.3 in maintaining telomere structural integrity in pluripotent embryonic stem cells. Genome Res, 20, 351–360.

18. Jin, C.Y., Zang, C.Z., Wei, G., Cui, K.R., Peng, W.Q., Zhao, K.J. and Felsenfeld, G. (2009) H3.3/H2A.Z double variant-containing nucleosomes mark ’nucleosome-free regions’ of active promoters and other regulatory regions. Nat Genet, 41, 941–U112.

19. Jin, C. and Felsenfeld, G. (2007) Nucleosome stability mediated by histone variants H3.3 and H2A.Z. Genes Dev, 21, 1519–1529.

20. Hake, S.B., Garcia, B.A., Duncan, E.M., Kauer, M., Dellaire, G., Shabanowitz, J., Bazett-Jones, D.P., Allis, C.D. and Hunt, D.F. (2006) Expression patterns and post-translational modifications associated with mammalian histone H3 variants. J Biol Chem, 281, 559–568.

21. McKittrick, E., Gafken, P.R., Ahmad, K. and Henikoff, S. (2004) Histone H3.3 is enriched in covalent modifications associated with active chromatin. Proc Natl Acad Sci U S A, 101, 1525–1530.

22. Siegel, T.N., Hekstra, D.R., Kemp, L.E., Figueiredo, L.M., Lowell, J.E., Fenyo, D., Wang, X., Dewell, S. and Cross, G.A. (2009) Four histone variants mark the boundaries of polycistronic transcription units in Trypanosoma brucei. Genes Dev, 23, 1063–1076.

23. Talbert, P.B. and Henikoff, S. (2009) Chromatin-based transcriptional punctuation. Genes Dev, 23, 1037–1041.

24. Dyer, M.A., Qadeer, Z.A., Valle-Garcia, D. and Bernstein, E. (2017) ATRX and DAXX: Mechanisms and Mutations. Cold Spring Harb Perspect Med, 7.

25. Buschbeck, M. and Hake, S.B. (2017) Variants of core histones and their roles in cell fate decisions, development and cancer. Nat Rev Mol Cell Bio, 18, 299–314.

26. Ilter, D., Blenis, J. and Gomes, A.P. (2020) Histone H3 variants at the root of metastasis. Mol Cell Oncol, 7, 1684128.

27. Ghiraldini, F.G., Filipescu, D. and Bernstein, E. (2021) Solid tumours hijack the histone variant network. Nat Rev Cancer.

28. Chen, P., Zhao, J., Wang, Y., Wang, M., Long, H., Liang, D., Huang, L., Wen, Z., Li, W., Li, X. et al. (2013) H3.3 actively marks enhancers and primes gene transcription via opening higher-ordered chromatin. Genes Dev, 27, 2109–2124.

29. Martire, S., Gogate, A.A., Whitmill, A., Tafessu, A., Nguyen, J., Teng, Y.C., Tastemel, M. and Banaszynski, L.A. (2019) Phosphorylation of histone H3.3 at serine 31 promotes p300 activity and enhancer acetylation. Nat Genet, 51, 941-+.

30. Chang, F.T., Chan, F.L., JD, R.M., Udugama, M., Mayne, L., Collas, P., Mann, J.R. and Wong, L.H. (2015) CHK1-driven histone H3.3 serine 31 phosphorylation is important for chromatin maintenance and cell survival in human ALT cancer cells. Nucleic Acids Res, 43, 2603–2614.

31. Armache, A., Yang, S., Martinez de Paz, A., Robbins, L.E., Durmaz, C., Cheong, J.Q., Ravishankar, A., Daman, A.W., Ahimovic, D.J., Klevorn, T., et al. (2020) Histone H3.3 phosphorylation amplifies stimulation-induced transcription. Nature, 583, 852–857.

32. Rasool, R.U., Natesan, R., Deng, Q., Aras, S., Lal, P., Sander Effron, S., Mitchell-Velasquez, E., Posimo, J.M., Carskadon, S., Baca, S.C. et al. (2019) CDK7 Inhibition Suppresses Castration-Resistant Prostate Cancer through MED1 Inactivation. Cancer Discov, 9, 1538–1555.

33. Nyquist, M.D., Li, Y., Hwang, T.H., Manlove, L.S., Vessella, R.L., Silverstein, K.A., Voytas, D.F. and Dehm, S.M. (2013) TALEN-engineered AR gene rearrangements reveal endocrine uncoupling of androgen receptor in prostate cancer. Proc Natl Acad Sci U S A, 110, 17492–17497.

34. Ye, X., Zerlanko, B., Kennedy, A., Banumathy, G., Zhang, R. and Adams, P.D. (2007) Downregulation of Wnt signaling is a trigger for formation of facultative heterochromatin and onset of cell senescence in primary human cells. Mol Cell, 27, 183–196.

35. Ishov, A.M., Vladimirova, O.V. and Maul, G.G. (2002) Daxx-mediated accumulation of human cytomegalovirus tegument protein pp71 at ND10 facilitates initiation of viral infection at these nuclear domains. J Virol, 76, 7705–7712.

36. Johnson, D.S., Mortazavi, A., Myers, R.M. and Wold, B. (2007) Genome-wide mapping of in vivo protein-DNA interactions. Science, 316, 1497–1502.

37. Dobin, A., Davis, C.A., Schlesinger, F., Drenkow, J., Zaleski, C., Jha, S., Batut, P., Chaisson, M. and Gingeras, T.R. (2013) STAR: ultrafast universal RNA-seq aligner. Bioinformatics, 29, 15–21.

38. Love, M.I., Huber, W. and Anders, S. (2014) Moderated estimation of fold change and dispersion for RNA-seq data with DESeq2. Genome Biol, 15, 550.

39. Langmead, B. and Salzberg, S.L. (2012) Fast gapped-read alignment with Bowtie 2. Nat Methods, 9, 357–359.

40. Feng, J., Liu, T., Qin, B., Zhang, Y. and Liu, X.S. (2012) Identifying ChIP-seq enrichment using MACS. Nat Protoc, 7, 1728–1740.

41. Whyte, W.A., Orlando, D.A., Hnisz, D., Abraham, B.J., Lin, C.Y., Kagey, M.H., Rahl, P.B., Lee, T.I. and Young, R.A. (2013) Master transcription factors and mediator establish super-enhancers at key cell identity genes. Cell, 153, 307–319.

42. Sarwar, S., Morozov, V.M., Purayil, H., Daaka, Y. and Ishov, A.M. (2022) Inhibition of Mps1 kinase enhances taxanes efficacy in castration resistant prostate cancer. Cell Death Dis, 13, 868.

43. Giovinazzi, S., Lindsay, C.R., Morozov, V.M., Escobar-Cabrera, E., Summers, M.K., Han, H.S., McIntosh, L.P. and Ishov, A.M. (2012) Regulation of mitosis and taxane response by Daxx and Rassf1. Oncogene, 31, 13–26.

44. Morozov, V.M., Giovinazzi, S. and Ishov, A.M. (2017) CENP-B protects centromere chromatin integrity by facilitating histone deposition via the H3.3-specific chaperone Daxx. Epigenetics Chromatin, 10, 63.

45. Yan, Y. and Huang, H. (2019) Interplay Among PI3K/AKT, PTEN/FOXO and AR Signaling in Prostate Cancer. Adv Exp Med Biol, 1210, 319–331.

46. Michalaki, V., Syrigos, K., Charles, P. and Waxman, J. (2004) Serum levels of IL-6 and TNF-alpha correlate with clinicopathological features and patient survival in patients with prostate cancer. Br J Cancer, 90, 2312–2316.

47. Cao, Z. and Kyprianou, N. (2015) Mechanisms navigating the TGF-beta pathway in prostate cancer. Asian J Urol, 2, 11–18.

48. Rodriguez-Berriguete, G., Fraile, B., Martinez-Onsurbe, P., Olmedilla, G., Paniagua, R. and Royuela, M. (2012) MAP Kinases and Prostate Cancer. J Signal Transduct, 2012, 169170.

49. Murillo-Garzon, V. and Kypta, R. (2017) WNT signalling in prostate cancer. Nat Rev Urol, 14, 683–696.

50. Schwartz, B.E. and Ahmad, K. (2005) Transcriptional activation triggers deposition and removal of the histone variant H3.3. Genes Dev, 19, 804–814.

51. Wirbelauer, C., Bell, O. and Schubeler, D. (2005) Variant histone H3.3 is deposited at sites of nucleosomal displacement throughout transcribed genes while active histone modifications show a promoter-proximal bias. Genes Dev, 19, 1761–1766.

52. Mito, Y., Henikoff, J.G. and Henikoff, S. (2005) Genome-scale profiling of histone H3.3 replacement patterns. Nat Genet, 37, 1090–1097.

53. Tafessu, A., O’Hara, R., Martire, S., Dube, A.L., Saha, P., Gant, V.U. and Banaszynski, L.A. (2023) H3.3 contributes to chromatin accessibility and transcription factor binding at promoter-proximal regulatory elements in embryonic stem cells. Genome Biol, 24, 25.

54. Wang, H., Walsh, S.T. and Parthun, M.R. (2008) Expanded binding specificity of the human histone chaperone NASP. Nucleic Acids Res, 36, 5763–5772.

55. Morozov, V.M., Gavrilova, E.V., Ogryzko, V.V. and Ishov, A.M. (2012) Dualistic function of Daxx at centromeric and pericentromeric heterochromatin in normal and stress conditions. Nucleus, 3.

56. Li, H., Leo, C., Zhu, J., Wu, X., O’Neil, J., Park, E.J. and Chen, J.D. (2000) Sequestration and inhibition of Daxx-mediated transcriptional repression by PML. Mol Cell Biol, 20, 1784–1796.

57. Hollenbach, A.D., McPherson, C.J., Mientjes, E.J., Iyengar, R. and Grosveld, G. (2002) Daxx and histone deacetylase II associate with chromatin through an interaction with core histones and the chromatin-associated protein Dek. J Cell Sci, 115, 3319–3330.

58. Lin, D.Y., Fang, H.I., Ma, A.H., Huang, Y.S., Pu, Y.S., Jenster, G., Kung, H.J. and Shih, H.M. (2004) Negative modulation of androgen receptor transcriptional activity by Daxx. Mol Cell Biol, 24, 10529–10541.

59. Tafessu, A. and Banaszynski, L.A. (2020) Establishment and function of chromatin modification at enhancers. Open Biol, 10, 200255.

60. Herz, H.M., Garruss, A. and Shilatifard, A. (2013) SET for life: biochemical activities and biological functions of SET domain-containing proteins. Trends Biochem Sci, 38, 621–639.

61. Siddaway, R., Milos, S., Coyaud, E., Yun, H.Y., Morcos, S.M., Pajovic, S., Campos, E.I., Raught, B. and Hawkins, C. (2022) The in vivo Interaction Landscape of Histones H3.1 and H3.3. Mol Cell Proteomics, 21, 100411.

62. Jahangiri, L., Tsaprouni, L., Trigg, R.M., Williams, J.A., Gkoutos, G.V., Turner, S.D. and Pereira, J. (2020) Core regulatory circuitries in defining cancer cell identity across the malignant spectrum. Open Biol, 10, 200121.

63. Rahnamoun, H., Lee, J., Sun, Z., Lu, H., Ramsey, K.M., Komives, E.A. and Lauberth, S.M. (2018) RNAs interact with BRD4 to promote enhanced chromatin engagement and transcription activation. Nat Struct Mol Biol, 25, 687–697.

64. Asangani, I.A., Dommeti, V.L., Wang, X., Malik, R., Cieslik, M., Yang, R., Escara-Wilke, J., Wilder-Romans, K., Dhanireddy, S., Engelke, C. et al. (2014) Therapeutic targeting of BET bromodomain proteins in castration-resistant prostate cancer. Nature, 510, 278–282.

65. Zaret, K.S. (2020) Pioneer Transcription Factors Initiating Gene Network Changes. Annu Rev Genet, 54, 367–385.

66. Maeda, K., Habara, M., Kawaguchi, M., Matsumoto, H., Hanaki, S., Masaki, T., Sato, Y., Matsuyama, H., Kunieda, K., Nakagawa, H. et al. (2022) FKBP51 and FKBP52 regulate androgen receptor dimerization and proliferation in prostate cancer cells. Mol Oncol, 16, 940–956.

67. Magee, J.A., Chang, L.W., Stormo, G.D. and Milbrandt, J. (2006) Direct, androgen receptor-mediated regulation of the FKBP5 gene via a distal enhancer element. Endocrinology, 147, 590–598.

68. Makkonen, H., Kauhanen, M., Paakinaho, V., Jaaskelainen, T. and Palvimo, J.J. (2009) Long-range activation of FKBP51 transcription by the androgen receptor via distal intronic enhancers. Nucleic Acids Res, 37, 4135–4148.

69. Jaaskelainen, T., Makkonen, H. and Palvimo, J.J. (2011) Steroid up-regulation of FKBP51 and its role in hormone signaling. Curr Opin Pharmacol, 11, 326–331.

70. Paakinaho, V., Makkonen, H., Jaaskelainen, T. and Palvimo, J.J. (2010) Glucocorticoid receptor activates poised FKBP51 locus through long-distance interactions. Mol Endocrinol, 24, 511–525.

71. Loven, J., Hoke, H.A., Lin, C.Y., Lau, A., Orlando, D.A., Vakoc, C.R., Bradner, J.E., Lee, T.I. and Young, R.A. (2013) Selective inhibition of tumor oncogenes by disruption of super-enhancers. Cell, 153, 320–334.

72. Kumari, S., Senapati, D. and Heemers, H.V. (2017) Rationale for the development of alternative forms of androgen deprivation therapy. Endocr Relat Cancer, 24, R275–R295.

73. Cirillo, L.A., Lin, F.R., Cuesta, I., Friedman, D., Jarnik, M. and Zaret, K.S. (2002) Opening of compacted chromatin by early developmental transcription factors HNF3 (FoxA) and GATA-4. Mol Cell, 9, 279–289.

74. Yu, X., Gupta, A., Wang, Y., Suzuki, K., Mirosevich, J., Orgebin-Crist, M.C. and Matusik, R.J. (2005) Foxa1 and Foxa2 interact with the androgen receptor to regulate prostate and epididymal genes differentially. Ann N Y Acad Sci, 1061, 77–93.

75. Jin, H.J., Zhao, J.C., Wu, L., Kim, J. and Yu, J. (2014) Cooperativity and equilibrium with FOXA1 define the androgen receptor transcriptional program. Nat Commun, 5, 3972.

76. Dixon, J.R., Selvaraj, S., Yue, F., Kim, A., Li, Y., Shen, Y., Hu, M., Liu, J.S. and Ren, B. (2012) Topological domains in mammalian genomes identified by analysis of chromatin interactions. Nature, 485, 376–380.

77. Phillips-Cremins, J.E., Sauria, M.E., Sanyal, A., Gerasimova, T.I., Lajoie, B.R., Bell, J.S., Ong, C.T., Hookway, T.A., Guo, C., Sun, Y. et al. (2013) Architectural protein subclasses shape 3D organization of genomes during lineage commitment. Cell, 153, 1281–1295.

78. Claessens, F., Joniau, S. and Helsen, C. (2017) Comparing the rules of engagement of androgen and glucocorticoid receptors. Cell Mol Life Sci, 74, 2217–2228.

79. Arora, V.K., Schenkein, E., Murali, R., Subudhi, S.K., Wongvipat, J., Balbas, M.D., Shah, N., Cai, L., Efstathiou, E., Logothetis, C. et al. (2013) Glucocorticoid receptor confers resistance to antiandrogens by bypassing androgen receptor blockade. Cell, 155, 1309–1322.

80. Xie, N., Cheng, H., Lin, D., Liu, L., Yang, O., Jia, L., Fazli, L., Gleave, M.E., Wang, Y., Rennie, P. et al. (2015) The expression of glucocorticoid receptor is negatively regulated by active androgen receptor signaling in prostate tumors. Int J Cancer, 136, E27–38.

81. Gottlieb, B., Beitel, L.K., Nadarajah, A., Paliouras, M. and Trifiro, M. (2012) The androgen receptor gene mutations database: 2012 update. Hum Mutat, 33, 887–894.

82. Shafi, A.A., Cox, M.B. and Weigel, N.L. (2013) Androgen receptor splice variants are resistant to inhibitors of Hsp90 and FKBP52, which alter androgen receptor activity and expression. Steroids, 78, 548–554.

83. Hornberg, E., Ylitalo, E.B., Crnalic, S., Antti, H., Stattin, P., Widmark, A., Bergh, A. and Wikstrom, P. (2011) Expression of androgen receptor splice variants in prostate cancer bone metastases is associated with castration-resistance and short survival. PLoS One, 6, e19059.

84. Gomes, A.P., Ilter, D., Low, V., Rosenzweig, A., Shen, Z.J., Schild, T., Rivas, M.A., Er, E.E., McNally, D.R., Mutvei, A.P. et al. (2019) Dynamic Incorporation of Histone H3 Variants into Chromatin Is Essential for Acquisition of Aggressive Traits and Metastatic Colonization. Cancer Cell, 36, 402–417 e413.

85. Blessing, A.M., Ganesan, S., Rajapakshe, K., Ying Sung, Y., Reddy Bollu, L., Shi, Y., Cheung, E., Coarfa, C., Chang, J.T., McDonnell, D.P. et al. (2015) Identification of a Novel Coregulator, SH3YL1, That Interacts With the Androgen Receptor N-Terminus. Mol Endocrinol, 29, 1426–1439.

86. Tan, Y., Wang, L., Du, Y., Liu, X., Chen, Z., Weng, X., Guo, J., Chen, H., Wang, M. and Wang, X. (2018) Inhibition of BRD4 suppresses tumor growth in prostate cancer via the enhancement of FOXO1 expression. Int J Oncol, 53, 2503–2517.

87. Debes, J.D., Sebo, T.J., Lohse, C.M., Murphy, L.M., Haugen, D.A. and Tindall, D.J. (2003) p300 in prostate cancer proliferation and progression. Cancer Res, 63, 7638–7640.

88. Comuzzi, B., Nemes, C., Schmidt, S., Jasarevic, Z., Lodde, M., Pycha, A., Bartsch, G., Offner, F., Culig, Z. and Hobisch, A. (2004) The androgen receptor co-activator CBP is up-regulated following androgen withdrawal and is highly expressed in advanced prostate cancer. J Pathol, 204, 159–166.

89. Lasko, L.M., Jakob, C.G., Edalji, R.P., Qiu, W., Montgomery, D., Digiammarino, E.L., Hansen, T.M., Risi, R.M., Frey, R., Manaves, V. et al. (2017) Discovery of a selective catalytic p300/CBP inhibitor that targets lineage-specific tumours. Nature, 550, 128–132.

90. Jin, L., Garcia, J., Chan, E., de la Cruz, C., Segal, E., Merchant, M., Kharbanda, S., Raisner, R., Haverty, P.M., Modrusan, Z. et al. (2017) Therapeutic Targeting of the CBP/p300 Bromodomain Blocks the Growth of Castration-Resistant Prostate Cancer. Cancer Res, 77, 5564–5575.

91. Welti, J., Sharp, A., Brooks, N., Yuan, W., McNair, C., Chand, S.N., Pal, A., Figueiredo, I., Riisnaes, R., Gurel, B. et al. (2021) Targeting the p300/CBP Axis in Lethal Prostate Cancer. Cancer Discov, 11, 1118–1137.

92. Bradbury, R.H., Callis, R., Carr, G.R., Chen, H., Clark, E., Feron, L., Glossop, S., Graham, M.A., Hattersley, M., Jones, C. et al. (2016) Optimization of a Series of Bivalent Triazolopyridazine Based Bromodomain and Extraterminal Inhibitors: The Discovery of (3R)-4-[2-[4-[1-(3-Methoxy-[1,2,4]triazolo[4,3-b]pyridazin-6-yl)-4-piperidyl]phen oxy]ethyl]-1,3-dimethyl-piperazin-2-one (AZD5153). J Med Chem, 59, 7801–7817.

93. Booth, L., Roberts, J., Poklepovic, A. and Dent, P. (2018) The CHK1 inhibitor SRA737 synergizes with PARP1 inhibitors to kill carcinoma cells. Cancer Biol Ther, 19, 786–796.

94. Tang, S.C., Vijayakumar, U., Zhang, Y. and Fullwood, M.J. (2022) Super-Enhancers, Phase-Separated Condensates, and 3D Genome Organization in Cancer. Cancers (Basel), 14.

95. Sanyal, A., Lajoie, B.R., Jain, G. and Dekker, J. (2012) The long-range interaction landscape of gene promoters. Nature, 489, 109–113.

96. Ong, C.T. and Corces, V.G. (2011) Enhancer function: new insights into the regulation of tissue-specific gene expression. Nat Rev Genet, 12, 283–293.

97. Sabari, B.R., Dall’Agnese, A., Boija, A., Klein, I.A., Coffey, E.L., Shrinivas, K., Abraham, B.J., Hannett, N.M., Zamudio, A.V., Manteiga, J.C. et al. (2018) Coactivator condensation at super-enhancers links phase separation and gene control. Science, 361.

98. Ishov, A.M., Gurumurthy, A. and Bungert, J. (2020) Coordination of transcription, processing, and export of highly expressed RNAs by distinct biomolecular condensates. Emerg Top Life Sci, 4, 281–291.

99. Patel, M.C., Debrosse, M., Smith, M., Dey, A., Huynh, W., Sarai, N., Heightman, T.D., Tamura, T. and Ozato, K. (2013) BRD4 coordinates recruitment of pause release factor P-TEFb and the pausing complex NELF/DSIF to regulate transcription elongation of interferon-stimulated genes. Mol Cell Biol, 33, 2497–2507.

100. Donati, B., Lorenzini, E. and Ciarrocchi, A. (2018) BRD4 and Cancer: going beyond transcriptional regulation. Mol Cancer, 17, 164.

101. Forsyth, R.G., Krenacs, T., Athanasou, N. and Hogendoorn, P.C.W. (2021) Cell Biology of Giant Cell Tumour of Bone: Crosstalk between m/wt Nucleosome H3.3, Telomeres and Osteoclastogenesis. Cancers (Basel), 13.

102. Jiao, Y., Shi, C., Edil, B.H., de Wilde, R.F., Klimstra, D.S., Maitra, A., Schulick, R.D., Tang, L.H., Wolfgang, C.L., Choti, M.A. et al. (2011) DAXX/ATRX, MEN1, and mTOR pathway genes are frequently altered in pancreatic neuroendocrine tumors. Science, 331, 1199–1203.

103. Schwartzentruber, J., Korshunov, A., Liu, X.Y., Jones, D.T., Pfaff, E., Jacob, K., Sturm, D., Fontebasso, A.M., Quang, D.A., Tonjes, M. et al. (2012) Driver mutations in histone H3.3 and chromatin remodelling genes in paediatric glioblastoma. Nature, 482, 226–231.

