## Supplementary figure legends for "HIRA-mediated loading of histone variant H3.3 controls androgen-induced transcription by regulation of AR/BRD4 complex assembly at enhancers"

**Fig. S1. Production and characterization of R1-AD1 (AR-WT) FLAG/HA-H3.3 knock-in and knockout cells. A:** In R1-AD1 cells, CRISPR/Cas9 strategy was used to introduce FLAG/HA tag at 5’ end of H3F3A gene by homologous recombination. Single-cell clones were characterized for expression of HA. Microscopy analysis of endogenous tagged FLAG/HA-H3.3 in exponentially grown cells. HA (H3.3): green. DNA: blue. Arrow: mitotic cell confirming chromosome association of FLAG/HA-H3.3. **B:** CRISPR/Cas9 strategy was used to knockout (KO) AR, HIRA, and Daxx in FLAG/HA-H3.3 tagged R1-AD1 cells. Western analysis to confirm KO. Modification of AR, HIRA or Daxx does not affect protein levels of remaining two proteins; HIRA KO reduced levels of H3.3. H3: total histone H3; actin: loading control. **C:** R1-AD1 cells were androgen-deprived (72h) and androgen R1881 stimulated (4h, 12h, 24h); levels of BRD4, H3.3 (HA) and total H3 are shown; actin: loading control.

**Fig. S2. Expression analysis (RNA-seq) of R1-AD1 cells, parental (WT), AR KO, Daxx KO and HIRA KO. A: Analysis of genes deregulated by AR KO, Daxx KO, HIRA KO.** Venn diagram of genes that are at least two-folds (p-adjusted <0.05) up- and downregulated by 4h of R1881 stimulation (compared with androgen-deprived (72h)) in parental R1-AD1 and in KO cells. **B:** **Analysis of genes that are deregulated in AR KO and HIRA KO.** Top, Venn diagram; androgen-regulated genes that are affected in AR KO or HIRA KO cells; same parameters as in **A**. Bottom: Heatmaps showing expression of androgen-regulated genes that are affected by both AR KO and HIRA KO identified by KEGG pathway enrichment analysis. Visualization by “pheatmap”, http://cran.nexr.com/web/packages/pheatmap/index.html.

**Fig. S3. H3.3 association with TSS of androgen-regulated genes; analysis within time points (complementary to Fig. 3).** ChIP-seq profiles of H3.3 at TSS of 409 androgen-up- (**left**) and 328 androgen-downregulated (**right**) genes in R1-AD1 parental (Control), AR KO, Daxx KO and HIRA KO cells in the androgen-deprived (0h) and androgen-induced conditions (at 4h, 12h, 24h). 0: TSS.

**Fig. S4. H3.3 association with androgen-regulated genes; analysis within cell lines.** ChIP-seq profiles of H3.3 at the 409 androgen-up- (**left**) and 328 androgen-downregulated (**right**) genes in R1-AD1 parental (Control), AR KO, Daxx KO and HIRA KO cells in the androgen-deprived (0h) and androgen-induced conditions (at 4h, 12h, 24h). Transcription start sites: TSS; transcription end sites: TES.

**Fig. S5. H3K27Ac association with TSS of androgen-regulated genes; analysis within time points (complementary to Fig. 4).** ChIP-seq profiles of H3K27Ac at TSS of 409 androgen-up- (**left**) and 328 androgen-downregulated (**right**) genes in R1-AD1 parental (Control), AR KO, Daxx KO and HIRA KO cells in the androgen-deprived (0h) and androgen-induced conditions (at 4h, 12h, 24h). 0: TSS.

**Fig. S6. H3K27Ac association with androgen-regulated genes; analysis within cell lines.** ChIP-seq profiles of H3K27Ac at the 409 androgen-up- (**left**) and 328 androgen-downregulated (**right**) genes in R1-AD1 parental (Control), AR KO, Daxx KO and HIRA KO cells in the androgen-deprived (0h) and androgen-induced conditions (at 4h, 12h, 24h). Transcription start sites: TSS; transcription end sites: TES.

**Fig. S7. AR association with chromatin is regulated by H3.3 chaperone HIRA; analysis within time points (complementary to Fig. 5).** **A** **Left:** AR analysis of AR peaks (position “0”: AR at 4h in R1-AD1 parental cells) in R1-AD1 parental (Control, red), Daxx KO (blue), and HIRA KO (yellow), in androgen-deprived (72h) and R1881 stimulated for 4h, 12h, 24h. Analysis of AR at enhancers associated with 409 up- (**middle**) and 328 downregulated (**right**) genes. **B:** boxplots of total AR signal around AR peaks in enhancers associated with 409 up- and 328 downregulated genes.

**Fig. S8. H3K4me1 profiling at enhancers**. H3K4me1 at AR-positive enhancers nearest to 409 up- and 328 downregulated genes (top), all AR-positive enhancers (1980; middle left), AR-positive enhancers that are not associated with regulated genes (1489; middle right), all AR-negative enhancers (40644; bottom left) and all enhancers (42624; bottom right). Analysis in R1-AD1 parental (Control, red), AR KO (green), Daxx KO (blue) and HIRA KO (yellow) in androgen-deprived (72h) conditions.

**Fig. S9. Dynamics of H3.3 at enhancers; analysis within time points (complementary to Fig. 6).** **A:** Metaplot and **B:** boxplot of H3.3 at AR peaks (position “0”: AR at 4h in R1-AD1 parental cells) at enhancers associated with up- (left) and down-regulated (right) genes in R1-AD1 parental (Control, red), AR KO (green), Daxx KO (blue), and HIRA KO (yellow) in androgen-deprived (72h) and R1881 stimulated for 4h, 12h, 24h.

**Fig. S10. Dynamics of H3K27Ac at enhancers; analysis within time points (complementary to Fig. 7).** **A:** Metaplot and **B:** boxplot of H3K27Ac ChIP-seq analysis at AR peaks (position “0”: AR at 4h in R1-AD1 parental cells) at enhancers associated with up- (left) and down-regulated (right) genes in R1-AD1 parental (Control, red), AR KO (green), Daxx KO (blue), and HIRA KO (yellow) in androgen-deprived (72h) and R1881 stimulated for 4h, 12h, 24h.

**Fig. S11. Dynamics of H3.3 S31ph at enhancers/ SE; analysis within time points (complementary to Fig. 8).** **A:** Metaplot and **B:** boxplot of H3.3 S31ph ChIP-seq analysis at AR peaks (position “0”: AR at 4h in R1-AD1 parental cells) at enhancers associated with up- (left) and down-regulated (right) genes in R1-AD1 parental (Control, red), AR KO (green), Daxx KO (blue) and HIRA KO (yellow) in androgen-deprived (72h) and R1881 stimulated for 4h, 12h.

**Fig. S12. Dynamics of BRD4 at enhancers; analysis within time points (complementary to Fig. 9).** **A:** Metaplot and **B:** boxplot of BRD4 ChIP-seq analysis at AR peaks (position “0”: AR at 4h in R1-AD1 parental cells) at enhancers associated with up- (left) and down-regulated (right) genes in R1-AD1 parental (Control, red), AR KO (green), Daxx KO (blue), and HIRA KO (yellow) in androgen-deprived (72h) and R1881 stimulated for 4h, 12h.

**Fig. S13. DNA accessibility analyzed by ATAC-seq at enhancers; analysis within time points (complementary to Fig. 10).** Metaplot of ATAC-seq analysis at AR peaks (position “0”: AR at 4h in R1-AD1 parental cells) at enhancers associated with up- (left) and down-regulated (right) genes in R1-AD1 parental (Control, red), AR KO (green), Daxx KO (blue), and HIRA KO (yellow) in androgen-deprived (72h) and R1881 stimulated for 4h.

**Fig. S14. Examples of epigenetic profiles of enhancers/ SE associated with genes co-regulated by AR and HIRA.** R1-AD1 cells (parental, AR, Daxx, HIRA KO) were androgen-deprived for 72h (Control) and induced with 1 nM of R1881 for 4h, 12h, 24h. **A:** Androgen-induced expression of LONRF1, IL1R1, FKBP5 genes is reduced by AR and HIRA KO, and androgen-induced repression of ID2 gene is elevated by AR and HIRAKO (results of RNA-seq analysis). **B:** H3K4me1 (enhancer), H3K27Ac (active enhancer), H3.3, H3.3S31Ph, AR, BRD4, and CTCF profiling were analyzed by ChIP-seq. DNA accessibility analyzed by ATAC-seq. Arrows: transcription start; red lines outline SE regions.

**Fig. S15. DNA accessibility profiling at FKBP5 SE.** **Top:** Analysis of ATAC-seq at ARE-1 (1; half-site ARE, overlaps with FOXA1 binding site), ARE-2 and -3 (2 and 3; canonical AREs) at FKBP5 SE in R1-AD1 parental, AR KO, Daxx KO, HIRA KO at the androgen deprived (0h) and stimulated with 1nM of R1881 for 4h. Androgen stimulation for 4h induced ATAC peak at the ARE-1 (half-site) in parental, Daxx and HIRA KO cells. **Bottom:** Zoom-in comparison of ATAC-seq between parental and HIRA KO in androgen-deprived (72h) conditions. Bottom red bars: peaks increase in HIRA KO compared to parental cells. In androgen-deprived conditions, ARE-2 is more accessible in HIRA KO cells compared with parental cells. Vertical dashed lines: positions of AR binding.

**Fig. S16. Characterization of HIRA function in AR-V expressing cells. A. Intra-cellular localization of AR and AR-DLBD.** R1-AD1 (AR WT) and R1-D567 (AR-DLBD) cells androgen-deprived or treated with 1nM of R1881 for 2h, stained with anti-AR antibody (green); DNA (nuclei) blue. AR-WT accumulates in nuclei after androgen treatment, while AR-DLBD has nuclear localization in untreated and treated cells. **B. Characterization of FLAG/HA-H3.3 R1-D567 cells.** CRISPR/Cas9 strategy was used to tag endogenous H3.3 in R1-D567 cells using same strategy as in R1-AD1 cells. FLAG/HA was introduced at 5’ end of H3F3A gene by homologous recombination. Single-cell clones were characterized for expression of HA. Microscopy analysis of endogenous tagged FLAG/HA-H3.3 in exponentially grown cells. HA-H3.3: green. DNA: blue. Arrow: mitotic cells confirming chromosome association of FLAG/HA-H3.3. **C:** **KO and knock down (KD) characterization.** CRISPR/Cas9 strategy was used to KO AR and Daxx and shRNA to KD HIRA in R1-567 cells. Western analysis to confirm KO and KD. Modification of AR, HIRA or Daxx does not affect protein levels of remaining two proteins; HIRA KD reduced levels of H3.3, similarly to HIRA KO in R1-D1 cells. H3: total histone H3; actin: loading control. **D. AR status determines effect of HIRA depletion.** Representative images of colony formation assay with R1-AD1 (AR-WT) and R1-D567 (AR-DLBD) cells expressing control (CTR) or HIRA shRNA and transfected with HIRA siRNA. HIRA depletion reduces number of R1-AD1 colonies and eliminates most R1-D567 colonies. **E. H3.3 profiling at enhancers.** ChIP-seq analysis of H3.3 (endogenous HA-H3.3) at AR peaks (position “0”) at enhancers associated with 215 up- (left) and 185 down-regulated (right) genes in AR KO compared with R1-D567 parental cells. H3.3 plots in parental cells (Control, red), AR KO (green), Daxx KO (blue), and HIRA KD (yellow) in androgen-deprived (72h) conditions.

**Fig. S17. HIRA complex in PC. A:** HIRA and UBN1 expression (TCGA) in normal prostate tissue (gray) and prostate adenocarcinoma by Gleason score: 6 = Low, green; 7 = Intermediate, blue; 8-10 = High, red. HIRA and UBN1 expression are increased in PC compared with normal prostate and within Gleason groups. Numbers in legend: samples in group; p-values by Mann-Whitney U test. **B:** Disease free survival analysis based on HIRA expression normalized on GAPDH. Logrank p test (0.15) compares the survival distributions of high (magenta) and low (blue) expression groups. Hazard Rate (HR, the survive model calculated based on Cox proportional hazards model) >1 (1.9) indicates reduced survival of patients with high expression of HIRA. **C:** Kaplan-Meier plots for PC patients with high (magenta) and low (blue) levels of HIRA protein. High levels of HIRA are associated with negative survival prognoses; p: 0.09. Images: representative HIRA staining sections. Analysis at the Human Protein Atlas. **D:** Levels of HIRA in prostate by single cell RNA analysis. The glandular epithelial prostate cells, that are the main origin of prostate adenocarcinoma, have elevated expression of HIRA (compared with basal prostatic and urothelial cells); analysis at the Human Protein Atlas.

**Fig. S18. HIRA KO elevates cytotoxic effect of BRD4 inhibition. A:** R1-AD1 parental (CTL, blue), AR (red), Daxx (gray), and HIRA KO (yellow) cells were androgen-deprived for 72h, induced with 1 nM of R1881 and treated with corresponding concentrations of BRD4 inhibitor JQ1 (uM). AlamarBlue™ cell viability assay was performed 96h later. JQ1 IC50: parental and Daxx KO ~1 uM, AR KO ~0.3 uM, HIRA KO <0.1 uM. **B:** HIRA KO affects prostaspheres response to JQ1. Cells were set up in Matrigel, and prostaspheres were treated with 200 nM of JQ1 on day 11 and day 14 and documented at day 21 with Leica fluorescent microscope. Representative images of R1-AD1 parental, Daxx, AR, HIRA KO prostaspheres in control and JQ1 treated conditions. HIRA KO reduced growth of prostaspheres and elevates effect of JQ1 treatment. Bar: 150 mkm.

**Table S1. Androgen-dependent genes deregulated by AR and HIRA KO (complementary to Figs. 2, S2A).** List of 688 genes that are at least two-folds (p-adjusted <0.05) up- or down-regulated by R1881 stimulation in parental R1-AD1 cells and are affected in both AR KO and HIRA KO cells.

**Table S2. Pathways deregulation by HIRA and AR KO (complementary to Figs. 2, S2B).** List of androgen-regulated genes enriched in pathways identified with KEGG databases that are affected by both AR KO and HIRA KO.
