## Supplementary figures and images for "HIRA-mediated loading of histone variant H3.3 controls androgen-induced transcription by regulation of AR/BRD4 complex assembly at enhancers"

### Fig. S1

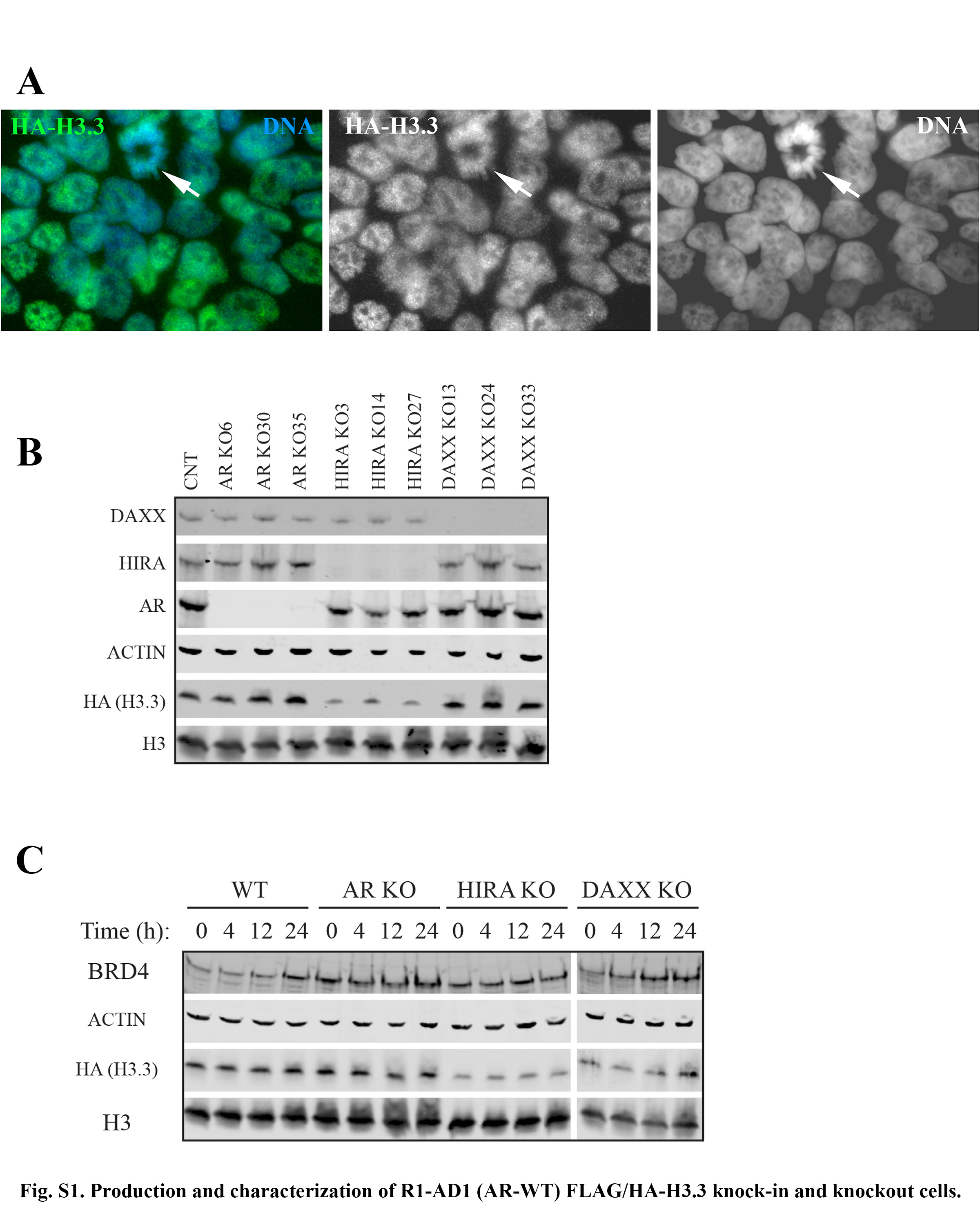

### Fig. S3

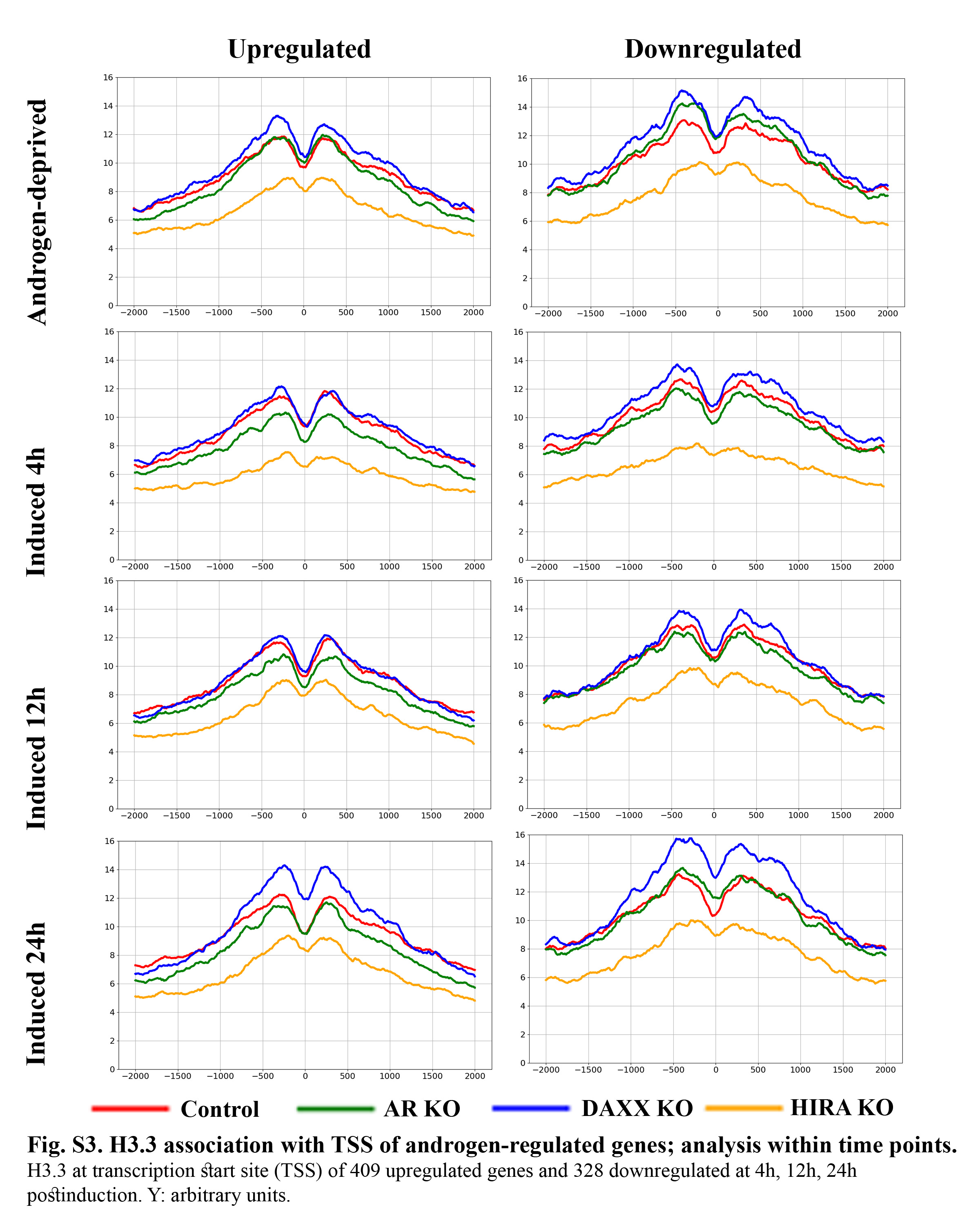

### Fig. S4

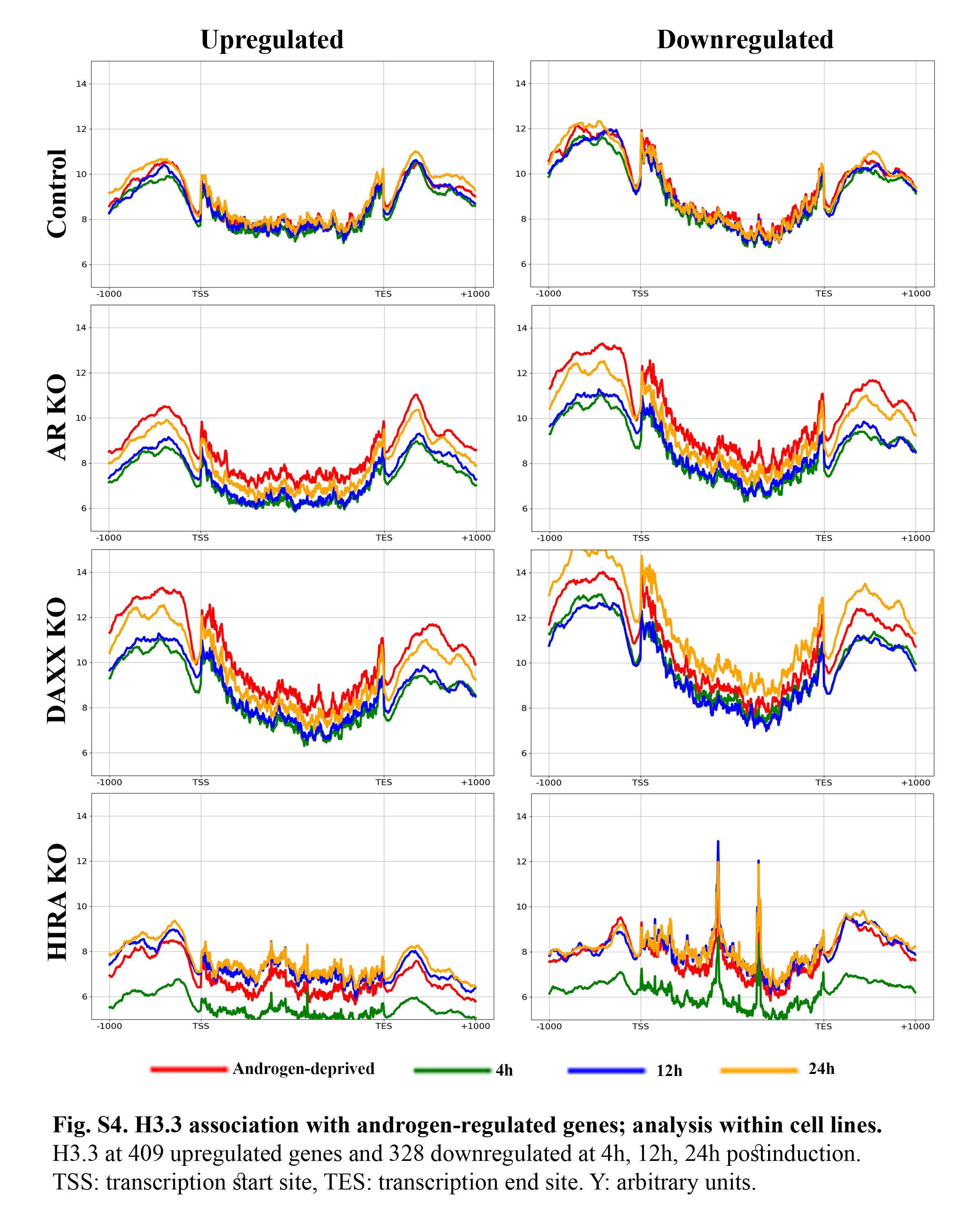

### Fig. S5

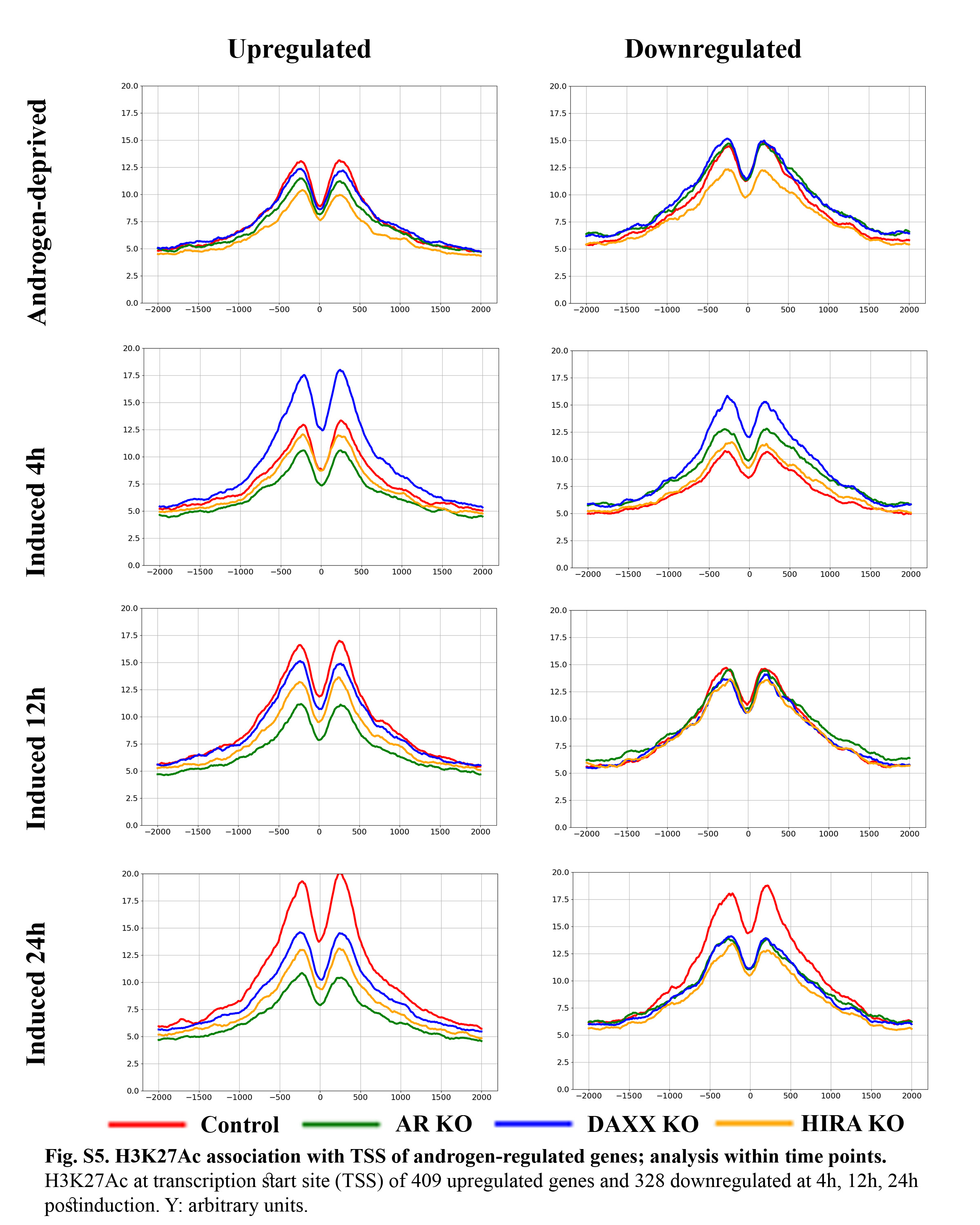

### Fig. S6

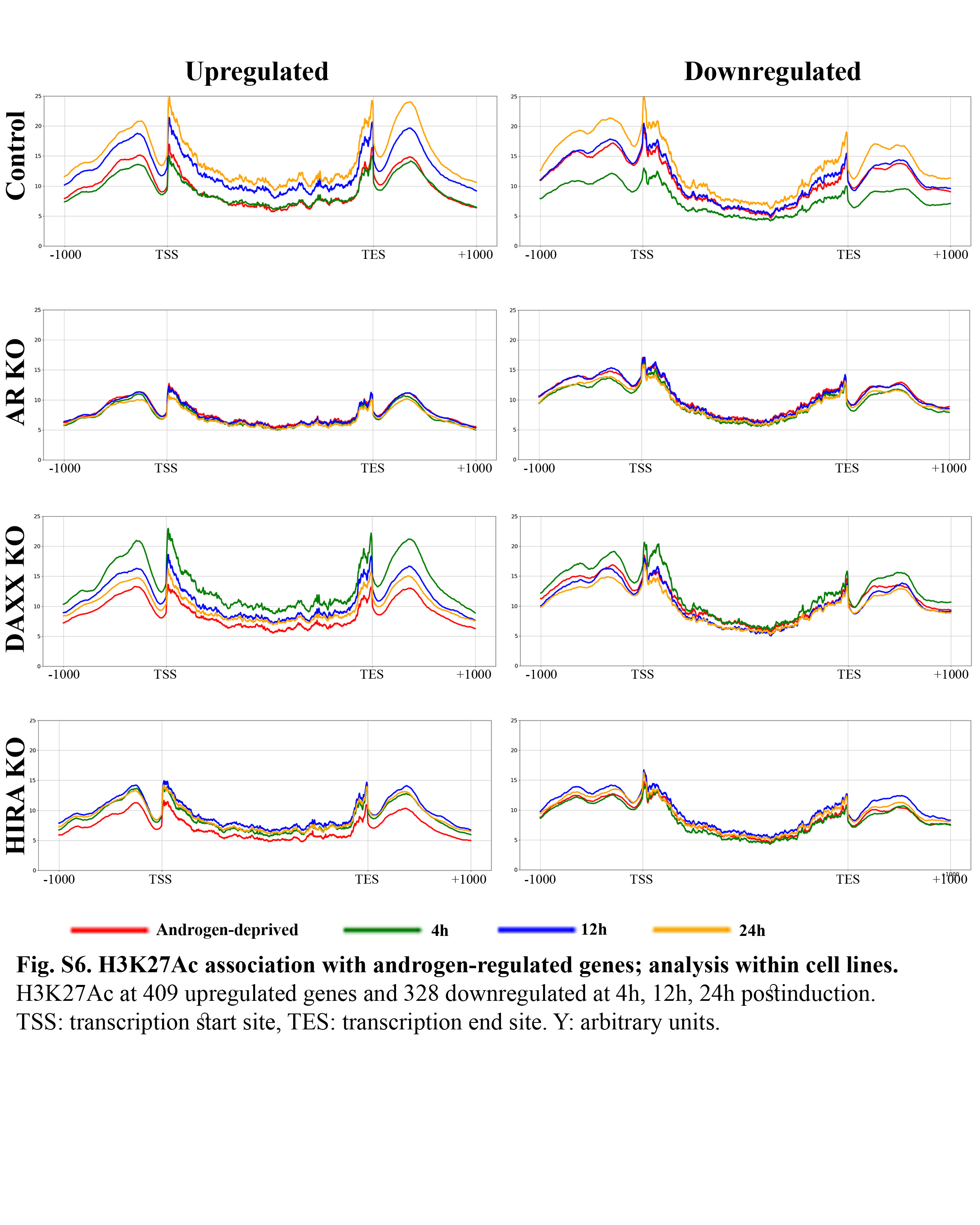

### Fig. S7A

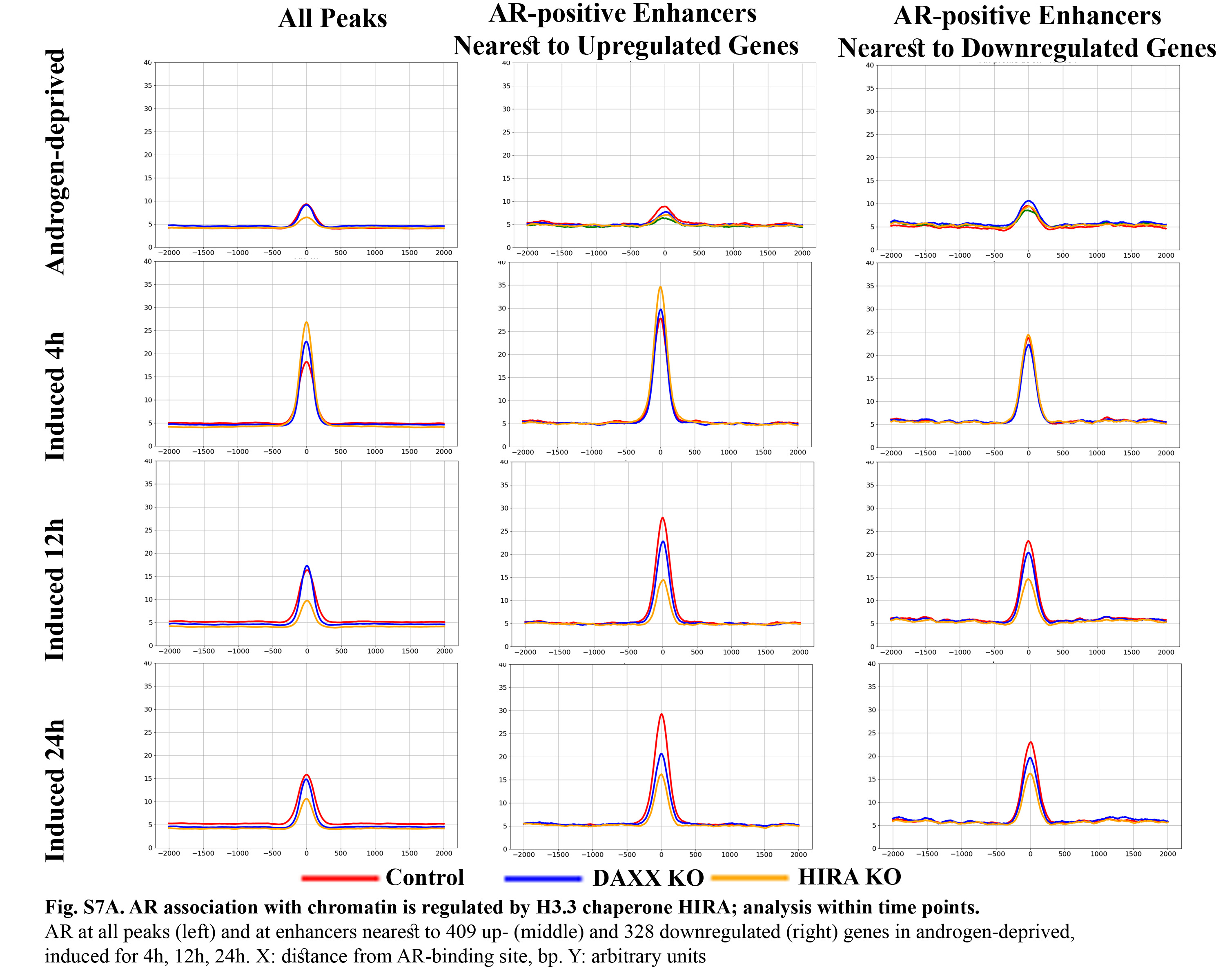

### Fig. S7B

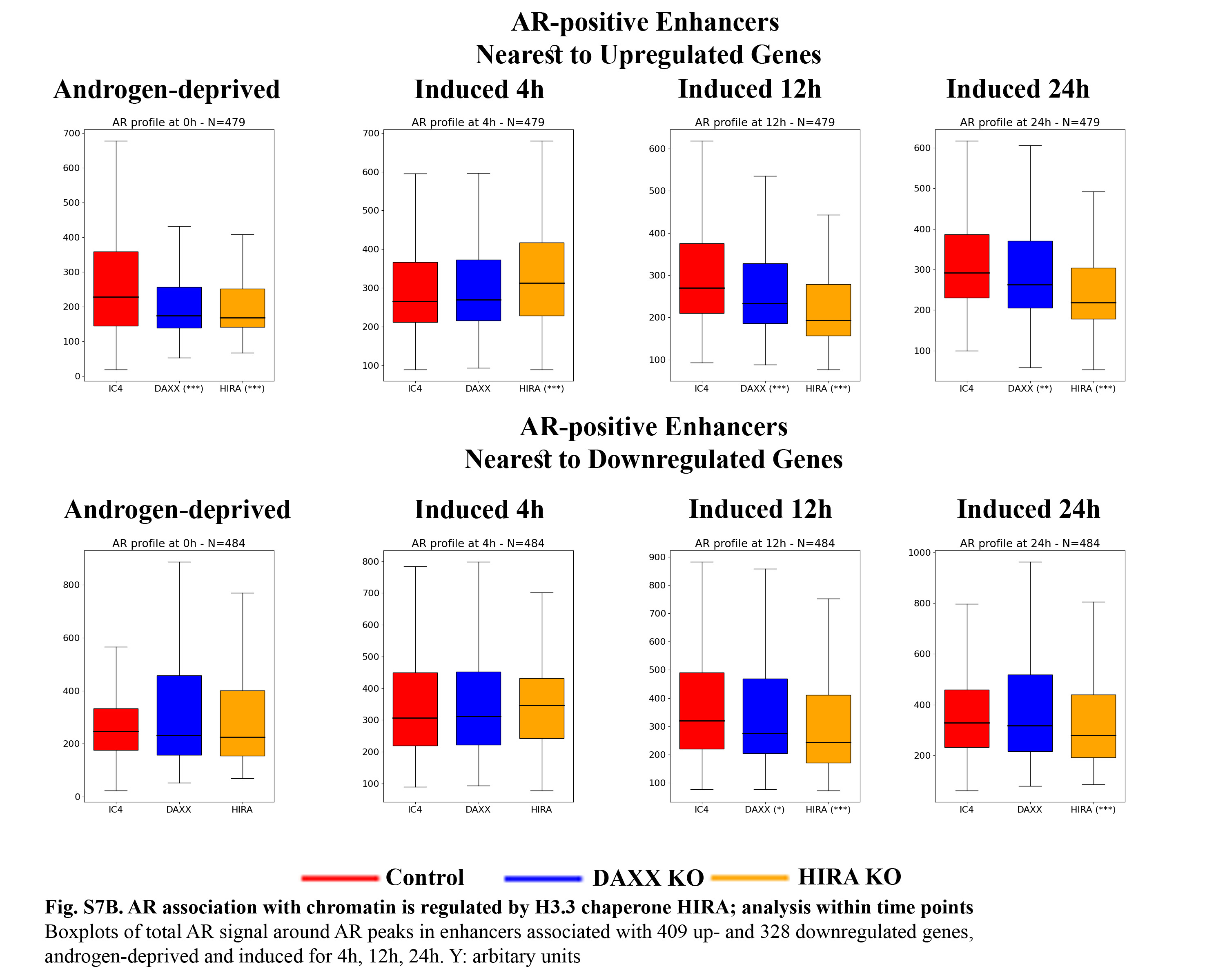

### Fig. S8

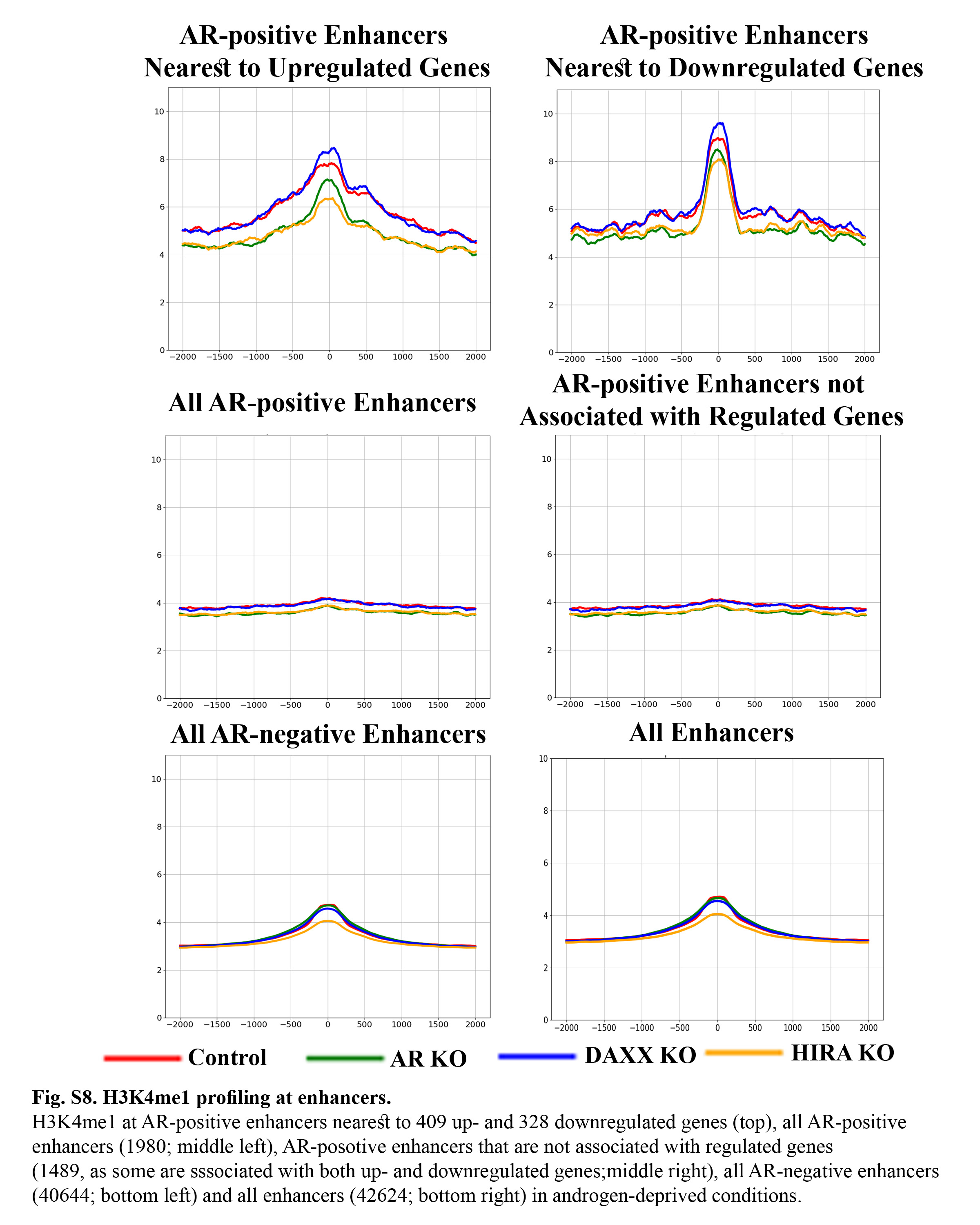

### Fig. S9A

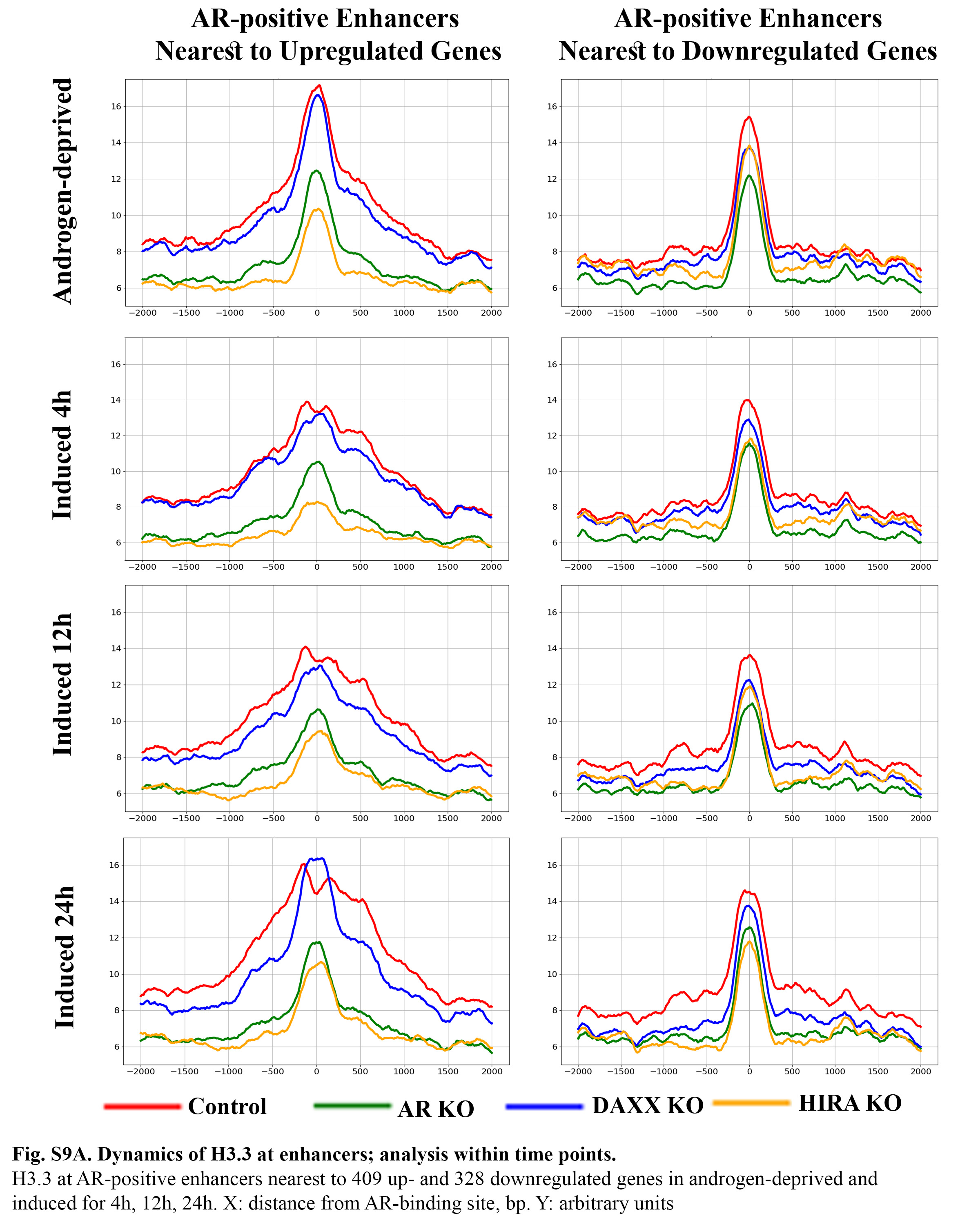

### Fig. S9B

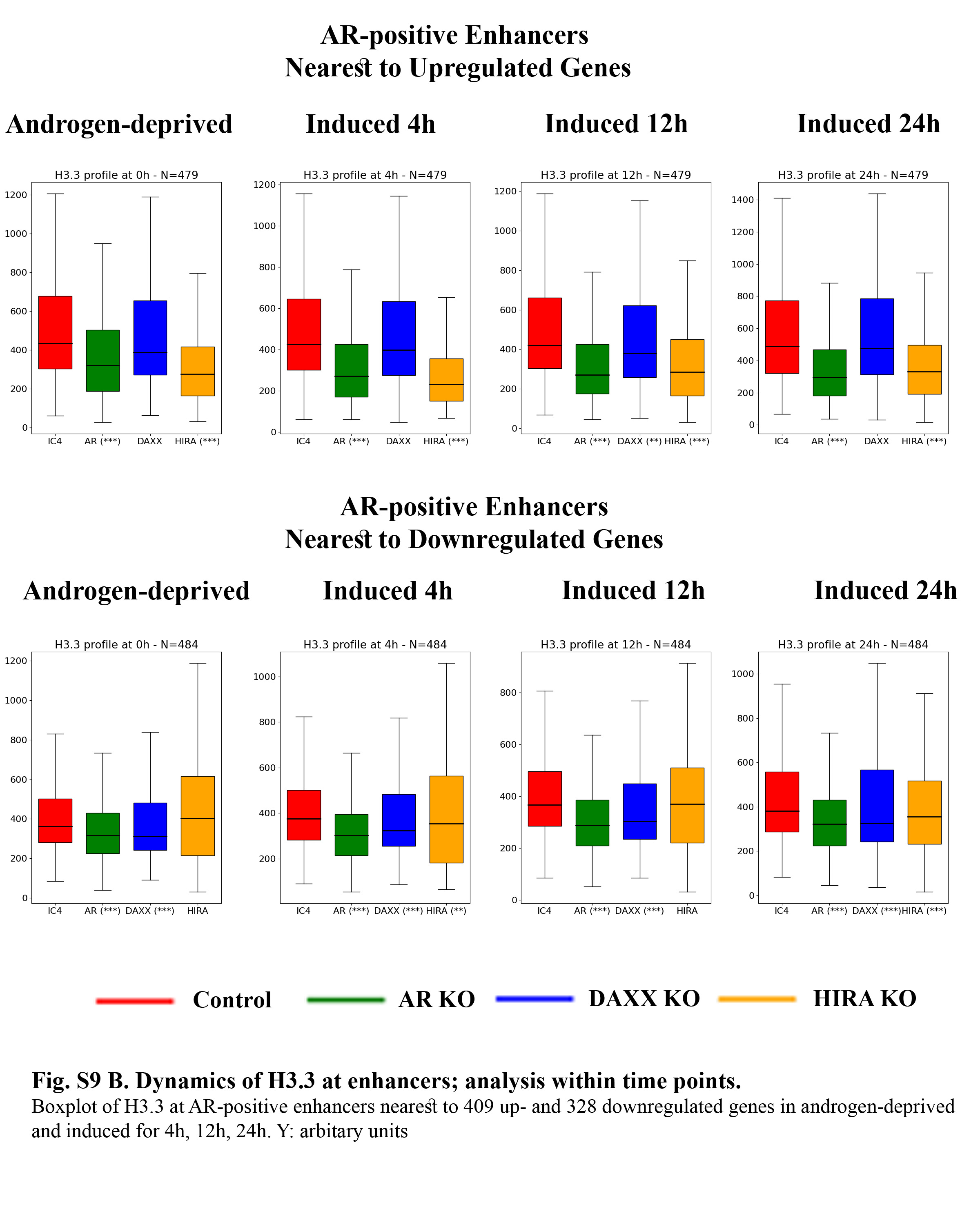

### Fig. S10A

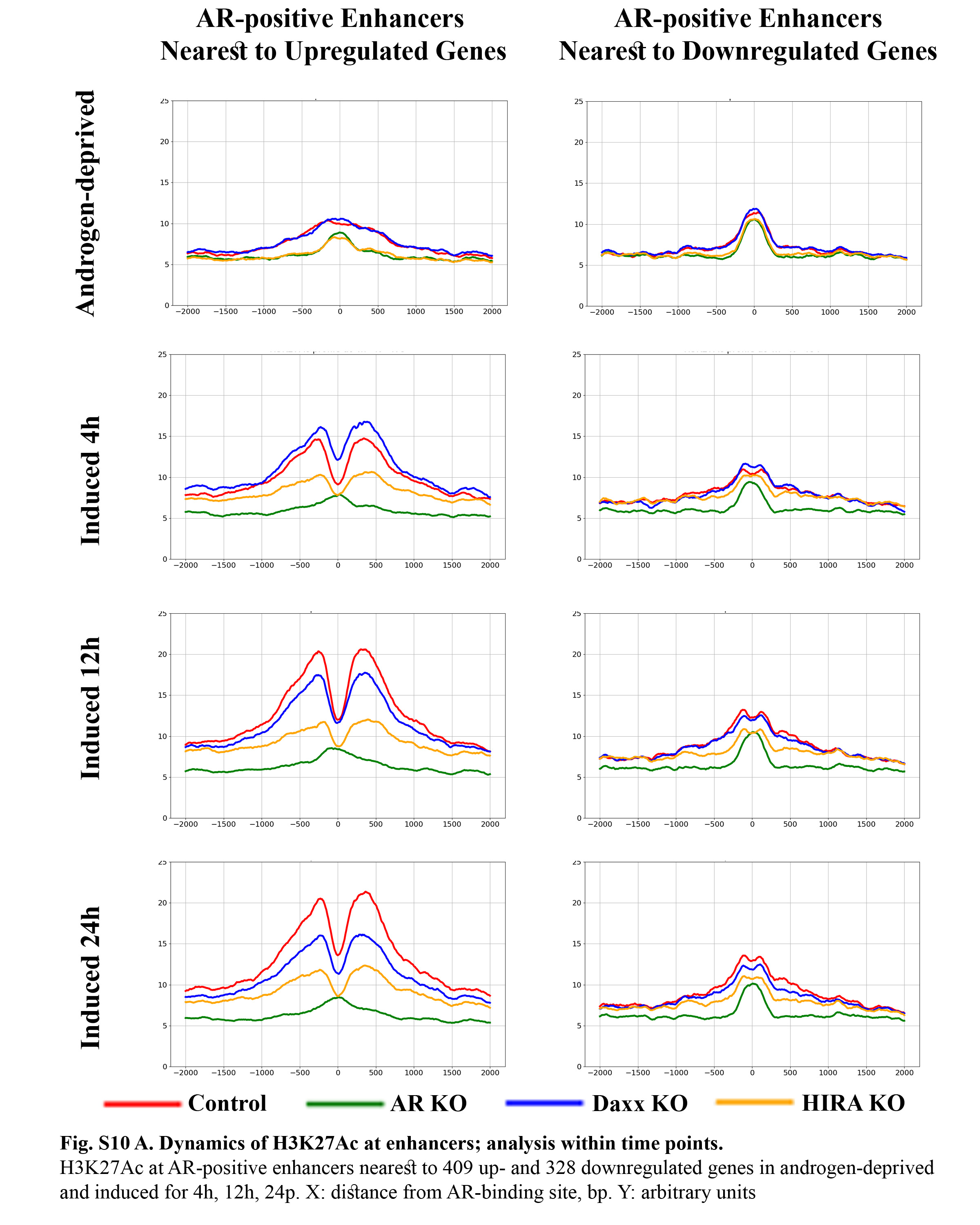

### Fig. S10B

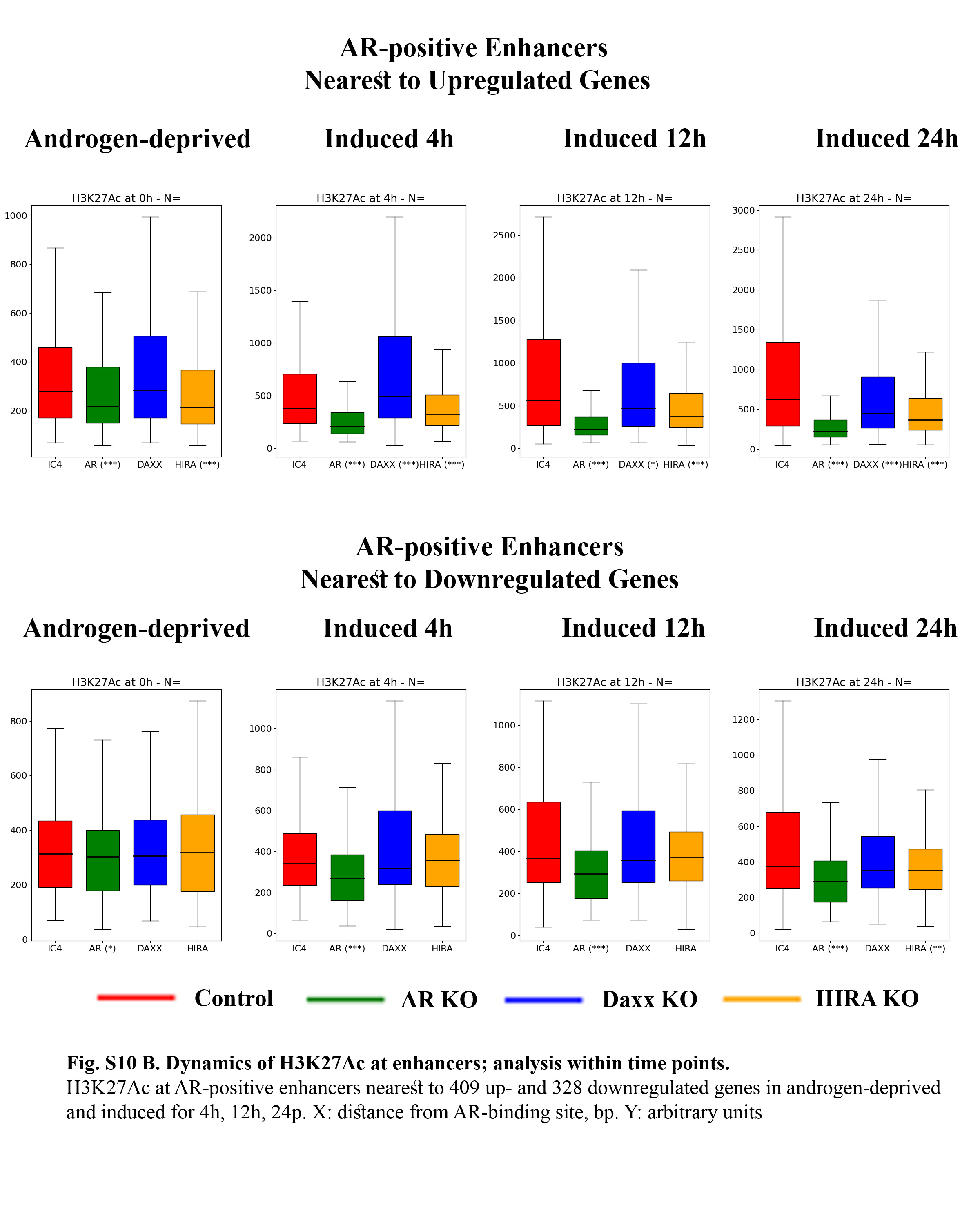

### Fig. S11A

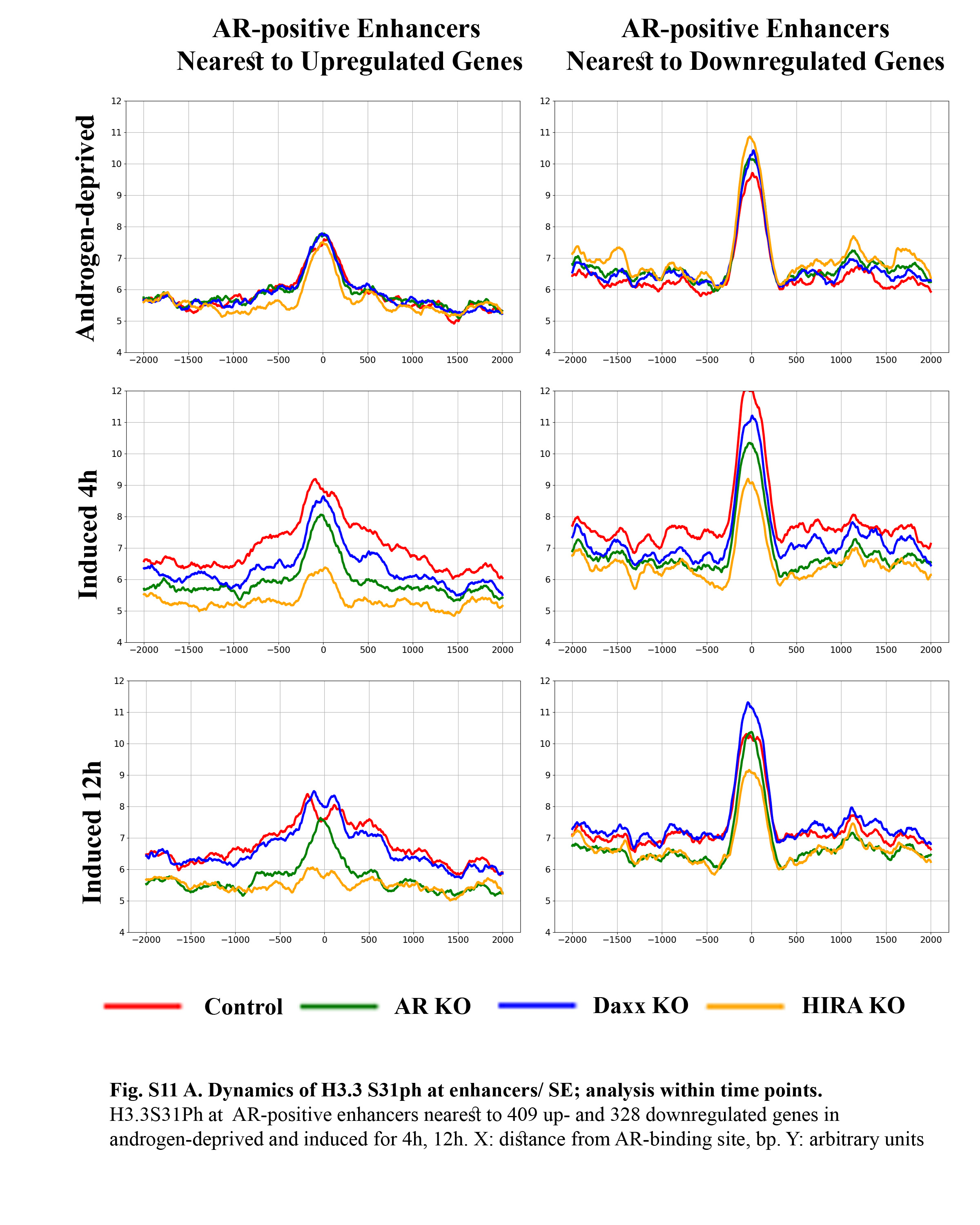

### Fig. S11B

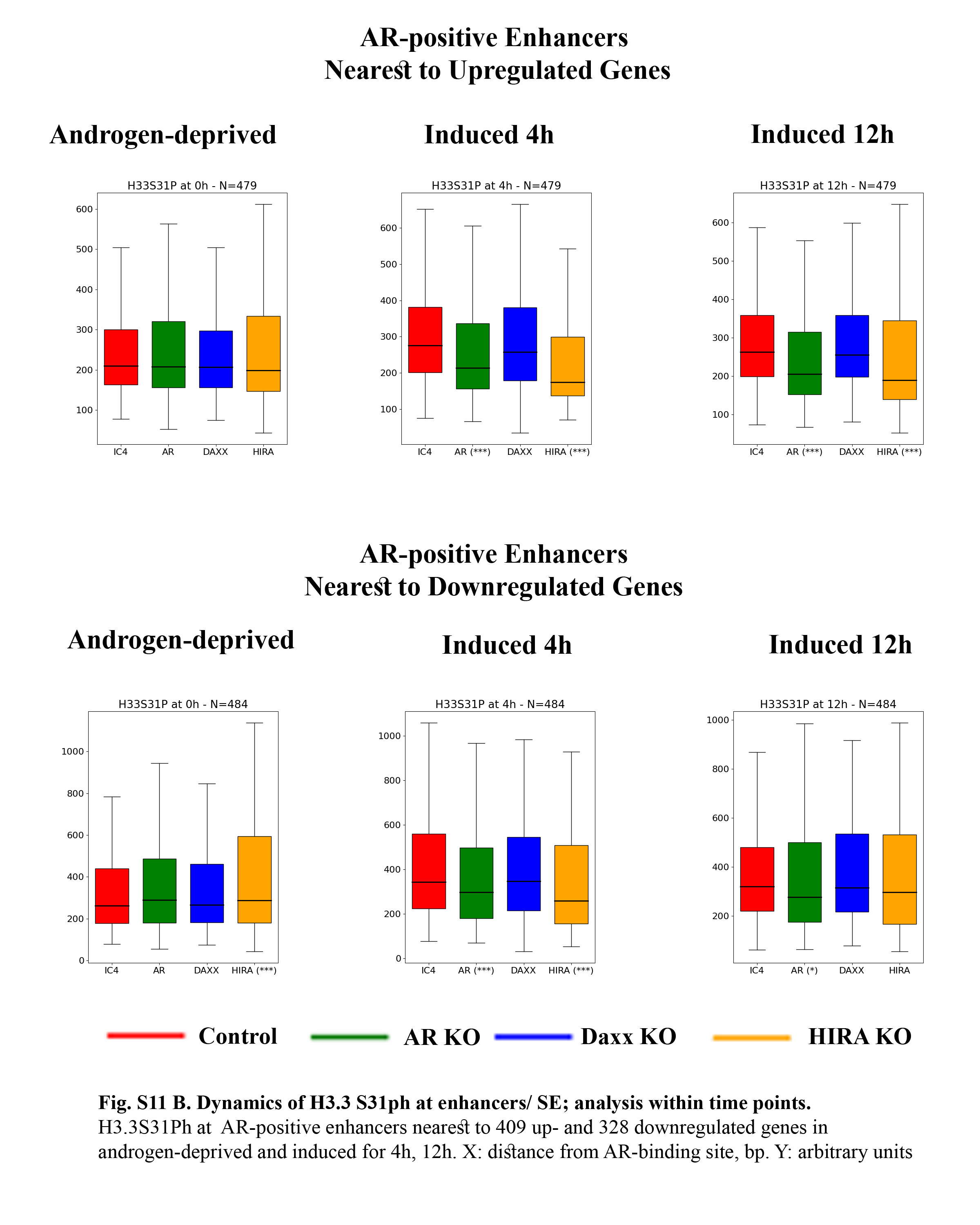

### Fig. S12A

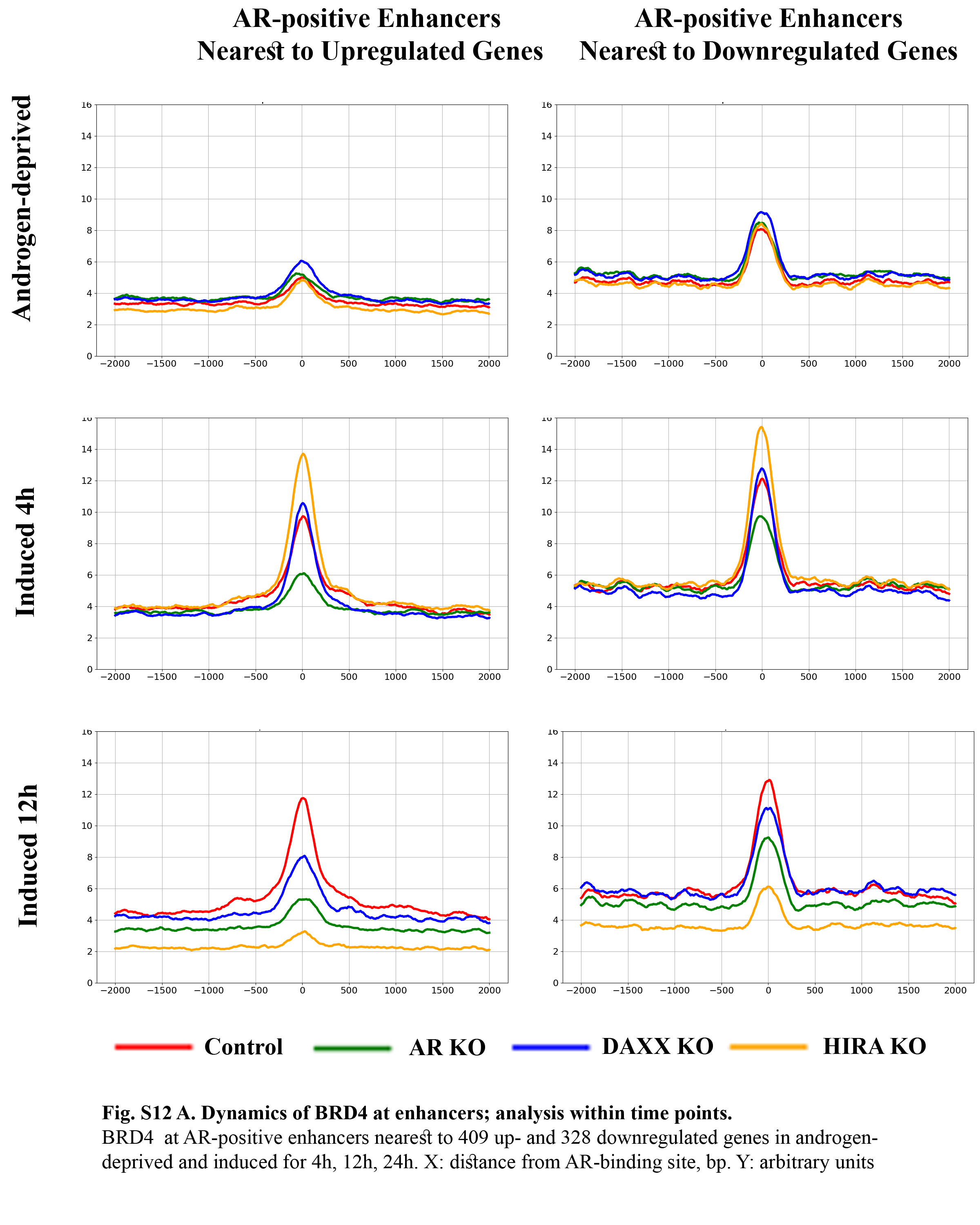

### Fig. S12B

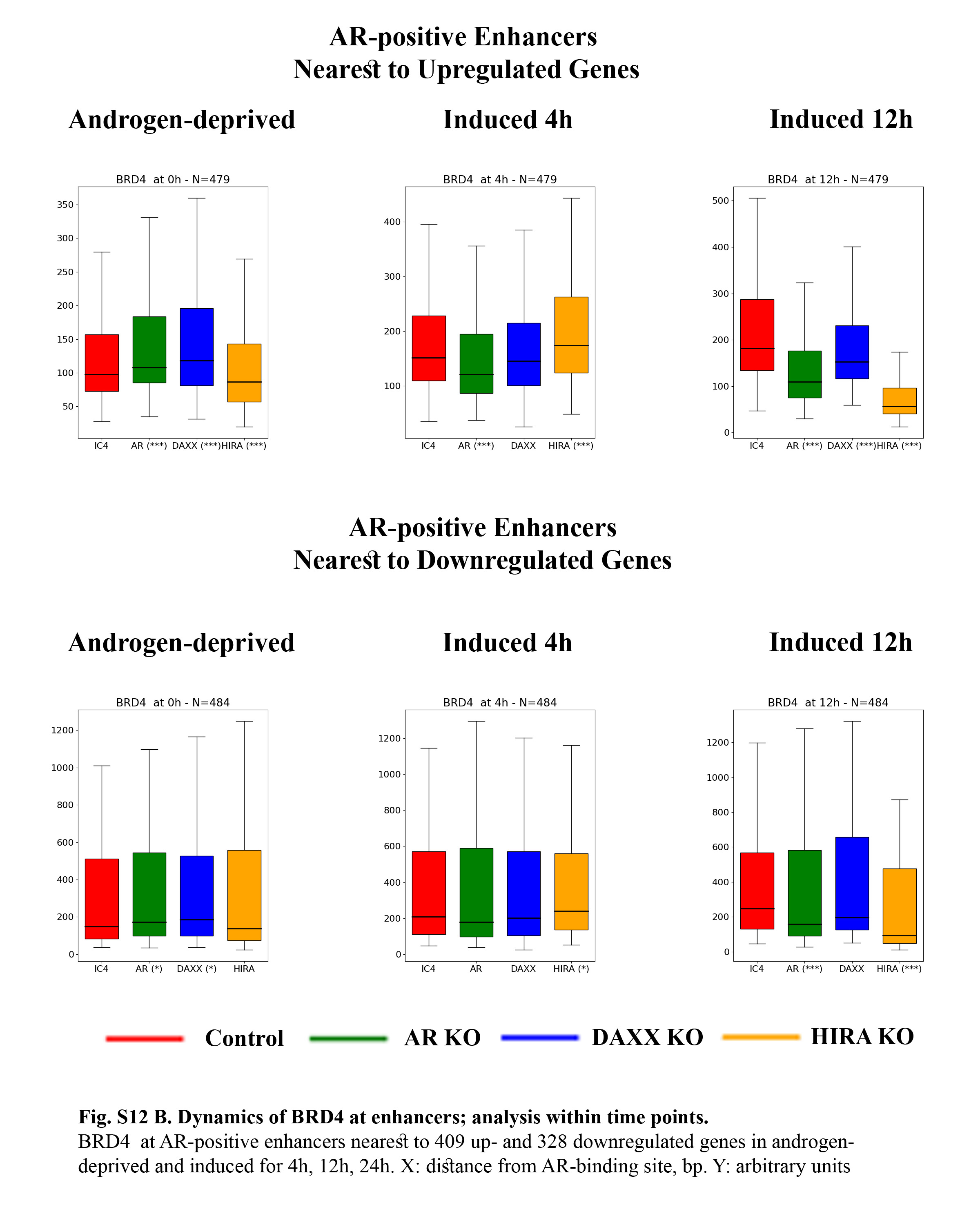

### Fig. S13

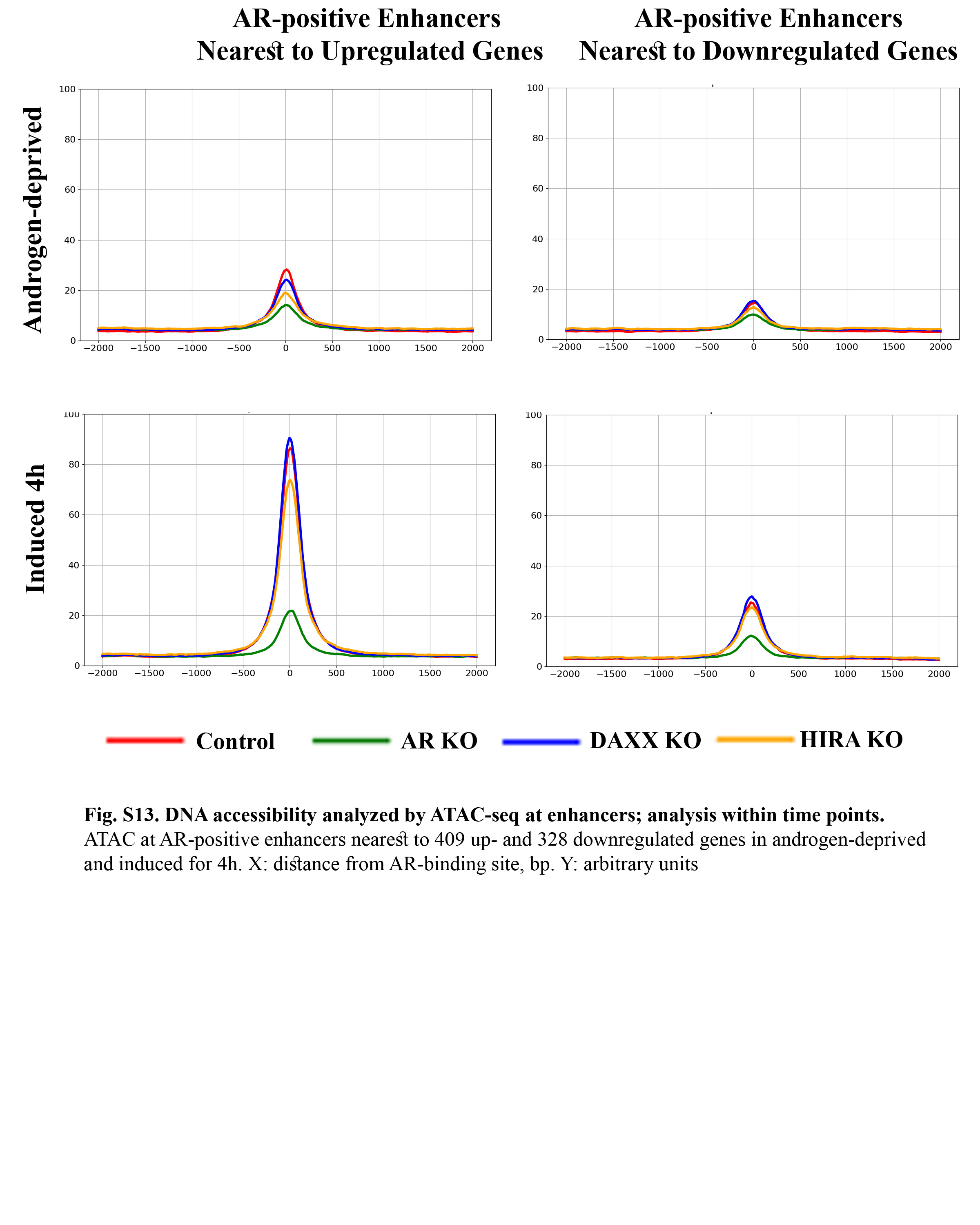

### Fig. S15

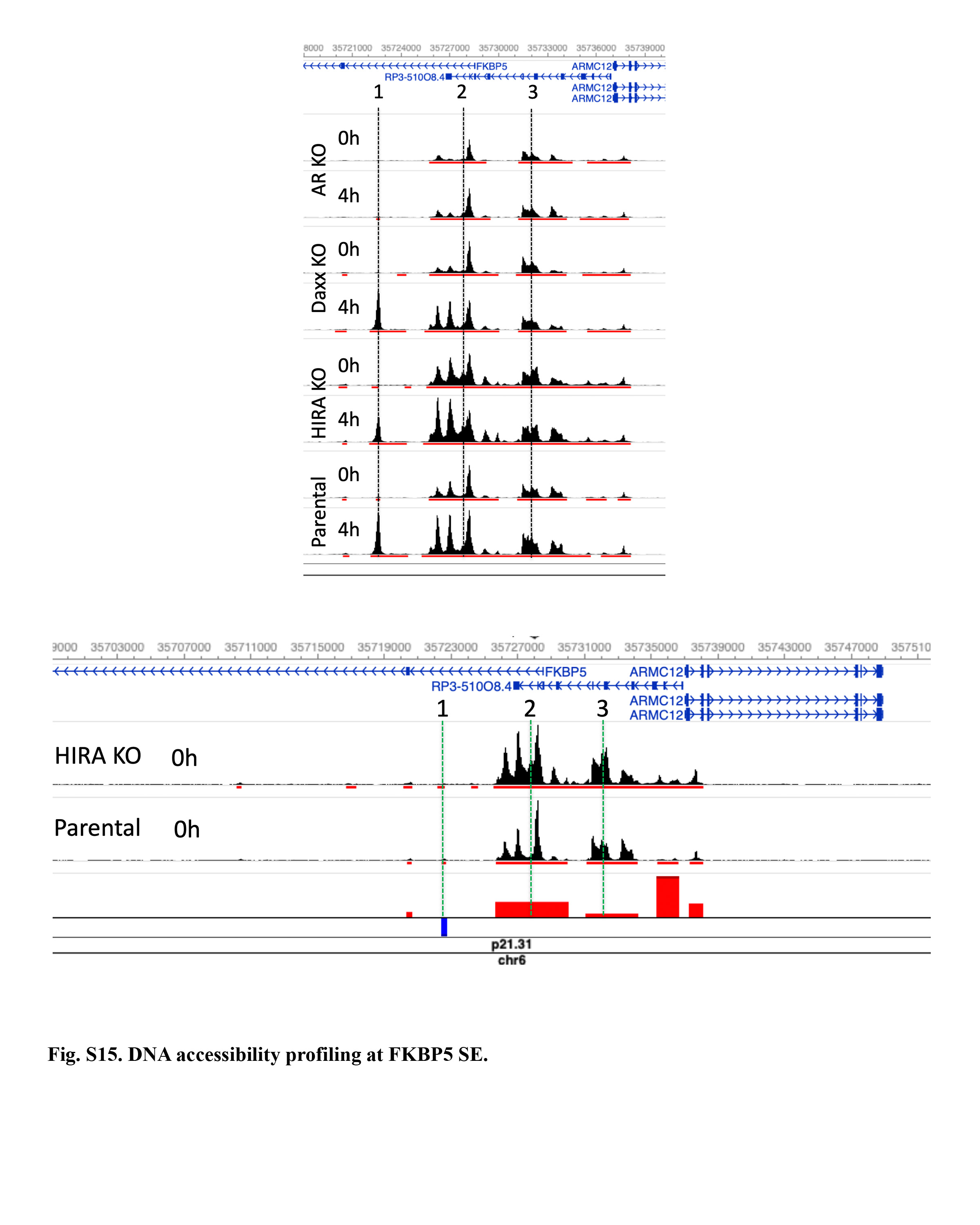

### Fig. S16

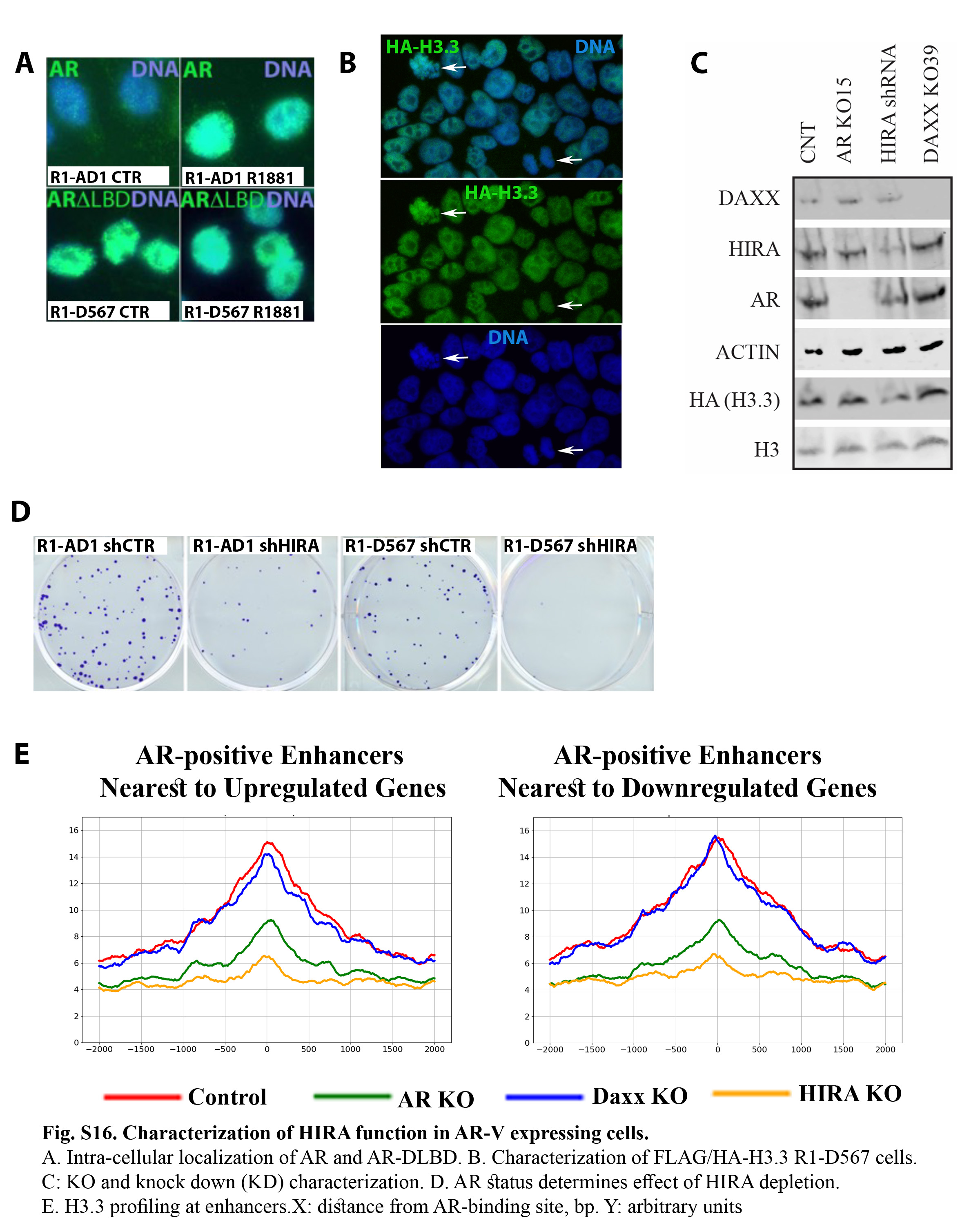

### Fig. S17

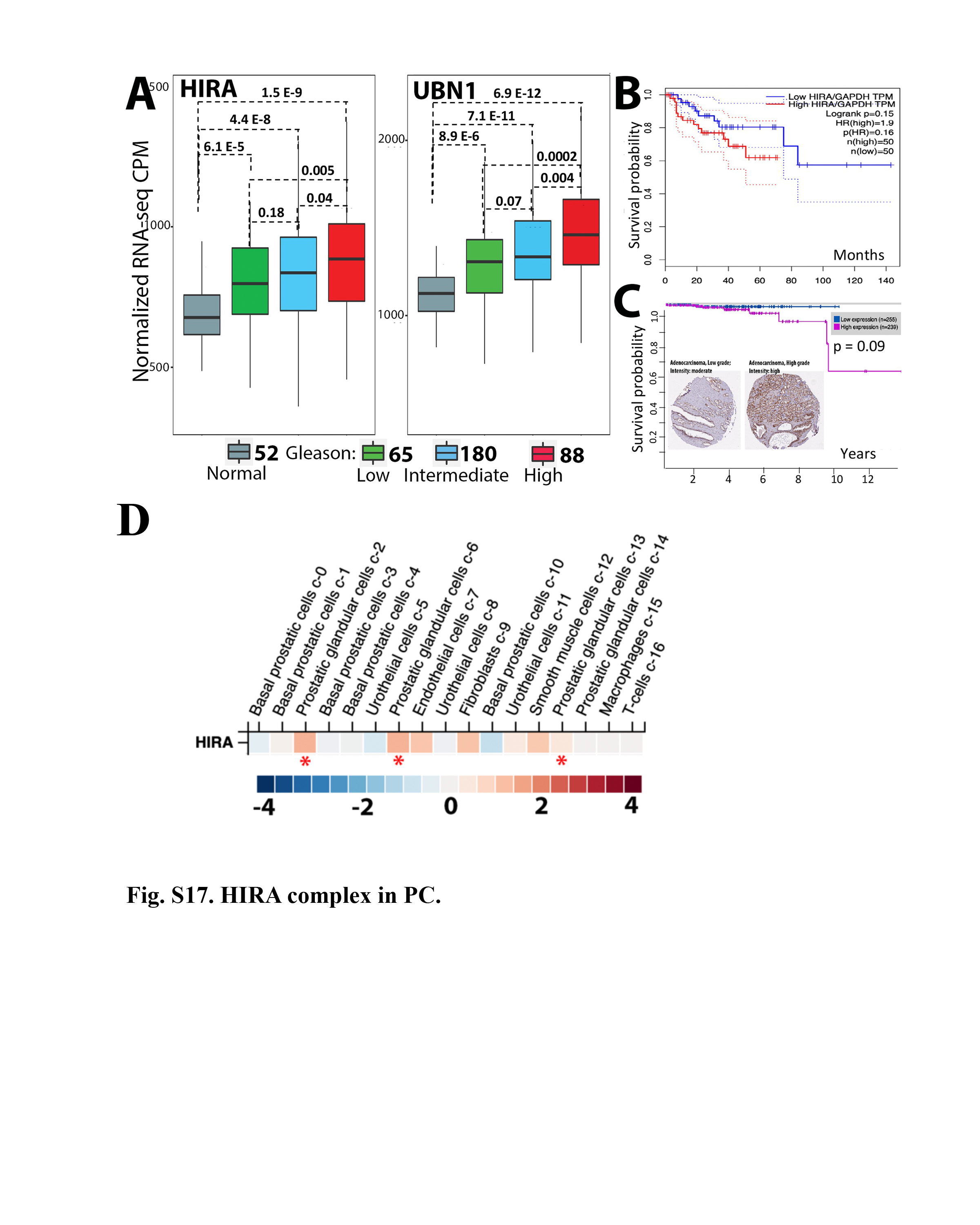

### Fig. S18

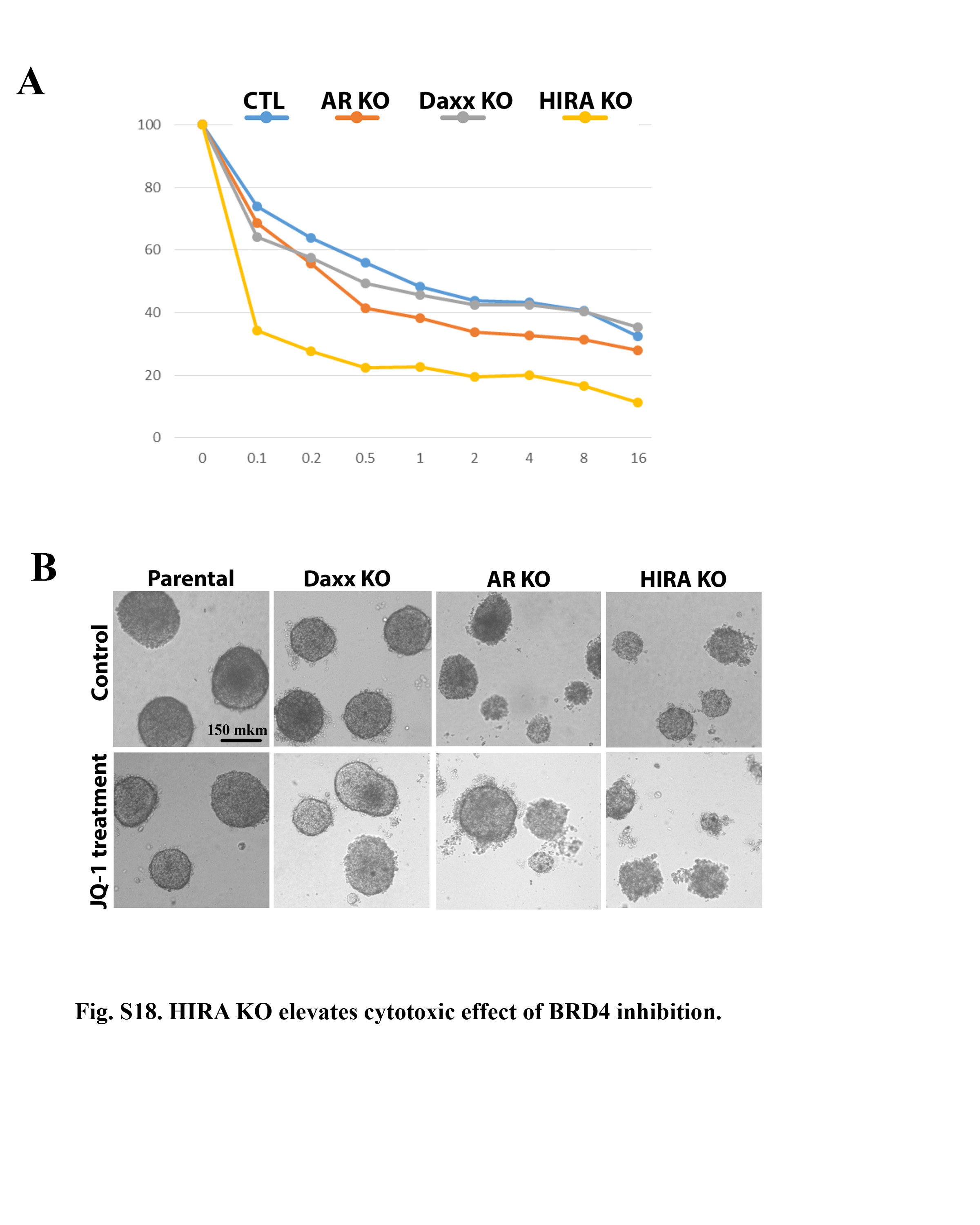
