## Supplementary material for "HIRA-mediated loading of histone variant H3.3 controls androgen-induced transcription by regulation of AR/BRD4 complex assembly at enhancers": Fig. S2

**Fig. S2. Expression analysis (RNA-seq) of R1-AD1 cells, parental (WT), AR KO, Daxx KO and HIRA KO.**

**A**

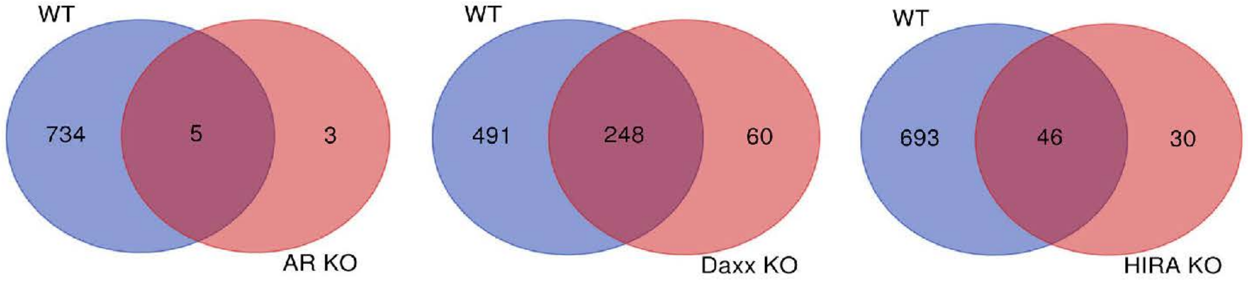


**B**

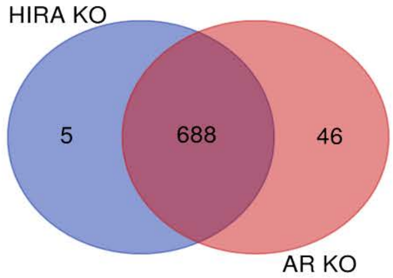


Cancer pathway

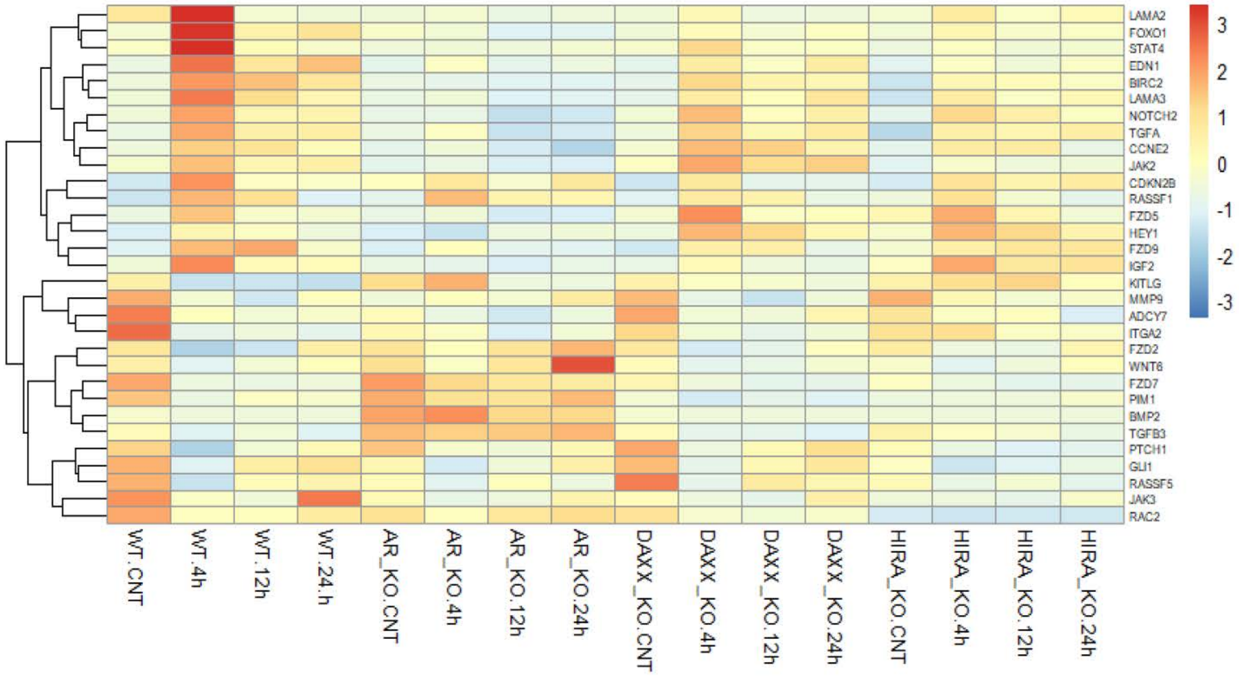

Hippo pathway

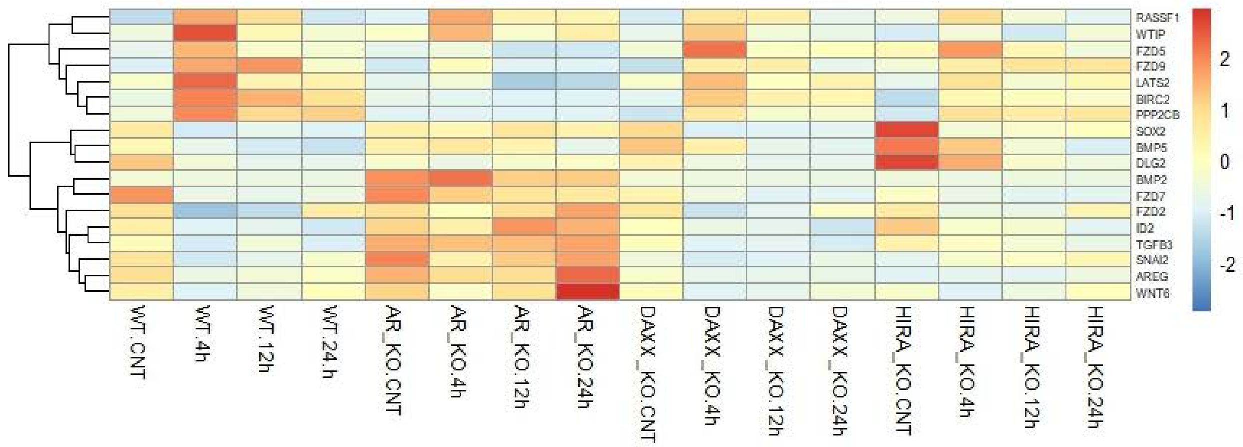


FoxO pathway

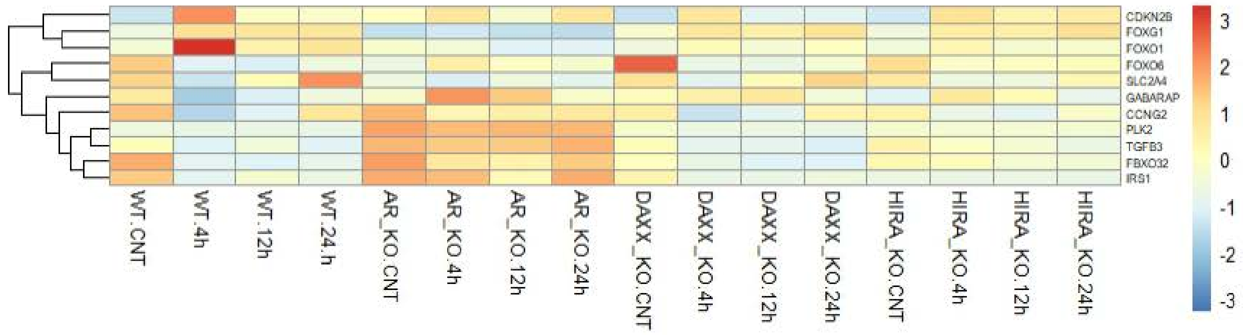


TGF-beta pathway

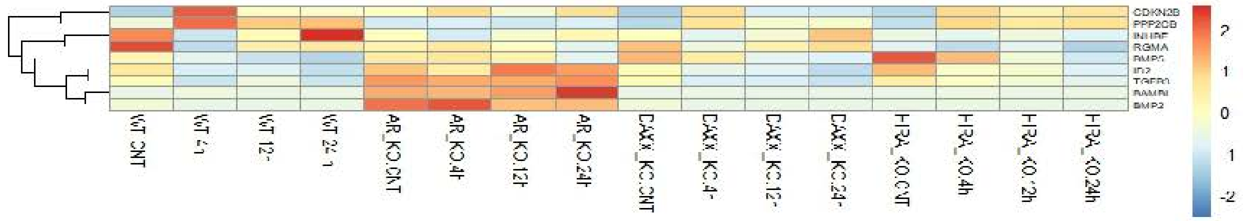


Wnt pathway

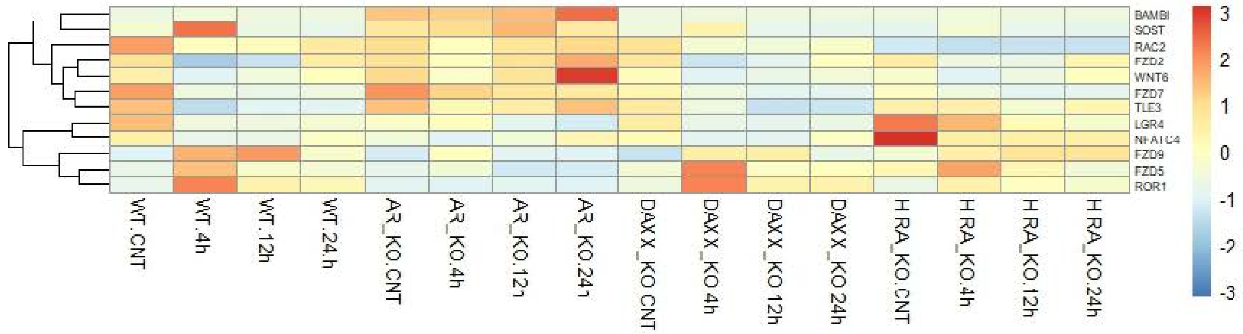

PI3-Akt pathway

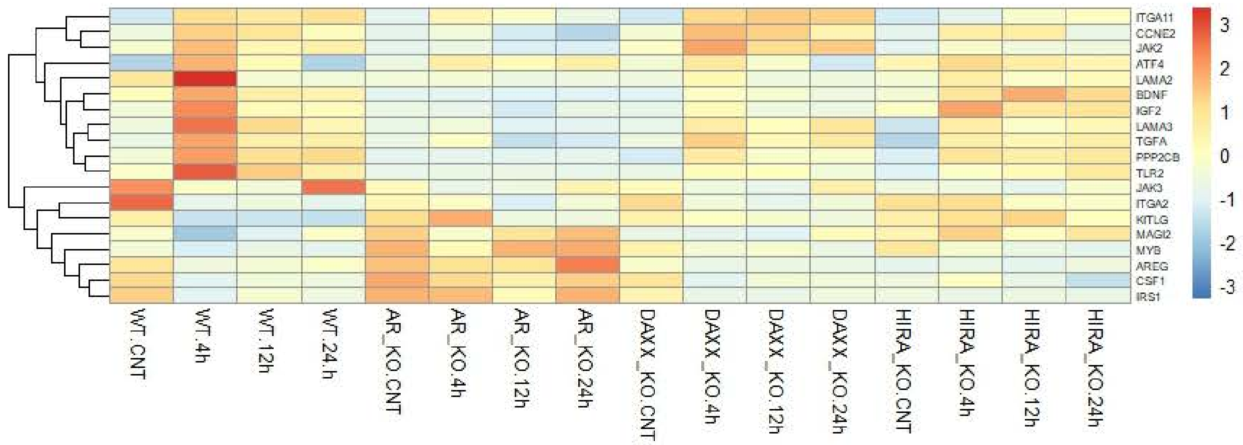
