## Supplementary material for "HIRA-mediated loading of histone variant H3.3 controls androgen-induced transcription by regulation of AR/BRD4 complex assembly at enhancers": Fig. S14

**Fig. S14A**

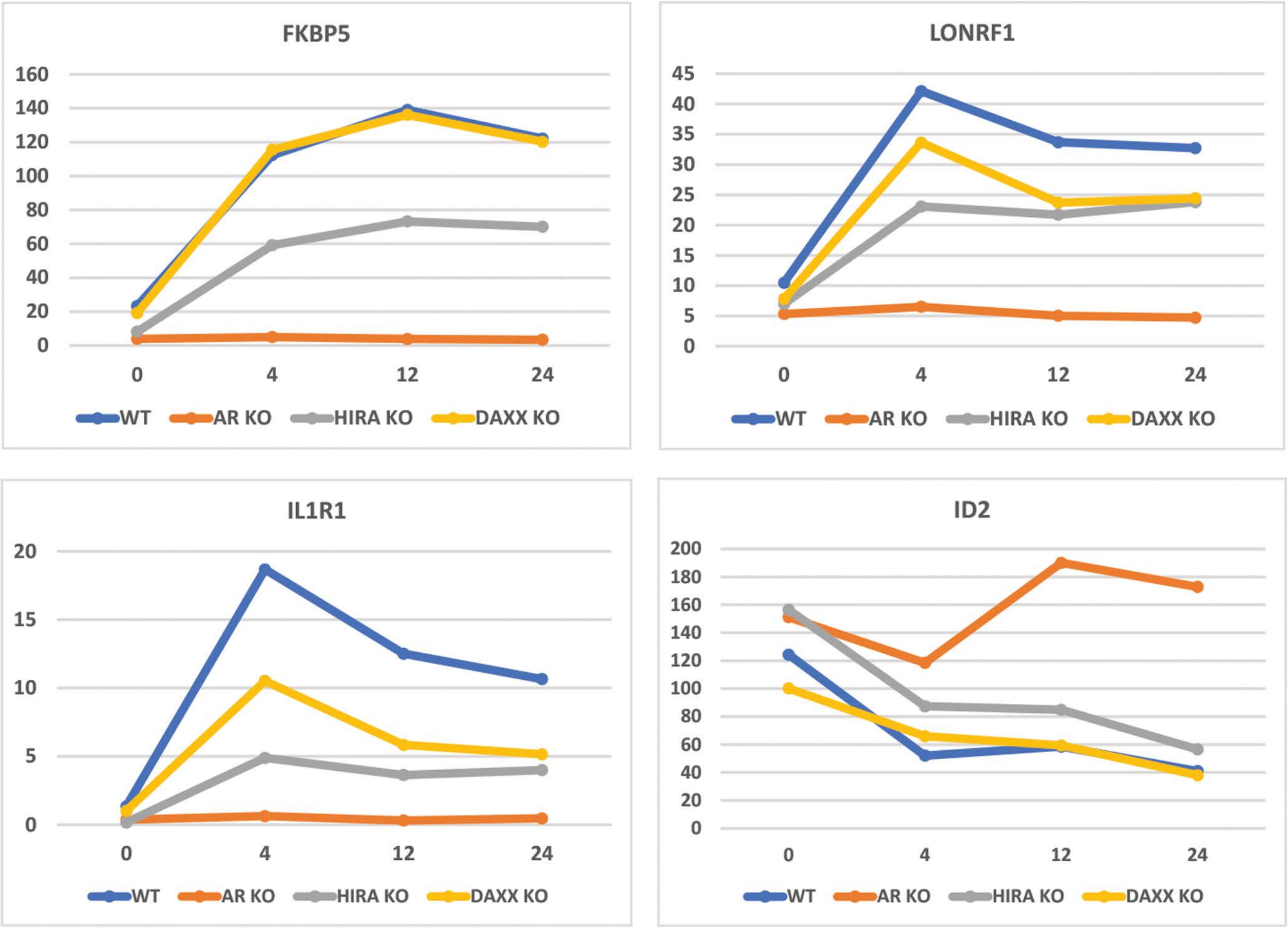


**Fig. S14. Examples of epigenetic profiles of enhancers/ SE associated with genes co-regulated by AR and HIRA.**
A: Androgen-induced expression of LONRF1, IL1R1, FKBP5 genes is reduced by AR and HIRA KO, and androgen-induced repression of ID2 gene is elevated by AR and HIRAKO (results of RNA-seq analysis).

**Fig. S14B**

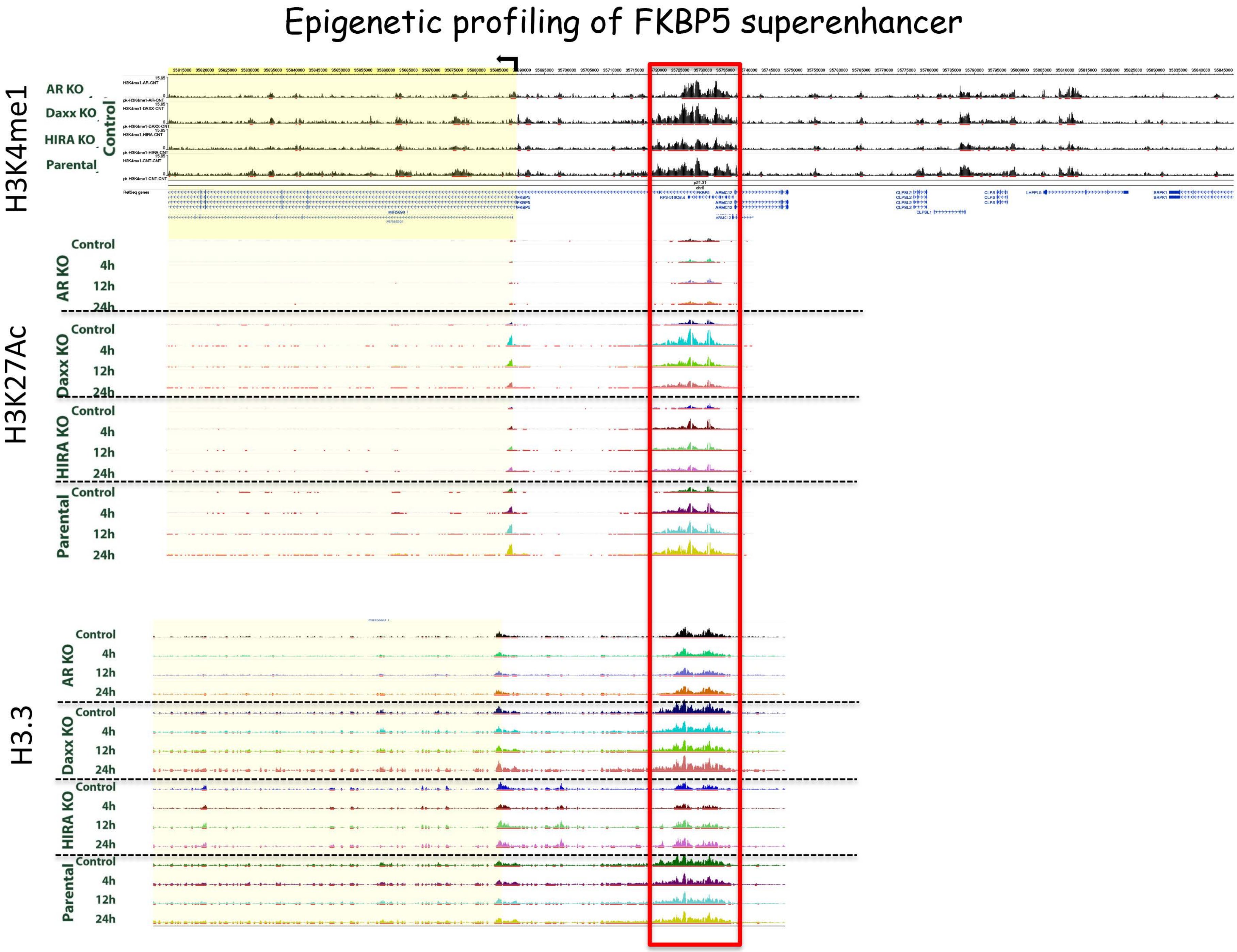


**Fig. S14. Examples of epigenetic profiles of enhancers/ SE associated with genes co-regulated by AR and HIRA.**
B: H3K4me1 (enhancer), H3K27Ac (active enhancer), H3.3, H3.3S31Ph, AR, BRD4, and CTCF profiling were analyzed by ChIP-seq. DNA accessibility analyzed by ATAC-seq. Arrows: transcription start; red lines outline SE regions.

**Fig. S14B**

Epigenetic profiling of FKBP5 superenhancer

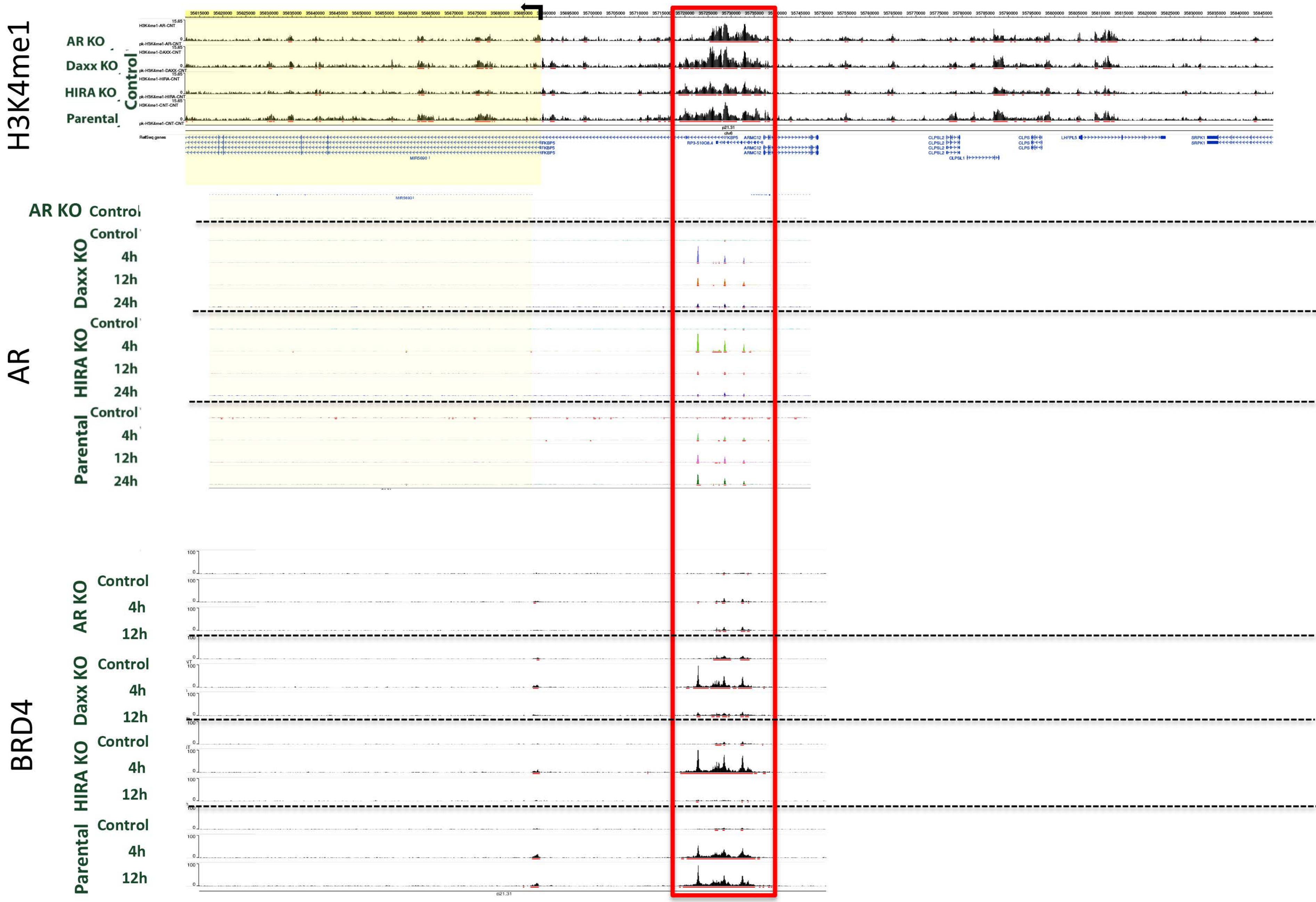

**Fig. S14B**

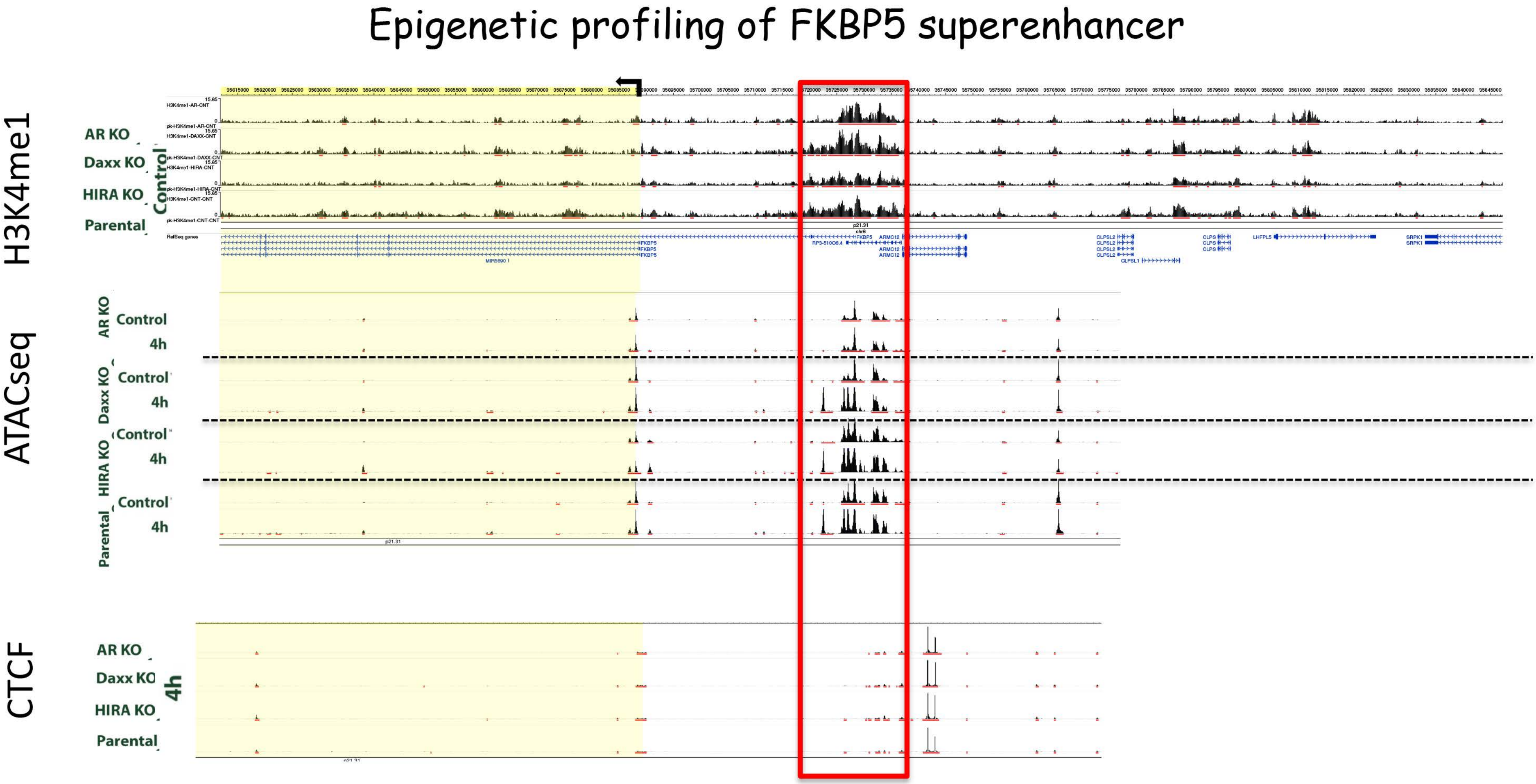
Epigenetic profiling of FKBP5 superenhancer
H3K4me1
AR KO
Daxx KO
HIRA KO
Parental
Control
ATACseq
AR KO
Control
4h
Daxx KO
Control
4h
HIRA KO
Control
4h
Parental
Control
4h
CTCF
AR KO
Daxx KO
HIRA KO
Parental
4h

### Fig. S14B

Epigenetic profiling of LONRF1 superenhancer

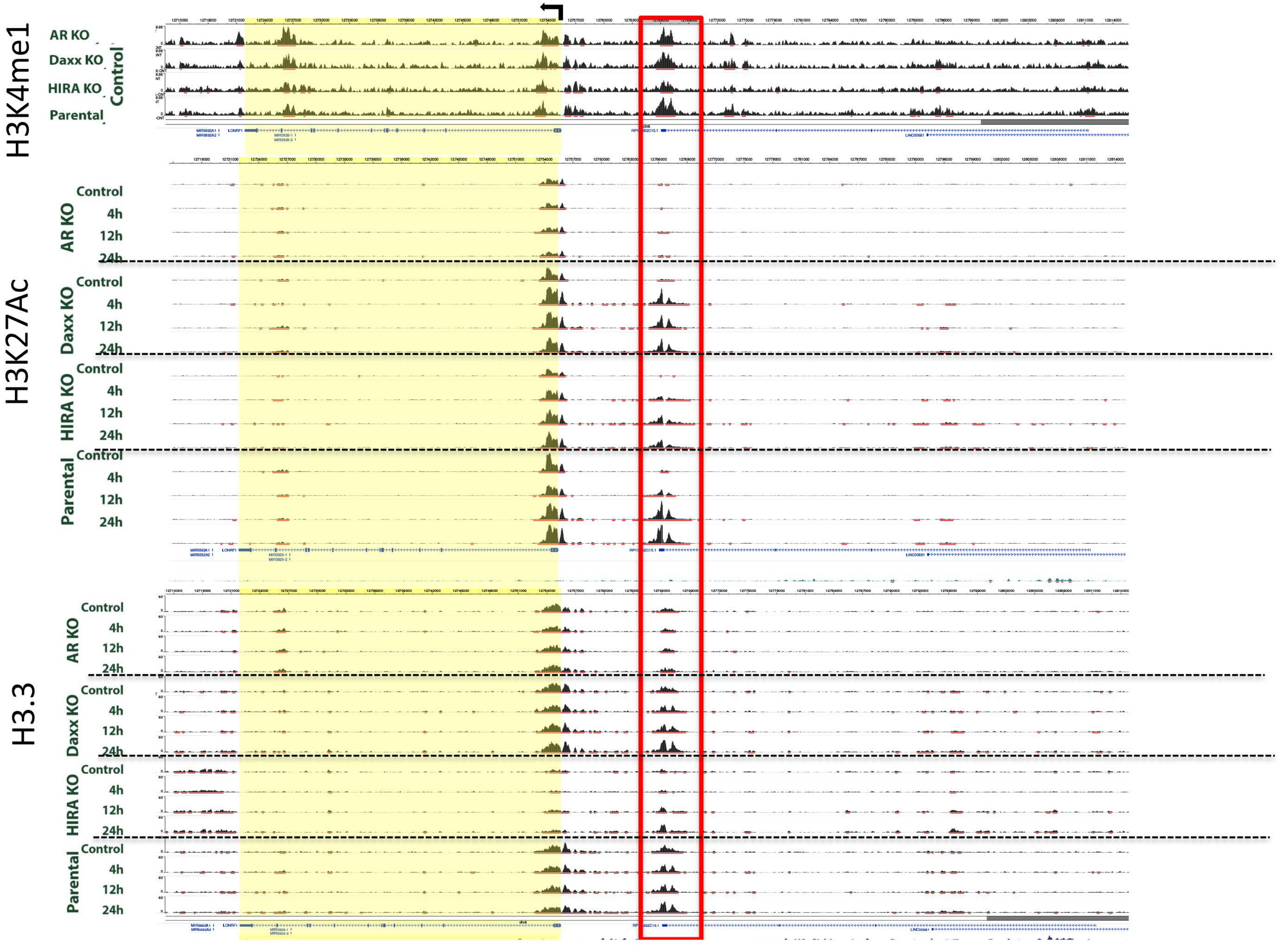

**Fig. S14B**

Epigenetic profiling of LONRF1 superenhancer

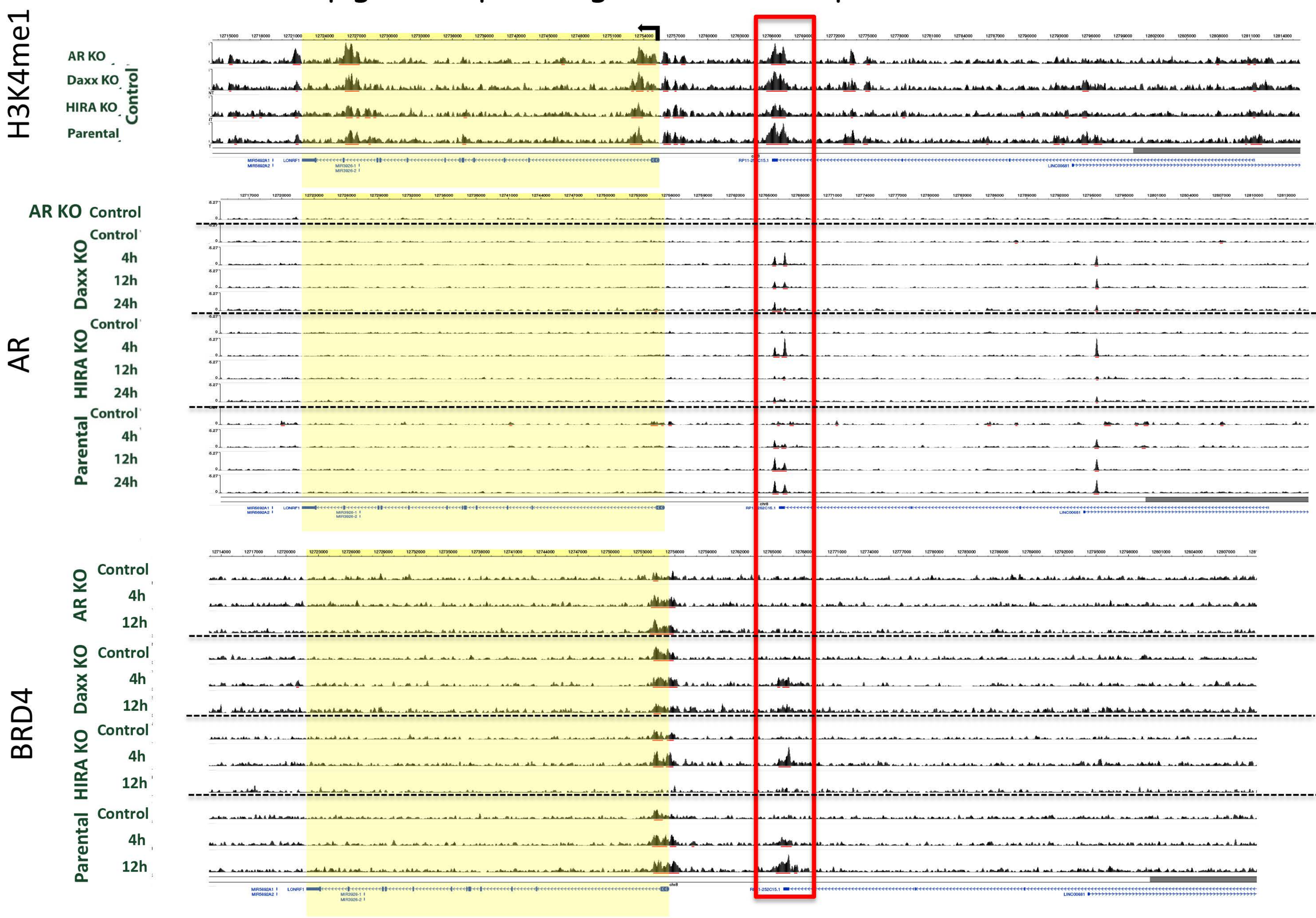

### Fig. S14B

Epigenetic profiling of LONRF1 superenhancer

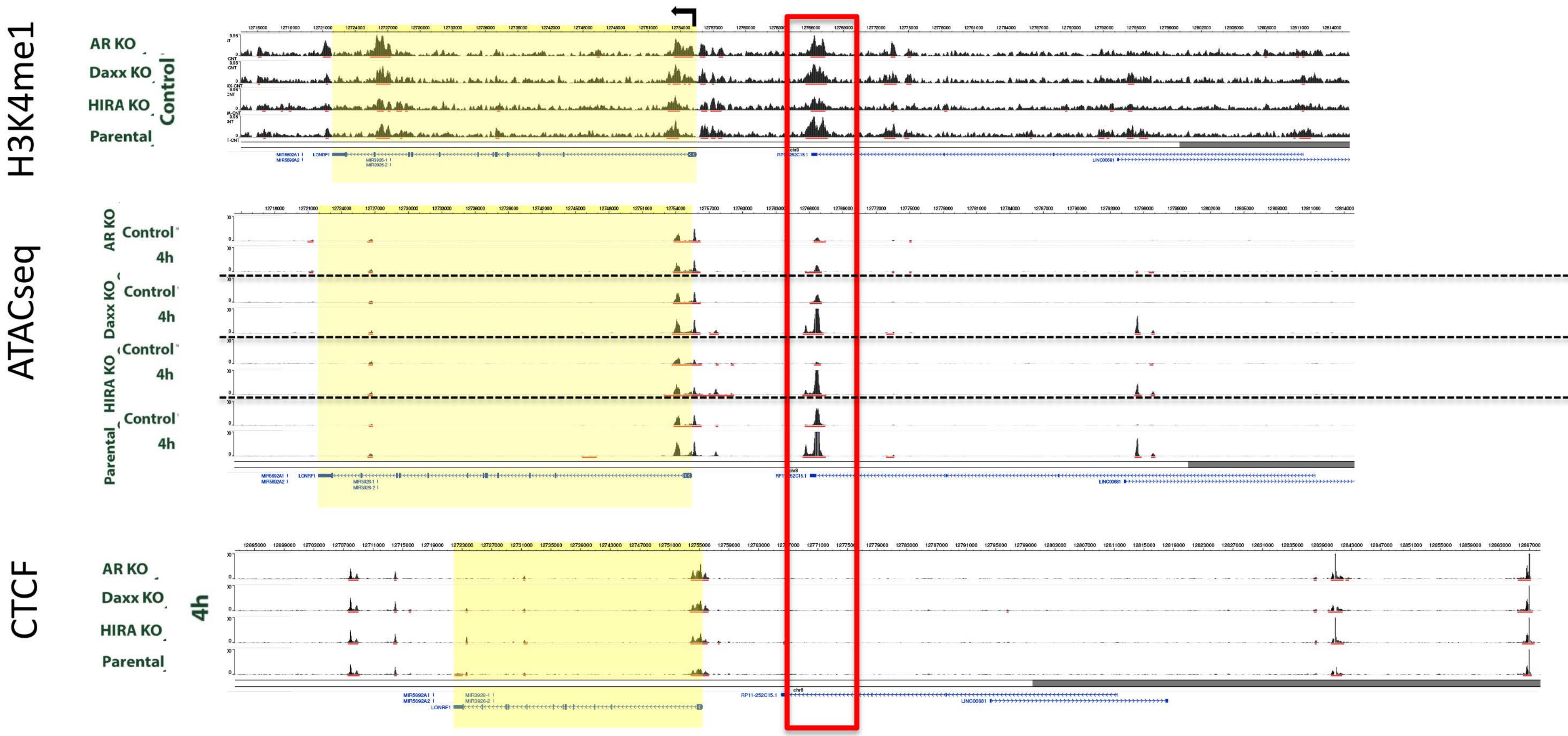

### Fig. S14B

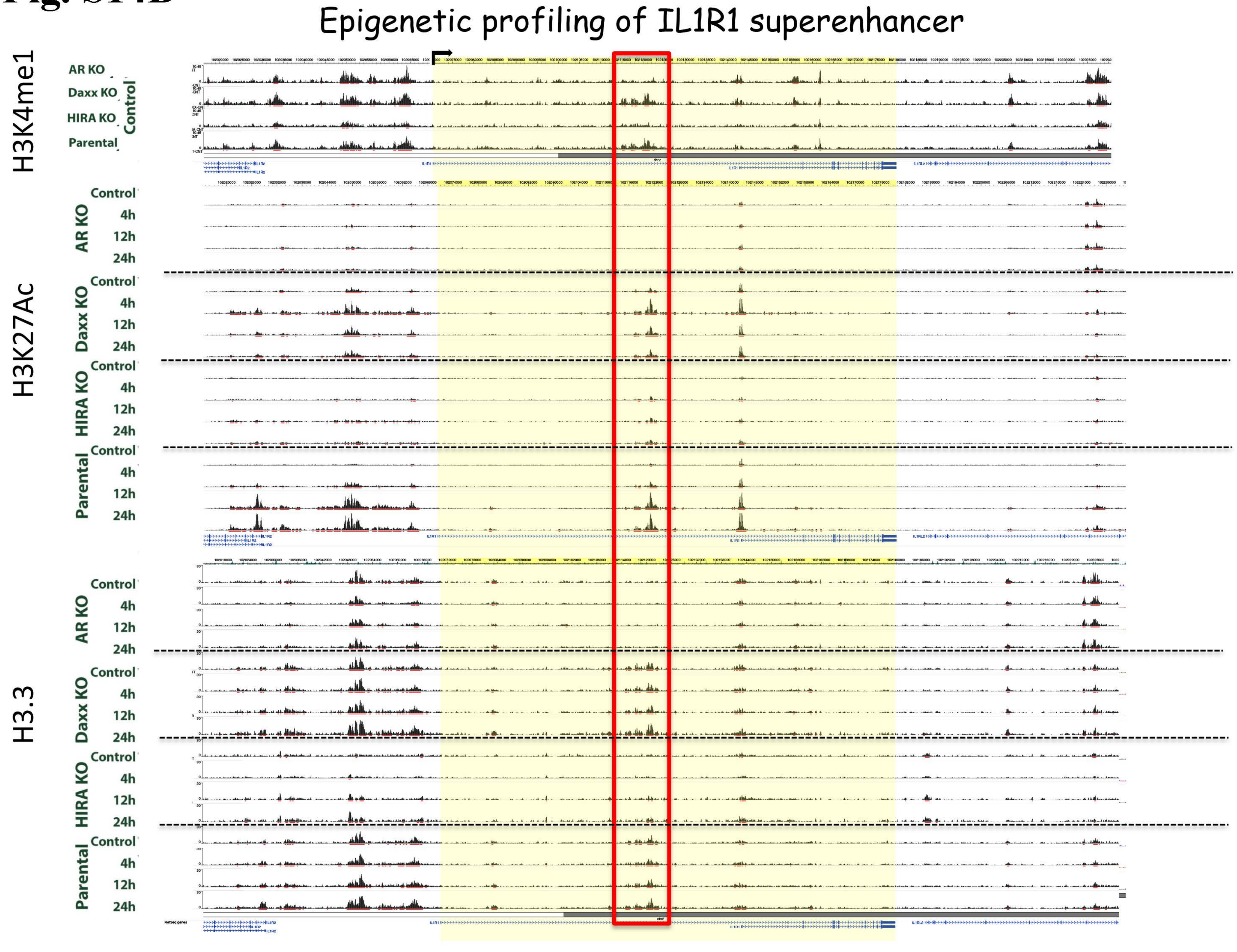

Epigenetic profiling of IL1R1 superenhancer
H3K4me1
AR KO
Daxx KO
HIRA KO
Parental
Control
H3K27Ac
Control
4h
12h
24h
H3.3
IL1R2
IL1R1
IL1RL2

### Fig. S14B

Epigenetic profiling of IL1R1 superenhancer

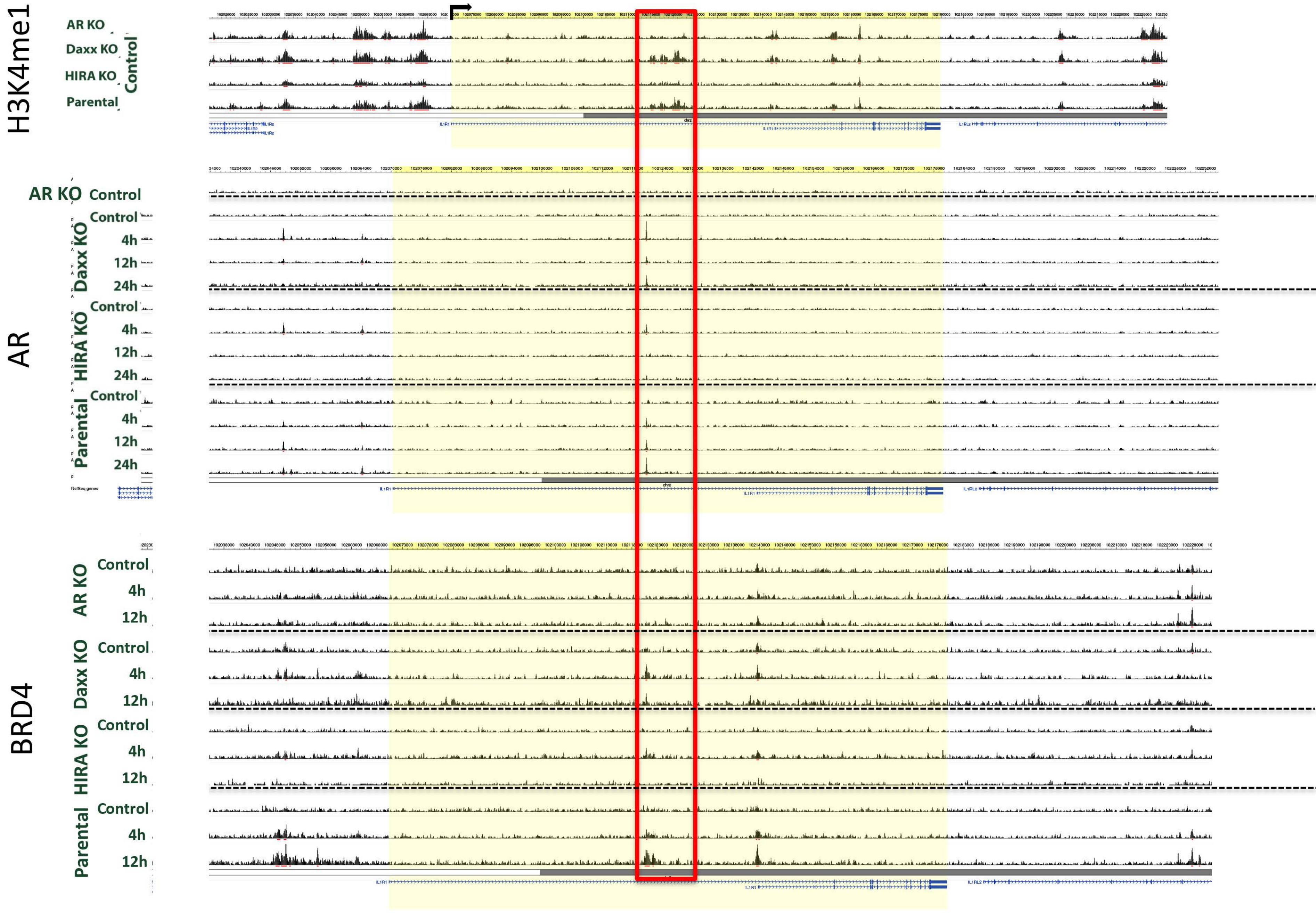

### Fig. S14B

Epigenetic profiling of IL1R1 superenhancer

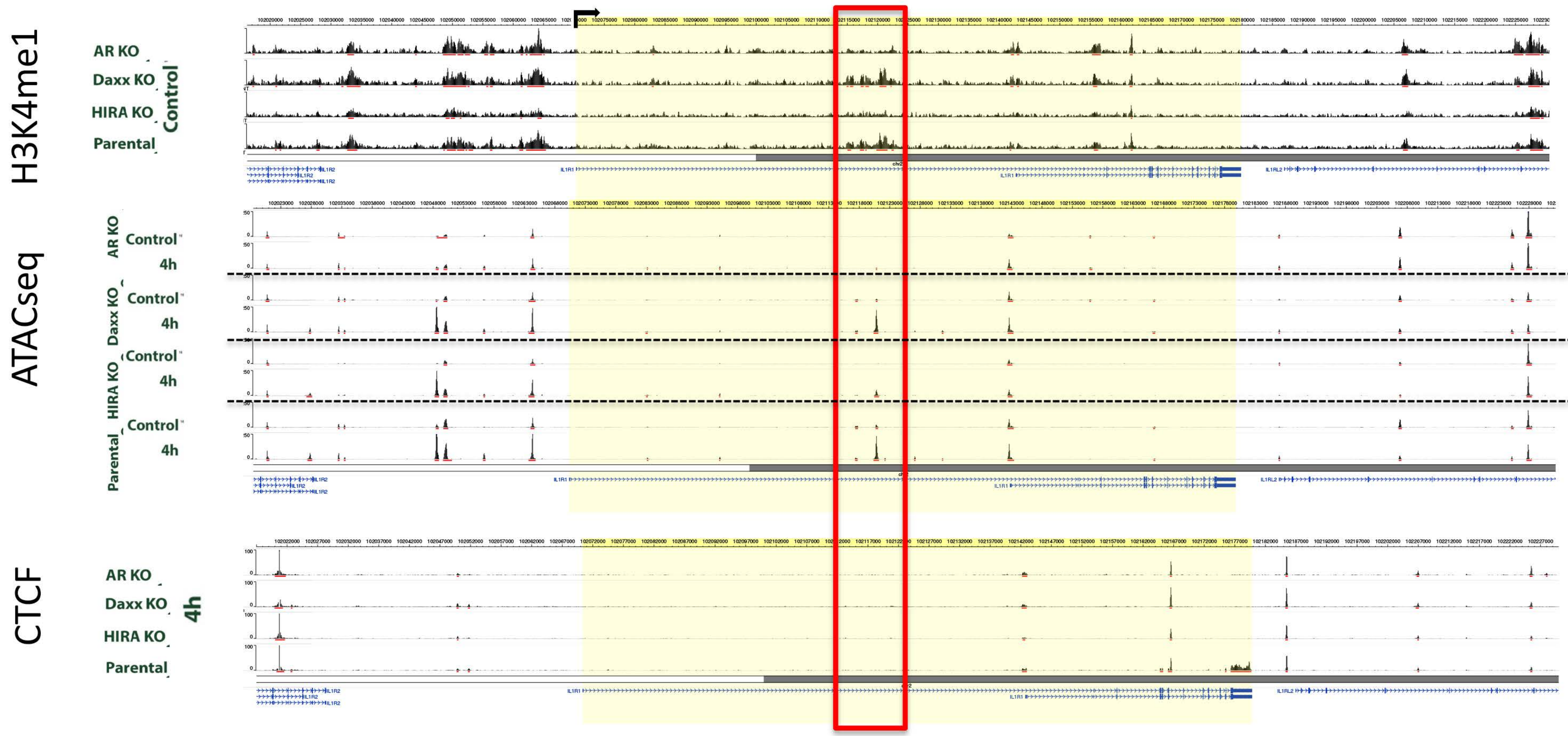

### Fig. S14B

Epigenetic profiling of ID2 superenhancer

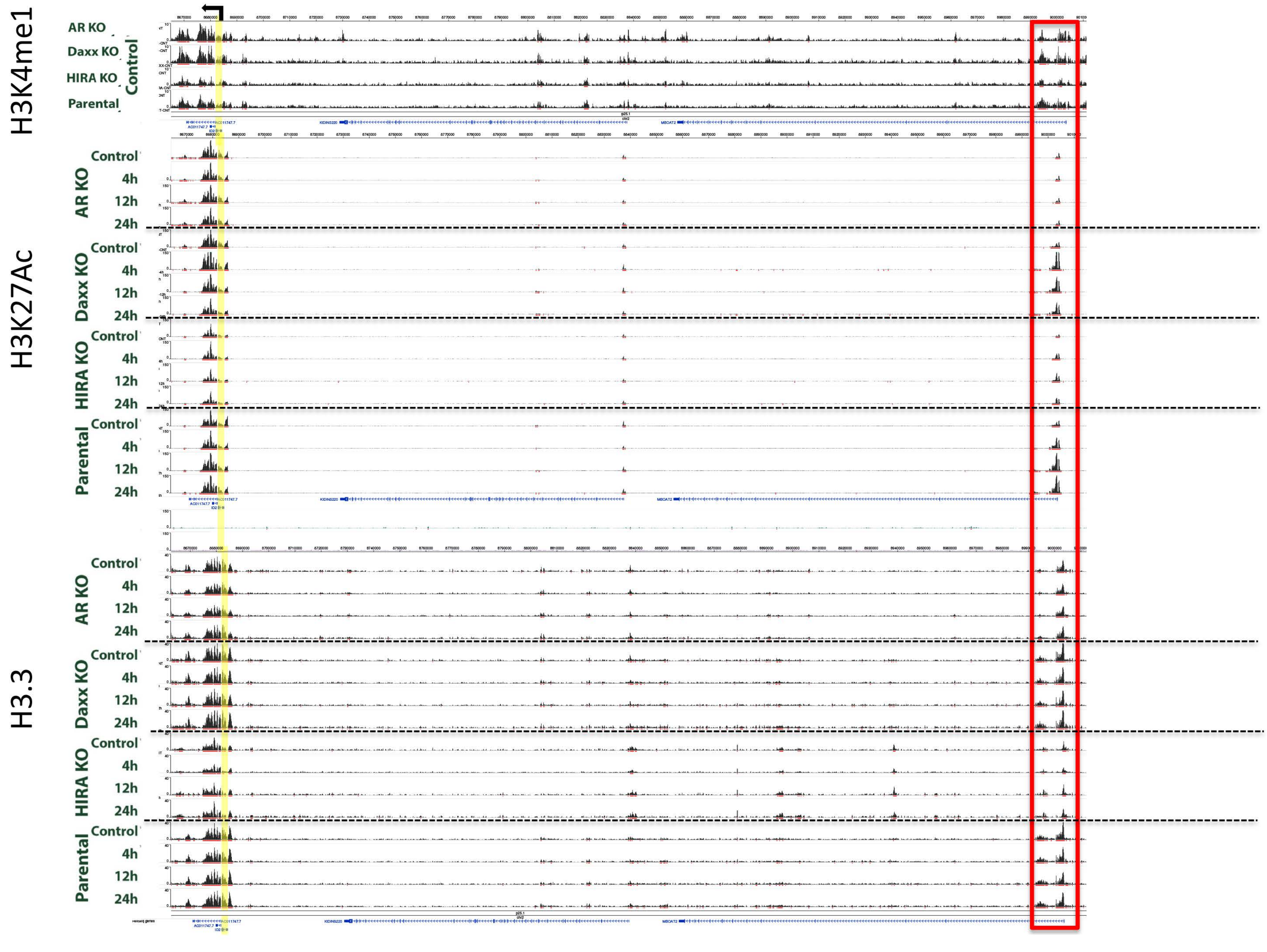

### Fig. S14B

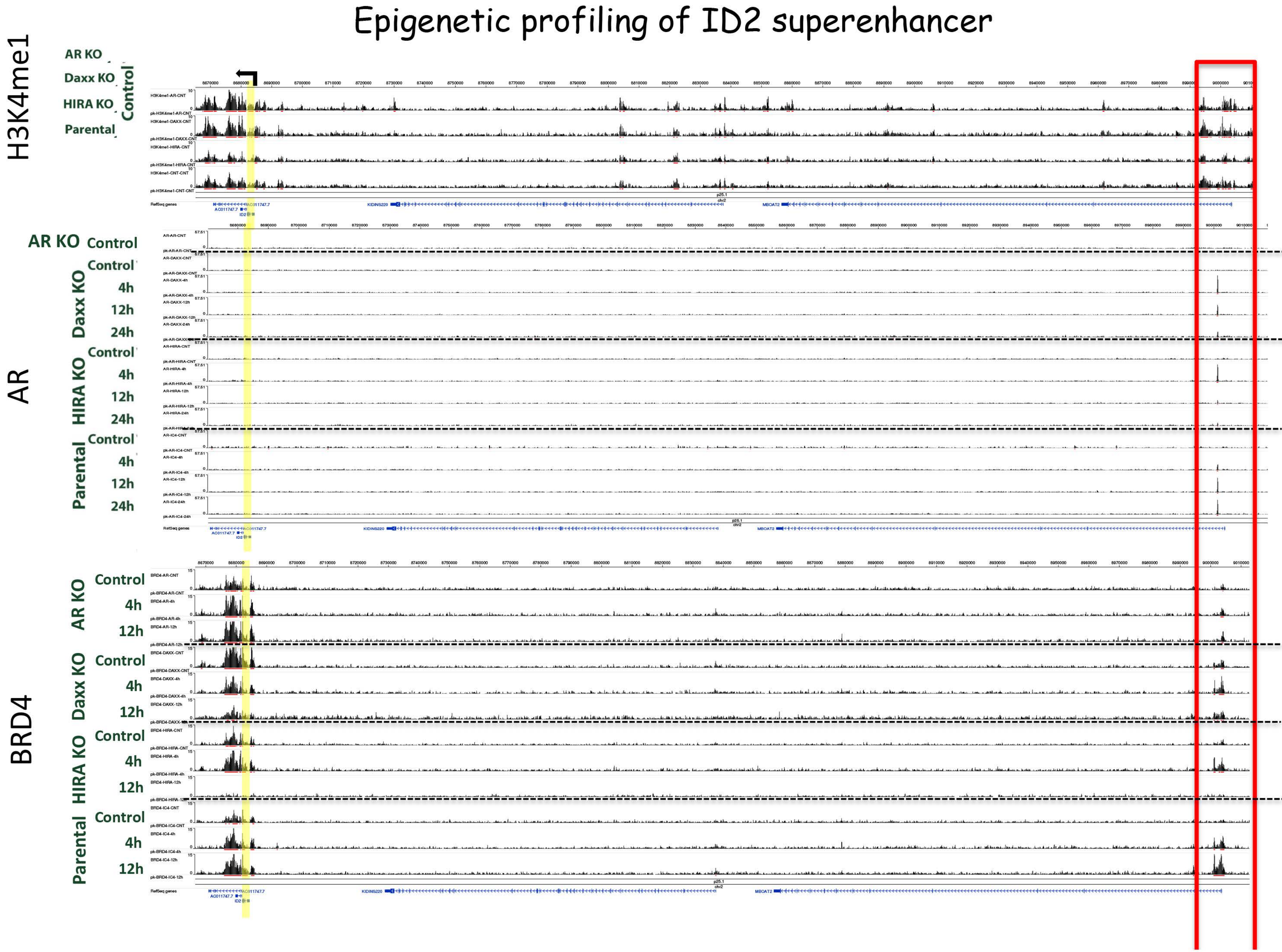
Epigenetic profiling of ID2 superenhancer
H3K4me1
AR KO
Daxx KO
HIRA KO
Parental
Control
AR
AR KO Control
Daxx KO
Control
4h
12h
24h
HIRA KO
Control
4h
12h
24h
Parental
Control
4h
12h
24h
BRD4
AR KO
Control
4h
12h
Daxx KO
Control
4h
12h
HIRA KO
Control
4h
12h
Parental
Control
4h
12h

### Fig. S14B

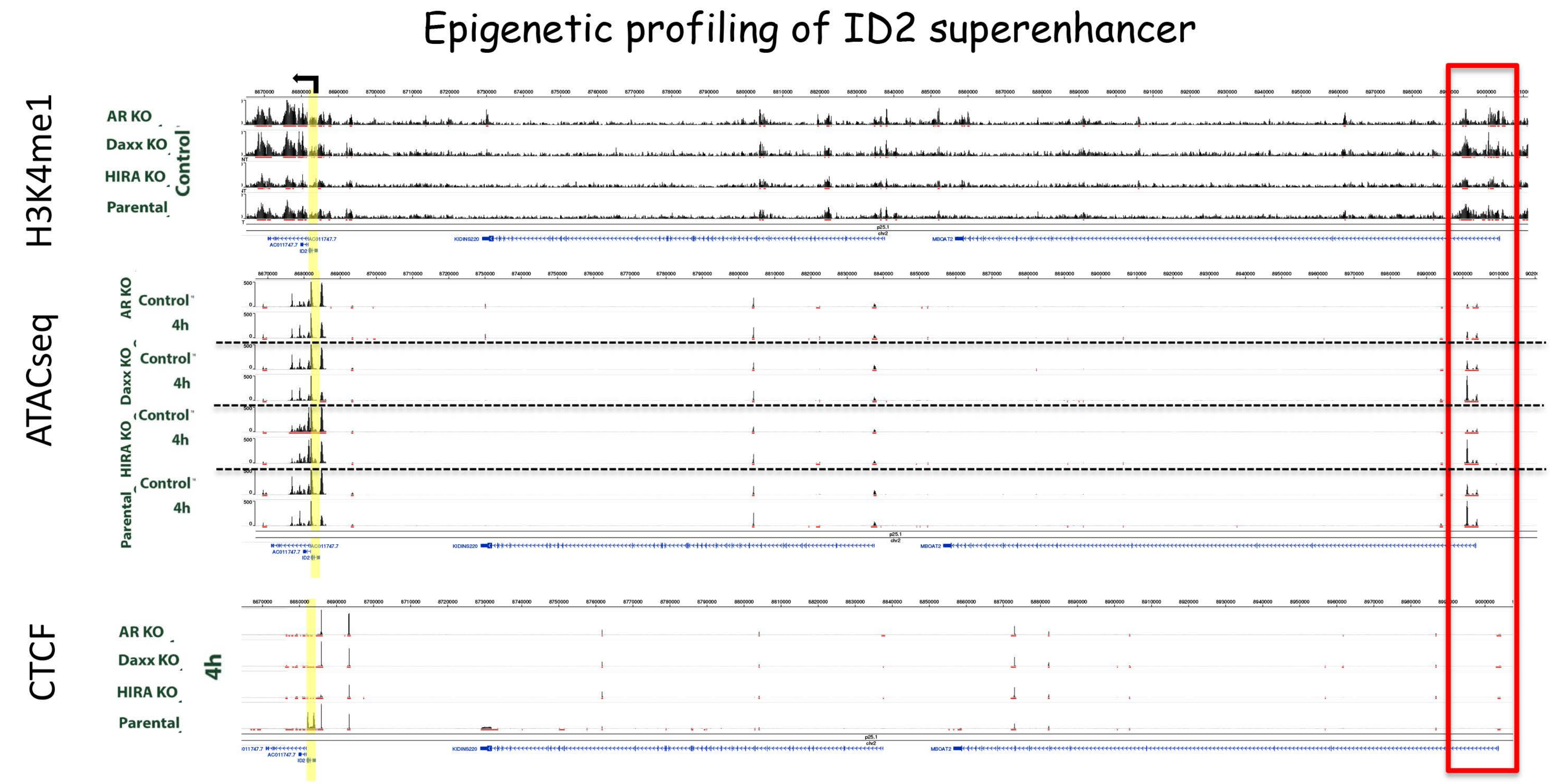
Epigenetic profiling of ID2 superenhancer
H3K4me1
AR KO
Daxx KO
HIRA KO
Parental
Control
ATACseq
AR KO
Control
4h
Daxx KO
Control
4h
HIRA KO
Control
4h
Parental
Control
4h
CTCF
AR KO
Daxx KO
HIRA KO
Parental
4h
